# Adapting Nanopore Sequencing Basecalling Models for Modification Detection via Incremental Learning and Anomaly Detection

**DOI:** 10.1101/2023.12.19.572431

**Authors:** Ziyuan Wang, Yinshan Fang, Ziyang Liu, Ning Hao, Hao Helen Zhang, Xiaoxiao Sun, Jianwen Que, Hongxu Ding

## Abstract

We leverage machine learning approaches to adapt nanopore sequencing basecallers for nucleotide modification detection. We first apply the incremental learning technique to improve the basecalling of modification-rich sequences, which are usually of high biological interests. With sequence backbones resolved, we further run anomaly detection on individual nucleotides to determine their modification status. By this means, our pipeline promises the single-molecule, single-nucleotide and sequence context-free detection of modifications. We benchmark the pipeline using control oligos, further apply it in the basecalling of densely-modified yeast tRNAs and *E.coli* genomic DNAs, the cross-species detection of N6-methyladenosine (m6A) in mammalian mRNAs, and the simultaneous detection of N1-methyladenosine (m1A) and m6A in human mRNAs. Our IL-AD workflow is available at: https://github.com/wangziyuan66/IL-AD.

## INTRODUCTION

The nanopore sequencing technology translates biomolecule chemical structures into ionic current signals, therefore opens up opportunities for routinely detecting DNA and RNA modifications^1^. State-of-the-art modification detection algorithms, e.g. EpiNano^2^, differr^3^, DRUMMER^4^, nanoRMS^5^, ELIGOS^6^ and Dinopore^7^, determine “error signatures” for modification detection. The rationale is that chemical modifications deviate nanopore sequencing signals further disrupting basecalling and alignment. The assumption is that yielded bioinformatic errors represent informative signatures that encode modifications. The dilemma is that, besides generating “error signatures”, shifted signals could disrupt basecalling and alignment completely. As a result, a large amount of reads, in particular those densely-modified thus of high biological interests, will never be analyzed. Another group of methods, e.g. tombo^8^, signalAlign^9^, nanopolish^10–12^, DeepMod^13^, DeepSignal^14^, MINES^15^, nanoDoc^16^, nanom6A^17^, Nanocompore^18^, Yanocomp^19^, xPore^20^, Penguin^21^ and m6Anet^22^, determine modification status of nucleotides from corresponding sequencing signals. As prerequisites of the workflow, basecalling and alignment results are used to correspond signal chunks with sequence kmers. These methods are therefore also less capable of analyzing modification-rich sequences.

Densely modified loci, however, deliver profound biological insights. For example, most methylated cytosines cluster within short genomic regions of high C+G frequency (CpG islands)^23^. It is widely-acknowledged that the CpG island methylation dynamics controls diverse biological processes, e.g. carcinogenesis^24^ and development^25^. Meanwhile, up to 20% tRNA nucleotides can be modified. Such modifications stabilize tRNA structures, therefore crucial for the protein translation^26^. Besides, artificial modification hotspots are commonly introduced as biological probes. For example, BrdU is supplied to substitute thymidine during cell cycle, therefore can be used to track nascent DNA chunks further pinpointing replication origins^27^; the exogenous GC-specific methyltransferase is used to methylate accessible genomic sequences, further labeling open chromatin regions^12^.

To better analyze such modification-rich sequences, we exploit incremental learning (IL) techniques to upgrade existing basecallers. IL extends existing deep learning models to address both old and new tasks^28^. In our case, IL will generalize basecallers to resolve sequence backbones for both canonical (old) and modified (new) nanopore sequencing readouts. IL-basecallers will therefore provide sequence backbones for each individual molecule, on top of which modifications could be analyzed.

This basecalling-based modification detection paradigm has three unique advantages. First of all, basecalling, together with the subsequent alignment, could polish sequence backbones against references. Such polished sequences could facilitate more precise modification detections, especially considering the relatively high error rate of nanopore sequencing compared to next generation sequencing platforms. Meanwhile, basecalling determines sequence basebones for individual sequencing readouts, therefore enabling the single-molecule, single-nucleotide modification detection. In addition, state-of-the-art basecallers, e.g. guppy and dorado from Oxford Nanopore Technologies, are sequence context-free. By this means, molecules of virtually all kinds, ranging from genomes and transcriptomes to artificially synthesized oligos, could be accurately basecalled. Based on such context-free sequence backbones, modifications residing in any motifs could be resolved. Such a capability will greatly facilitate, e.g. the discovery of novel modification motifs involved in epigenetic^29^ and epitranscriptomic regulations^30^, and understandings towards mutagenesis by randomly incorporated tobacco-chemicals^31^ and oxyradicals^32^.

Based on sequence backbones determined from basecalling, we next leverage anomaly detection (AD) techniques to scrutinize modification status of individual nucleotides. AD summarizes a group of statistical approaches for identifying significantly deviated data observations^33^, in our case modification-induced signals. Our two-step IL-AD workflow is summarized in Figure 1A.

**Figure 1.**
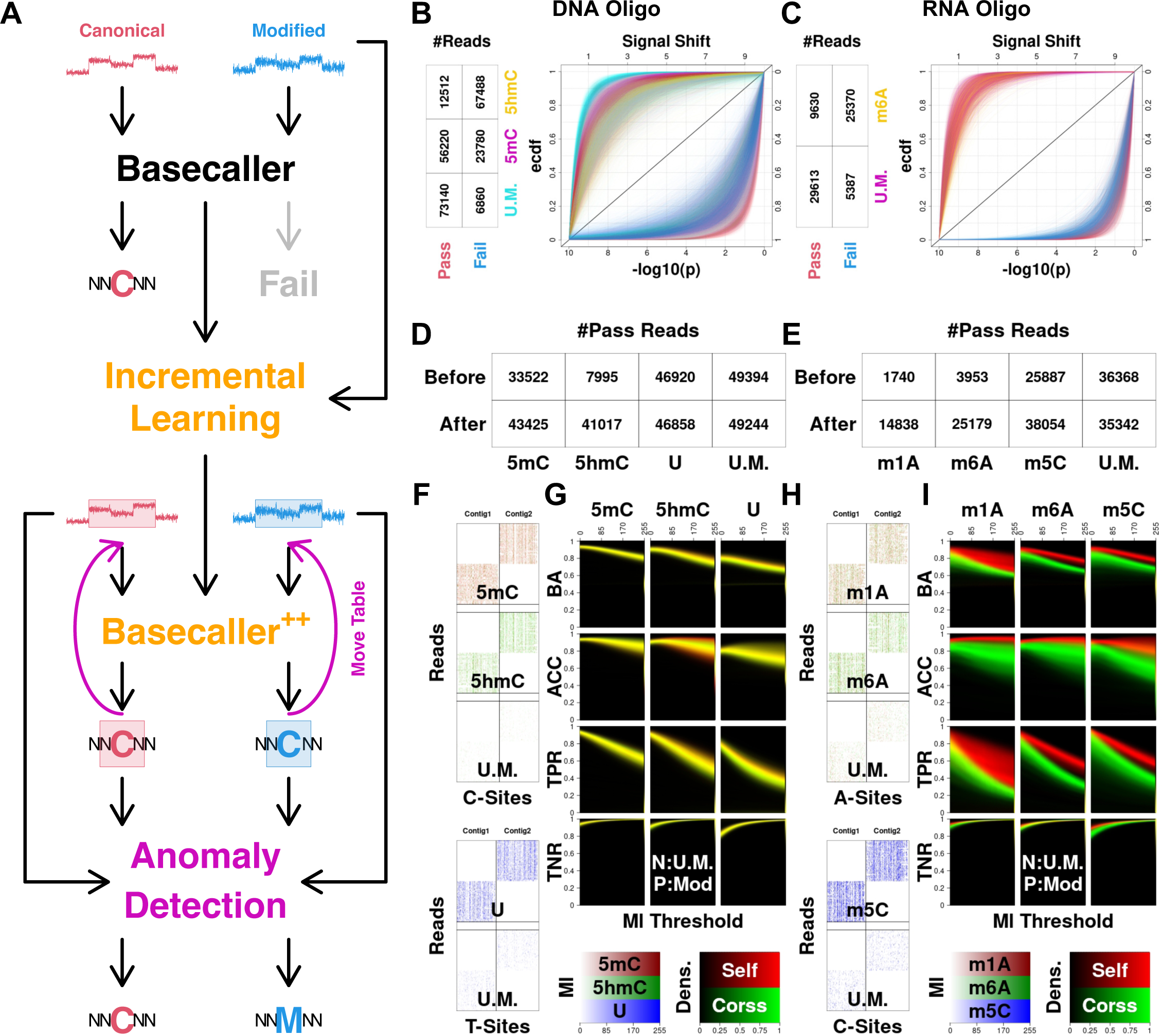
Adapt nanopore sequencing basecallers for modification detection. (A) Workflow overview. The move table records signal-nucleotide correspondences. C and M denote canonical and modified nucleotides, respectively. (B, C) Fully modified oligos cause basecalling failures, which can be explained by significantly shifted signals. Pass and fail denote sequencing reads that can and cannot be basecalled, respectively. For each kmer instance, its signal shift was quantified against the canonical kmer model by a p-value. P-values yielded from the same kmer were concluded with an ecdf (empirical cumulative distribution function) curve. U.M. denotes unmodified sequencing reads. (D, E) Ameliorate basecalling failures with the incremental learning adaptation. Before and after denote pre and post-adaptation, respectively. (F, H) Per-read-per-site visualization of Ml-tags. Ml-tags represent modification scores. Without losing generality, 50 randomly selected full-length reads from contig 1 & 2 were visualized. (G, I) Quantify modification detection performance using confusion matrix statistics, including Balanced Accuracy (BA), Accuracy (ACC), True Positive Rate (TPR) and True Negative Rate (TNR). These statistics were calculated for each modification site at different Ml-tag thresholds, and visualized using density heatmaps. Positive and negative classes denote modified and unmodified nucleotides, respectively. Self and cross denote performance quantification on the same and the additional independent oligo ensembles, respectively.

## RESULTS

### Dense modifications disrupt nanopore sequencing basecalling and alignment

We noticed decreased mappabilities for modification-rich molecules, e.g. fully-modified RNA oligos^2,7^ and native tRNAs^26^. To confirm such observations, we designed, produced and sequenced our own control oligos. We then created ground-truth labels for these oligos via an iterative approach (see METHODS and Figure S1). Without losing generality, we surveyed mappabilities of 5-methylcytosine (5mC) and 5-hydroxymethylcytosine (5hmC) fully-modified DNA oligos, N6-methyladenosine (m6A) fully-modified RNA oligos, as well as their unmodified counterparts. As shown in Figure 1B & C, modified oligos are more likely to fail the basecalling and alignment process, resulting in decreased mappabilities. We further observed that unmappable failures are caused by more deviated nanopore sequencing readouts, by quantifying signal shifts against canonical kmer models (see METHODS). We reasoned that state-of-the-art basecalling and alignment pipelines are only able to process “canonical-like” signals. Those deviated “characteristic modification signals”, which are in general of high biological interests, are more likely to be ignored.

### Basecall modification-rich sequences via incremental learning (IL) adaptation

We ameliorated the above-described inability of processing modification-rich sequences via IL. Specifically, we fine-tuned existing basecallers with iteratively-labeled, fully-modified oligos. Such training data combines 5mC, 5hmC and U-oligos for DNA, and m1A, m6A and 5-methylcytosine (m5C)-oligos for RNA. We evaluated the effectiveness of IL using independent oligo datasets (see METHODS). As shown in Figure 1D & E, IL remarkably increases mappabilities of oligos that are less likely to be analyzed previously, e.g. DNA 5hmC, and RNA m1A and m6A. We further found that IL ameliorates “error signatures” by increasing basecalling accuracy (Figure S2). Importantly, we noticed that decreases in mappabilities (Figure 1D & E) and basecalling accuracy (Figure S2) within unmodified oligos are negligible. Taken together, we concluded the success of IL in the decoding of modification-rich signals, without the catastrophic forgetting^34^ of “old tasks” in decoding unmodified nanopore sequencing readouts.

### Determine nucleotide modification status via anomaly detection (AD)

With IL, we were able to resolve sequence backbones for both densely-modified and canonical-like signals. Along each resolved backbones, we then predicted modification status of every nucleotides, by examining their corresponding signal chunks using AD. Specifically, we trained one AD model per modification isoform, together with a baseline unmodified AD model. For a specific site to be tested, we inputted the corresponding signal chunk into all associated AD models, and calculated loss ratios between baseline and modification models as Ml-tag modification scores (see METHODS). For instance, when analyzing a DNA C-site, we will 1) train 5mC, 5hmC and C models, 2) calculate loss values against three models and 3) assess C/5mC and C/5hmC loss ratios as 5mC and 5hmC Ml-tags, respectively. Following such a rationale, we assessed Ml-tags for various DNA and RNA oligos. We visualized Ml-tags in selected reads (50 random full-length reads for contig 1 and 2), to demonstrate the single-molecule-single-nucleotide detection resolution of our approach (Figure 1F & H). We then quantified modification prediction performance with confusion matrices generated at various Ml-tag thresholds. Specifically, we surveyed all modification sites, and concluded results using density heatmaps (“Self” in Figure 1G & I). We noticed high and comparable prediction accuracy across sites, e.g. >0.8 average balanced accuracy with <0.02 standard deviation at Ml-threshold 5, for all oligo groups. We therefore highlighted the consistency of our modification detection analysis. We also noticed performance differences between modification groups, e.g. overall accuracy for U is lower then 5mC and 5hmC; cross-site consistency for m1A is lower then m6A and m5C. We speculated modification-specific signal patterns and dwell times to be major causes of such differences.

### Generalize existing modification detection models to new sequence contexts

We further examined whether trained modification detection models could be generalized to previously unseen sequence contexts. Specifically, we generated nanopore sequencing data from an independent oligo ensemble (see METHODS). We then executed trained modification detection models on such new datasets, and calculated confusion matrices as in the above section. We marked this scheme as “Cross”, and observed comparable performance compared to “Self” in DNA analyses. We therefore highlighted an accurate and sequence context-free modification detection in DNAs. Regarding RNA scenarios, however, we observed a moderate performance decrease in “Cross” groups, e.g. ∼0.1 average balanced accuracy at Ml-threshold 5 (Figure 1G & I). We further demonstrated that “Self-Cross” discrepancies can be ameliorated with shorter signal chunks, however at the cost of less accurate predictions (Figure S3). We argued that while longer signal chunks contain more information in judging nucleotide modification status, broader up and downstream contexts will also be represented. Such sequence contexts are largely different between the two oligo ensembles, thus resulting in “Self-Cross” discrepancies. We reasoned that such discrepancies are more noticeable in RNA due to dwell times.

### Basecall densely-modified biological sequences with IL

We exploited IL to address real-world biological challenges. As a proof-of-concept, we first basecalled native yeast tRNAs, which are densely-modified thus extremely difficult to be basecalled. As shown in Figure 2A & B, bioinformatic analyses with recommended mapping parameters^26^ yield low mappability and prevalent “error signatures”. We then ran IL on the RNA basecaller with the iteratively-labeled tRNA training data (see METHODS). As shown in Figure 2A, IL drastically increases mappabilities, e.g. a 50-fold increase for Ala-AGC, of virtually all tRNA species. Meanwhile, we noticed that IL could maintain tRNA relative abundance, which suggested an unbiased basecalling. We further noticed that IL delivers accurate sequence backbones by clearing “error signatures” (Figure 2B). We also ruled out the catastrophic forgetting by correctly interpreting unmodified DNA oligos (Figure S4A).

**Figure 2.**
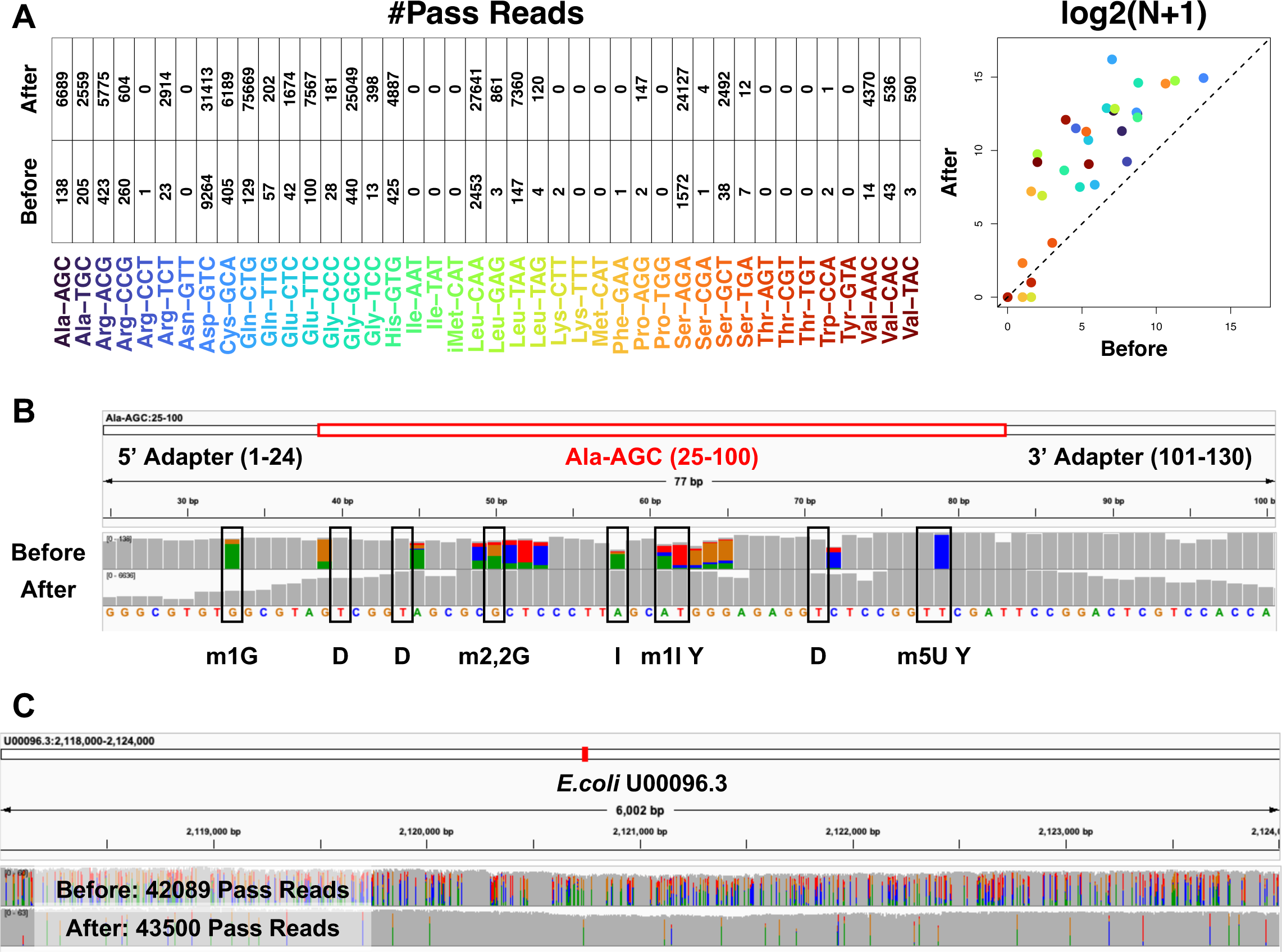
Basecall densely-modified sequences to facilitate real-world biological studies. (A) Mappabilities of native yeast tRNAs. (B) “Error signatures” of native yeast tRNAs. Without losing generality, we visualized the tRNA Ala-AGC as an example. (C) Mappabilities and “error signatures” of CpG and GpC methylated *E.coli* genomic DNAs. Without losing generality, we visualized a 6 kb segment as an example. Throughout this figure, before and after denote pre and post-IL, respectively.

We next basecalled artificially-created, CpG and GpC-methylated *E.coli* genomic DNAs. Such reads were used to train machine learning models for the simultaneous profiling of CpG and GpC methylation, which denote epigenetic status and chromatin accessibility, respectively^12^. Albeit the decent mappability that has already been achieved by regular bioinformatics, IL could still make improvements. Besides, IL is able to clear most “error signatures” which are common in conventional bioinformatics (Figure 2C). We also ruled out the catastrophic forgetting by correctly basecalling unmodified *E.coli* genomic DNAs (Figure S4B). We therefore concluded the success of IL in basecalling densely-modified DNAs and RNAs in real-world biological scenarios.

### Detect mRNA m6A sites across mammalian species with AD

We further examined whether AD is capable of detecting modifications in real-world biological scenarios. As a proof-of-concept, we first detected mRNA m6A sites in the human HEK293 cell line. We trained AD models combining native mRNA nanopore sequencing data^35^ and m6A site annotations determined using m6ACE-Seq^36^. We executed the model on a biologically independent HEK293 native mRNA nanopore sequencing dataset, then quantified the detection performance using confusion matrix statistics (see METHODS). As shown in Figure 3A, we achieved high prediction accuracy for m6A sites, e.g. balanced accuracy reaches ∼0.99 at Ml-threshold 5.

**Figure 3.**
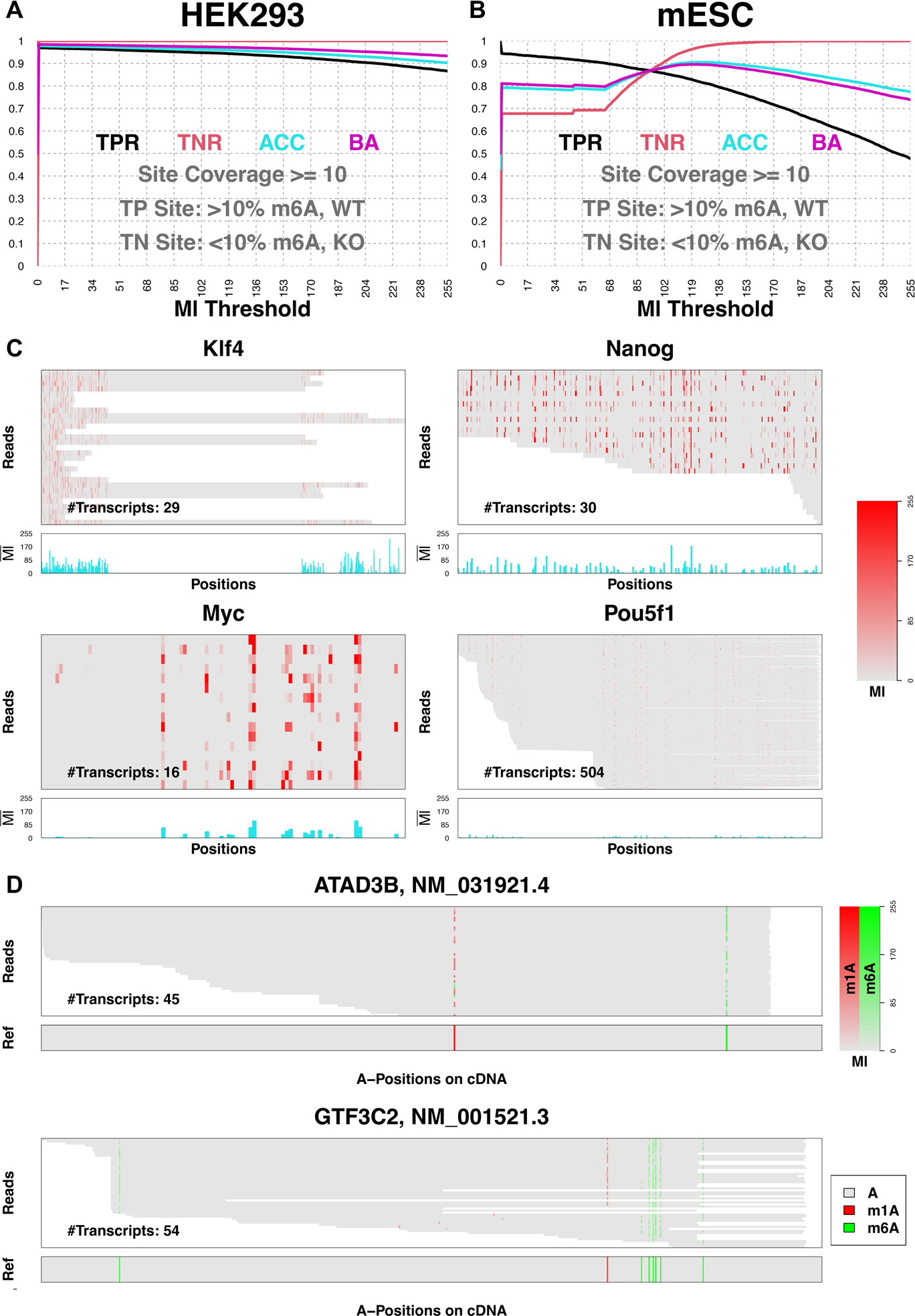
Detect mRNA modifications to facilitate real-world biological studies. (A, B) Quantifying mRNA m6A site detection accuracy in human HEK293 cells and mouse embryonic stem cells (mESCs) with confusion matrix statistics. TPR, True Positive Rate; TNR, True Negative Rate; ACC, Accuracy; BA, Balanced Accuracy. Such statistics were quantified for each modification site under different Ml-tag modification score thresholds. Only sites covered by ≥10 mRNAs were included to ensure statistical rigor. True positive and true negative sites were defined as having >10% m6A in the wild type (WT) sample and <10% m6A in the knock-out (KO) sample, respectively. (C) Surveying mESC mRNA m6A profiles of Yamanaka factors (*Klf4*, *Nanog*, *Myc*, *Pou5f1*). Per-read-per-site Ml-tags and per-site average Ml-tags were visualized with heatmaps and barplots, respectively. (D) Simultaneously detecting m1A and m6A in HEK293 *ATAD3B* and *GTF3C2* mRNAs. Per-read-per-site Ml-tags were visualized using heatmaps. Known m1A and m6A sites were marked in “Ref” lower-panels.

We next assessed the cross-species generalizability of the above human AD model. We thus applied the model to a mouse embryonic stem cell (mESC) native mRNA nanopore sequencing dataset^6^. We set m6A sites determined using m6A-CLIP^37^ as ground-truth, and quantified prediction performance with confusion matrix statistics (see METHODS). As shown in Figure 3B, we observed only a gentle performance decrease under such a cross-species scenario: the maximum balanced accuracy could still reach ∼0.9. We thus concluded the success in generalization, although species-specificity cannot be ignored.

To further confirm the cross-species generalizability, we surveyed the m6A landscape of Yamanaka factor transcripts. It has been demonstrated that 3’ UTRs of *Klf4*, *Nanog* and *Myc*, but not *Pou5f1*, are densely methylated with m6A^38^. Consistently, analysis with the human model revealed a similar m6A pattern in last exons of mouse Yamanaka factors (Figure 3C). Taken together, these results indicate the cross-species generalizability of the human m6A AD model.

### Detect m1A and m6A simultaneously in individual human mRNAs with AD

Finally we leveraged AD to explore a novel research challenge of simultaneously profiling m1A and m6A in the epitranscriptome. Besides the above HEK293 m6ACE-Seq annotations, we included ground-truth m1A sites determined by m1A-Seq^39^ to train AD models. We executed the model on the same HEK293 biological replicate as above-mentioned for the per-read-per-site and simultaneous detection of m1A and m6A (see METHODS). As a proof-of-concept, we reported the recapitulation of m1A and m6A sites in *ATAD3B* and *GTF3C2* transcripts, as shown in Figure 3D.

## DISCUSSION

Detecting any nucleotide modifications under any sequence motifs remains a prominent challenge in the nanopore sequencing field. Our IL-AD framework is uniquely suitable to solve such a challenge by achieving 1) high detection accuracy for diverse modifications and 2) generalizability towards new sequence contexts. With these potentials, our IL-AD provided new insights in real-world biological questions, e.g. analyzing densely-modified tRNAs and genomic DNAs, generalizing the mRNA m6A detection across species, and simultaneously profiling m1A and m6A epitranscriptomic markers. Albeit IL-AD to be an appropriate framework to solve the modification detection challenge, several limitations remain to be solved.

### Nanopore sequencing signal pattern and dwell time affect modification prediction performance

We speculated signal patterns and dwell time to be the two major factors affecting modification prediction performance in our current workflow design. First of all, if signal patterns between modified and canonical nucleotides are similar, then AD could become challenging. One example for such scenarios is the U-oligo analysis. As shown in Figure 1D, substituting T with U in DNAs will not significantly compromise basecalling and alignment. Such a result implies similar patterns between U and canonical-signals. Meanwhile, TPR and TNR for U-predictions were systematically and significantly lower as opposed to 5mC and 5hmC (Figure 1G), suggesting confusions when distinguishing U with T. We therefore concluded signal pattern differences as one major source of the modification-specificity. We further expect upgrades in nanopore sequencing hardware and experimental kits for more distinguishable signal patterns among modifications as potential solutions for such an issue.

Dwell time quantifies the biomolecule translocation speed during nanopore sequencing, and has been demonstrated as an intrinsic characteristic of modifications^18^. Meanwhile, because of different sequencer setups, dwell times for DNA and RNA are also different. More importantly, dwell time is largely affected by the translocation speed stochasticity. Since the current AD design takes fixed-length signal chunks, corresponding “sequence context scopes” could vary largely, as shown in Figure S5. Such varying scopes further compromise AD, by making 1) intra and inter-read signal chunks, as well as 2) baseline and modification models not comparable. Such artifacts together introduce performance variations, which further explain 1) the m1A-specific less consistent performance, and 2) the RNA-specific “Self-Cross” discrepancy. We speculated that taking a fixed number of consecutive signal segments, rather than a fixed number of signal points, during AD will be a potential solution for dwell time-related artifacts. We previously demonstrated that signal segments are yielded from sequence kmers (k equals 6 and 5 for DNA and RNA, respectively)^40^. Therefore, fixing the number of signal segments could largely ameliorate context scope variations. The signal segmentation, which is known as event detection in nanopore sequencing, can be addressed by change-point detection methods^41^.

### Full-motif AD models for modification detection under any sequence motifs

We asked whether the AD model trained using m6A-oligos can be generalized for the m6A detection in human and mouse native mRNAs. The accomplishment of this task implies oligo-based AD models to be universal for detecting modifications under any sequence motifs. However, we observed compromised generalizability of the m6A-oligo AD model in detecting m6As in mammalian mRNAs. We speculated the cause to be, besides the “Self-Cross” discrepancy as discussed above, that certain sequence motifs, e.g. A(m6A) were not represented in fully-modified oligos. To address this problem, further delivering universal modification detection models, we propose the production of full-motif training oligos by randomly incorporating modifications to be one potential solution.

### The production of modified oligos

We noticed that incorporating modifications into oligos could be challenging. We used PCR and *in vitro* transcription (IVT) to produce modified DNA and RNA oligos, respectively (see METHODS). We observed that modifications tend to terminate PCR and IVT prematurely, resulting in a large scale of truncated readouts. We also observed that certain modifications, such as DNA 6mA, are extremely challenging to be incorporated with our current experimental setups. We speculated that modifications could disrupt basepairing, which might further abort PCR and IVT. To address this problem, further deliver an universal experimental pipeline for producing training oligos of any kind, we believe novel biochemical assays are needed.

## METHODS

### Design and Synthesize DNA and RNA Oligos

We designed oligo sequences following procedures reported in ^2^ using the CURLCAKE software (http://cb.csail.mit.edu/cb/curlcake/). Specifically, we covered all possible DNA 6mers (4,096 in total, median occurrence as 5) and RNA 5mers (1,024 in total, median occurrence as 10). We also avoided secondary structures, the formation of which during nanopore sequencing will increase the translocation speed further biasing ionic current signals^42, 43^. We generated two independent sequence sets, for both DNA and RNA, with different CURLCAKE random seeds. Such sets represent different sequence contexts: as larger sequence scopes (k > 6 and 5 for DNA and RNA, respectively) were surveyed, the fraction of overlapping kmers against total kmers among sets drastically decreased. Lengths for yielded DNA and RNA sequences are 25 kb and 12.5 kb, respectively. We splitted CURLCAKE sequences into “curlcakes” of ∼2.5 kb for synthesis purposes. For each DNA curlcake, we added HindIII sites to both 3’ and 5’ ends, and removed all the internal HindIII sites. For each RNA curlcake, we first added a strong T7 promoter to the 5’ end. We then added EcoRV sites to both 3’ and 5’ ends, and removed all the internal EcoRV and BamHI sites. Final DNA and RNA sequence backbones were provided in Table S1 and S2, respectively. We synthesized and cloned all the DNA and RNA curlcakes into the pUC57 vector using blunt EcoRV and HindIII, through the service of GenScript Biotech Corporation.

For DNA curlcakes, we incorporated modified nucleotides using PCR. dNTP mixtures, including unmodified (dATP, dTTP, dCTP, dGTP), 5mC (dATP, dTTP, 5m-dCTP, dGTP), 5hmC (dATP, dTTP, 5-hme-dCTP, dGTP) and U (dATP, dUTP, dCTP, dGTP) were mixed with engineered pUC57 plasmids, primers, the reaction buffer and polymerase for PCR. Yielded products were analyzed by agarose gel electrophoresis and extracted using the gel extraction kit. The concentration of resulting DNA was determined by the NanoDrop 2000 Spectrophotometer.

For RNA curlcakes, we incorporated modified nucleotides using *in vitro* transcription (IVT). Engineered pUC57 plasmids were digested with EcoRV and BamHI restriction enzymes for at least two hours at 37 °C, and analyzed via agarose gel electrophoresis. Purification of the digested DNA was conducted using a PCR purification kit. Nanodrop was used to measure the concentration of extracted DNA prior to IVT. Ampliscribe^TM^ T7-Flash^TM^ Transcription Kit was used to generate IVT RNAs as per manufacturer’s instructions. During IVT, modified ribonucleoside triphosphates including N1-Methyl-ATP (m1A), N6-Methyl-ATP (m6A), 5-Methyl-CTP (m5C) and Pseudouridine-5-Triphosphate (Psi) were supplemented in place of their unmodified counterparts. DNAse I was added to the IVT reaction system after incubation for 4 hours at 42 °C to eliminate the residual template DNA. Yielded IVT RNAs were purified using the RNeasy Mini Kit following manufacturer’s instructions. NEB vaccinia capping enzyme was used for the 5’ capping of purified IVT RNAs, with an incubation for 30 min at 37 °C. Following purification with RNAClean XP Beads, the capped IVT RNAs were subjected to polyadenylation tailing. Concentration of capped and polyA-tailed IVT RNAs was determined by Qubit Fluorometric Quantitation.

### Nanopore Sequencing

DNA nanopore sequencing libraries were prepared using the ONT Ligation Sequencing Kit (SQK-LSK110) following protocol version ACDE_9110_v110_revN_10Nov2020 as per manufacturer’s instructions. Briefly, for each group (unmodified, 5mC, 5hmC and U), 100 fmol of PCR DNA was subjected to repair and end-prep with NEBNext PPFE DNA Repair Mix and NEBNext Ultra II End repair/dA-tailing Module kits, respectively. After purification using AMPure XP Beads, the product was subjected to adapter ligation with NEBNext Quick Ligation Module, as the DNA sequencing library. The Qubit fluorometer was used to determine the concentration of the DNA library. The DNA library was mixed with the sequencing buffer and loading beads prior to sequencing on a primed MinION flow cell. The flow cell version is R9.4.1, and the sequencer is MinION.

RNA nanopore sequencing libraries were built using the ONT Direct RNA Sequencing Kit (SQK-RNA002) following protocol version DRS_9080_v2_revQ_14Aug2019 as per manufacturer’s instructions. Briefly, for each group (unmodified, 5mC, 5hmC and Psi), 2 μg of capped and polyA-tailed IVT RNA was subjected to adapter ligation using the NEB T4 DNA Ligase, following reverse transcription using the SuperScript III Reverse Transcriptase. After purification using RNAClean XP Beads, yielded RNA:DNA hybrids were ligated to RNA adapters using the NEB T4 DNA Ligase. The concentration of the yielded RNA library was determined by the Qubit fluorometer. The RNA library was mixed with RNA Running Buffer prior to sequencing on a primed Flongle flow cell. The flow cell version is R9.4.1, and the sequencer is MinION with a Flongle adapter.

### Iterative Label Modification-Rich Sequences

Based on the observation that guppy is able to basecall a fraction of fully-modified DNA and RNA oligos, we reasoned that chemical moiety changes on nucleotides are, in most cases, less likely to drastically shift nanopore sequencing signal distributions. Therefore, modification signals with smaller shifts can still be correctly basecalled. Such basecalled signals will further be used to train guppy, and the yielded model will by this means gain more information for basecalling modification signals. Such a process will be iterated till convergence to label modification signals, as shown in SFigure 1.

For the DNA oligo labeling, guppy version 6.0.6+8a98bbc was used for basecalling and alignment. We used the template_r9.4.1_450bps_hac.jsn model for the first iteration, and the model trained from the previous iteration for the subsequent iteration. The --disable_qscore_filtering flag was used to keep “low-quality reads” that are usually artifacts caused by modifications. Samtools version 1.16 was used to merge, sort and index alignment results outputted by guppy. Taiyaki version 5.3.0 was used for guppy training. For preparing training data, get_refs_from_sam.py with default flags, generate_per_read_params.py with default flags, prepare_mapped_reads.py with the mLstm_flipflop_model_r941_DNA.checkpoint, merge_mappedsignalfiles.py with default flags were used. For training guppy models, train_flipflop.py with default flags and the template mLstm_cat_mod_flipflop.py, as well as dump_json.py with default flags on the final model checkpoint were used. We performed in total 3 iterations.

The RNA labeling followed the process except: 1) template_rna_r9.4.1_70bps_hac.jsn was used for the initial basecalling, 2) the --reverse flag of get_refs_from_sam.py was used, 3) the r941_rna_minion.checkpoint was used during prepare_mapped_reads.py, 4) flags --size 256 --stride 10 --winlen 31 were used for train_flipflop.py.

The iterative labeling of yeast native tRNAs followed the same process as RNA oligos, with the exception that -ax map-ont -k5 -w5 flags were used for the minimap2^44^ (version 2.24-r1122) alignment, which was recommended in^26^.

The iterative labeling of *E.coli* CpG and GpC-methylated genomic DNAs followed the same process as DNA oligos.

### Quantify Nanopore Sequencing Signal Shifts

We used ONT kmer models as references to measure signal shifts. During sequencing, consecutive nucleotide kmers translocate through nanopores, further producing signal events. Events aligned to the same kmer are in general summarized using a Gaussian distribution. The mean and standard decoration of Gaussian, together with other trivial parameters for all possible kmers are recorded in kmer models. Specifically, k equals 6 and 5 for DNA and RNA, respectively^40^.

For modified oligos, we first used iterative labeling to generate accurate basecalling and alignment profiles. Such profiles were subsequently used for nanopolish eventalign^10–12^ (version 0.13.3) with the flag --scale-events to make event tables. Event tables record event parameters, including signal levels and aligned kmers for all sequencing events. For events aligned to the same kmer, we calculated p-values against the corresponding Gaussian distribution in the kmer model. We summarized p-values with the empirical cumulative distribution function to measure the per-kmer signal shifts. We randomly selected ∼80,000 sequencing reads for each DNA oligo group (unmodified, 5mC, 5hmC), and ∼35,000 sequencing reads for each RNA oligo group (unmodified, m6A) for signal shift quantification analyses.

### Incremental Learning

We adopted knowledge distillation^45^ for the IL-adaptation of DNA and RNA basecallers. Specifically, we freezed original basecallers throughout the entire IL process as teacher models, and initialized tunable student models by duplicating teacher models. During IL, training data for new tasks (basecall modification-induced signals) flowed through both teacher and student models. Our goals are 1) making sure student models are capable of accomplishing new tasks precisely, and 2) controlling for the catastrophic forgetting of old tasks (general basecalling) by forcing student models to produce similar outputs with teacher models. We therefore introduced Connectionist Temporal Classification (CTC)^46^ and Response-Based Knowledge Distillation (RBKD)^47^ loss terms to reach such goals, respectively. We further balanced such contradicting terms as the final optimization goal for the basecaller IL-adaptation.

We denoted **X** as a signal chunk and **Y** as the corresponding nucleotide sequence. The student model transforms **X** as CTC matrix **U**, where 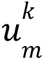 indicates the **U** value at position *k* ∈ [1, *K*] and alphabet *m* ∈ [1, *M*]. We summarized all paths traversing **U** that can be decoded as **Y** into the valid CTC path set **C**. We further wrote the CTC loss as:

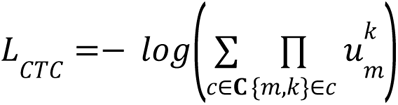

We further denoted the counterpart of **U** in the teacher model as **V**, and wrote the RBKD loss as:

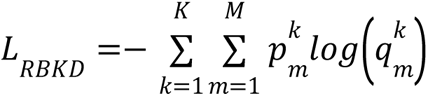

where 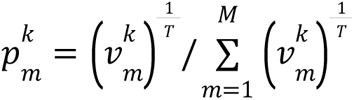 and 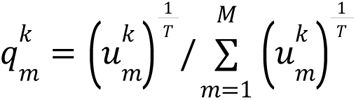, and *T* is the temperature parameter used to scale probability ratios.

We balanced *L*_*CTC*_ and *L*_*RBKD*_ using hyperparameter λ, and wrote the final loss as:

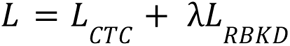

With such an optimization goal, we fine-tuned ONT taiyaki DNA and RNA basecallers, with training data prepared through iterative labeling, and using the AdamW optimizer^48^. For all IL analyses, we set λ, learning rate and epoch as 10, 1e-5 and 1e3, respectively.

### Anomaly Detection

We further leveraged AD to distinguish between canonical and modification-induced signals. Specifically, we trained one CTC network (template mLstm_flipflop.py for DNA, template mGru_flipflop.py for RNA) per nucleotide isoform, e.g. C, 5mC, 5hmC for DNA cytosine, as its AD model. During AD model training, we first used the “ref_to_signal” entry in the hdf5 file (in which taiyaki stores prepared training data) to retrieve the first signal point that corresponds to the candidate nucleotide for AD. We then took the upstream *n* and downstream *n* + *m* signal points, based on which further retrieved the underlying sequence. We minimized the signal-sequence CTC loss using the AdamW optimizer^48^. We set learning rate and epoch as 5e-5 and 5e3 for all AD analyses, respectively, except for the HEK293 m1A AD model, whose epoch was set as 2e3 for a better detection performance. We set *n* and *m* as 10 and 20 for DNA, respectively, and 45 and 60 for RNA, respectively.

When executing AD models to detect the modification status of a candidate nucleotide, we first retrieved the 2*n* + *m* signal chunk as above-mentioned as the model input, then calculated the modification likelihood as:

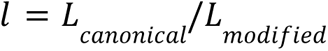

where *Lcanonical* and *Lmodification* denote CTC loss of the canonical and modification AD models, respectively. We further converted the *l* value as an Ml-tag with following rules:

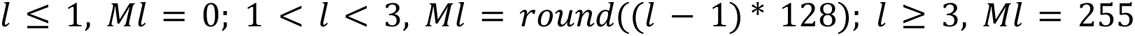

and wrote the result into the sam/bam file with pysam^49^.

### Analyze Synthesized Oligos with IL-AD

For the DNA scenario, we first trained a general IL-basecaller by combining 5mC, 5hmC and U oligos. We randomly sampled ∼80,000 sequencing reads for each oligo group as training data, and made ground-truth labels via iterative labeling. With the same training data, we further trained C, 5mC and 5hmC AD models to analyze C-modification status in unmodified, 5mC and 5hmC oligos, and T and U AD models to analyze T-modification status in unmodified and U oligos. We then assessed the trained IL-AD workflow with an independent ensemble of unmodified, 5mC, 5hmC and U oligos. We randomly sampled ∼50,000 sequencing reads and performed iterative labeling as test data. During the test phase, we first performed IL-basecalling and alignment to determine mappabilities and “error signatures”. For mappable reads, we further performed AD to determine Ml-tags.

For the analysis of RNA unmodified, m1A, m6A and 5mC oligos, we followed the same process as in DNA, except ∼100,000 sequencing reads per oligo group were used for IL-AD training.

### Basecall Yeast Native tRNAs and Artificially-Methylated *E.coli* Genomic DNAs

We trained the RNA IL-basecaller with native yeast tRNA nanopore sequencing dataset 7_NanotRNAseq_WTyeast_rep1. Specifically, we determined sequence backbones with iterative labeling, based on which then performed IL. To ensure a balanced training data representation, we randomly sampled 500 sequencing reads per tRNA type. For tRNA species with fewer than 500 reads, we used all available reads. With the trained model, we basecalled an independent biological replicate 11_NanotRNAseq_WTyeast_rep2. We then performed the minimap2^44^ (version 2.24-r1122) alignment with -ax map-ont -k5 -w5 flags following ^26^.

For the CpG and GpC-methylated *E.coli* genomic DNA analysis, we randomly sampled ∼80,000 sequencing reads for iterative labeling, based on which then performed IL. We randomly sampled an independent set of ∼80,000 sequencing reads as test data.

### Detect m6A in Human and Mouse mRNA Transcripts

AD models for m6A detection were trained using human HEK293 cell line datasets. We first surveyed m6ACE-Seq profiles and identified a total of 15,073 *METTL3*-dependent ground-truth m6A sites^36^. Focused on these sites, we subsequently trained A and m6A AD models using native mRNA nanopore sequencing profiles of wild type and *METTL3* knock-out samples^35^, respectively. During model training, we first used guppy version 6.0.6+8a98bbc and the model template_rna_r9.4.1_70bps_hac.jsn for basecalling. We next performed the minimap2^44^ (version 2.24-r1122) alignment using -ax splice -uf k14 flags against the GRCh38 reference transcriptome. We then extended the taiyaki script get_refs_from_sam.py to pinpoint known m6A sites among spliced mRNA reads, further preparing training data hdf5 files for both wild type (HEK293T-WT-rep1) and knock-out (HEK293T-Mettl3-KO..1) samples using prepare_mapped_reads.py paired with the r941_rna_minion.checkpoint. With wild type and knock-out hdf5 files, we trained m6A and A AD models, respectively.

We evaluated such models on a biologically independent sample pair (wild type sample HEK293T-WT-rep2 and knock-out sample HEK293T-Mettl3-KO..2). We next performed evaluation on mouse embryonic stem cells (mESCs), using native mRNA nanopore sequencing profiles from ^6^, and retrieving a total of 30,519 *METTL3*-dependent m6A sites from ^37^.

### Simultaneously Detect m1A and m6A in Human mRNA Transcripts

We followed the above-described AD model training pipeline in this section. Specifically, we used the same HEK293 nanopore sequencing collections and m6A annotations. We considered sites revealed in ^39^ as m1A ground-truth. Without losing generality, we took *ATAD3B* transcript isoform NM_031921.4 and *GTF3C2* transcript isoform NM_001521.3 as examples to demonstrate the recapitulation of m1A-m6A co-occurrence.

### Data Availability

The yeast native tRNA nanopore sequencing data was downloaded from European National Archive (ENA) under accession number PRJEB55684. Corresponding reference genome and modification annotation were downloaded from https://github.com/novoalab/Nano-tRNAseq/tree/main/ref. The CpG and GpC methylated, and the unmodified *E.coli* genomic DNA nanopore sequencing datasets were downloaded from https://sra-pub-src-2.s3.amazonaws.com/SRR11953238/ecoli_CpGGpC.fast5.tgz.2 and https://sra-pub-src-2.s3.amazonaws.com/SRR11953241/ecoli_Unmethylated.fast5.tgz.1, respectively. Corresponding reference genome was downloaded from https://www.ncbi.nlm.nih.gov/nuccore/U00096. The human HEK293 cell line native mRNA nanopore sequencing data was downloaded from ENA under accession number PRJEB40872. Corresponding m1A and m6A ground-truth annotations were downloaded from the Supplementary Table 2 of ^39^ and the Supplementary Data 4 of ^36^, respectively. The mouse ESC native mRNA nanopore sequencing data was downloaded from NCBI Sequence Read Archive (SRA) under the accession number SRP166020. Corresponding m6A ground-truth annotations were downloaded from NCBI Gene Expression Omnibus (GEO) under the accession number GSM2300431. DNA and RNA oligo datasets were deposited at NCBI under the BioProject PRJNA1050579 (reviewer link: https://dataview.ncbi.nlm.nih.gov/object/PRJNA1050579?reviewer=p904dqm4m3879vgr9jjof4gg0). ONT DNA and RNA kmer models were downloaded from https://github.com/nanoporetech/kmer_models. Original DNA and RNA basecalling models were downloaded from https://github.com/nanoporetech/taiyaki/blob/master/models/mLstm_flipflop_model_r941_DNA.checkpoint and https://s3-eu-west-1.amazonaws.com/ont-research/taiyaki_modbase.tar.gz, respectively. DNA and RNA taiyaki model templates were downloaded from https://github.com/nanoporetech/taiyaki/tree/master/models.

### Code Availability

The IL-AD workflow is available at: https://github.com/wangziyuan66/IL-AD.

## ACKNOWLEDGEMENTS

We thank the University of Arizona High Performance Computing team and the College of Pharmacy Information Technology Group for their support. H.D. is supported by the University of Arizona Health Sciences Career Development Award. J.Q. is supported by HL159675, HL152293, AI163753 and DK132251.

## AUTHOR CONTRIBUTIONS

Z.W. and H.D. conceived the idea. Y.F. performed the experiment. Z.W., Z.L. and H.D. performed the analysis. N.H., H.H.Z., X.S., J.Q. and H.D. supervised the project. Z.W., Y.F. and H.D. wrote the manuscript.

