## Supplementary for "Adapting Nanopore Sequencing Basecalling Models for Modification Detection via Incremental Learning and Anomaly Detection"

#### Supplementary Information

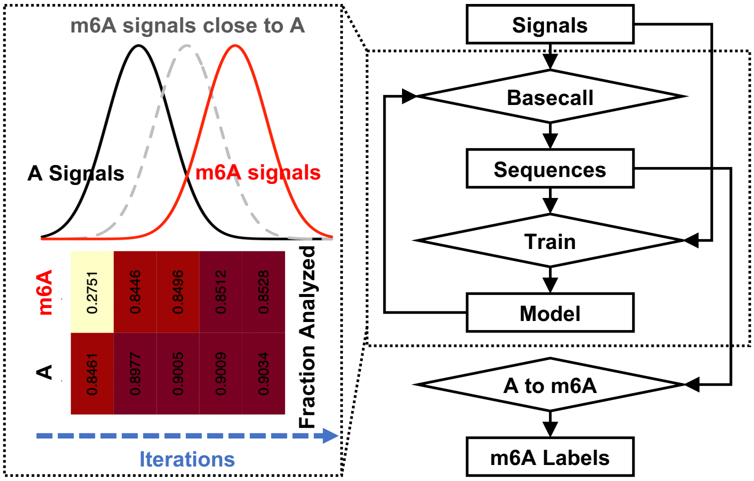
m6A signals close to A
A Signals
m6A signals
m6A
A
0.2751
0.8446
0.8496
0.8512
0.8528
0.8461
0.8977
0.9005
0.9009
0.9034
Fraction Analyzed
Iterations
Signals
Basecall
Sequences
Train
Model
A to m6A
m6A Labels

**Figure S1. The iterative labeling workflow.**

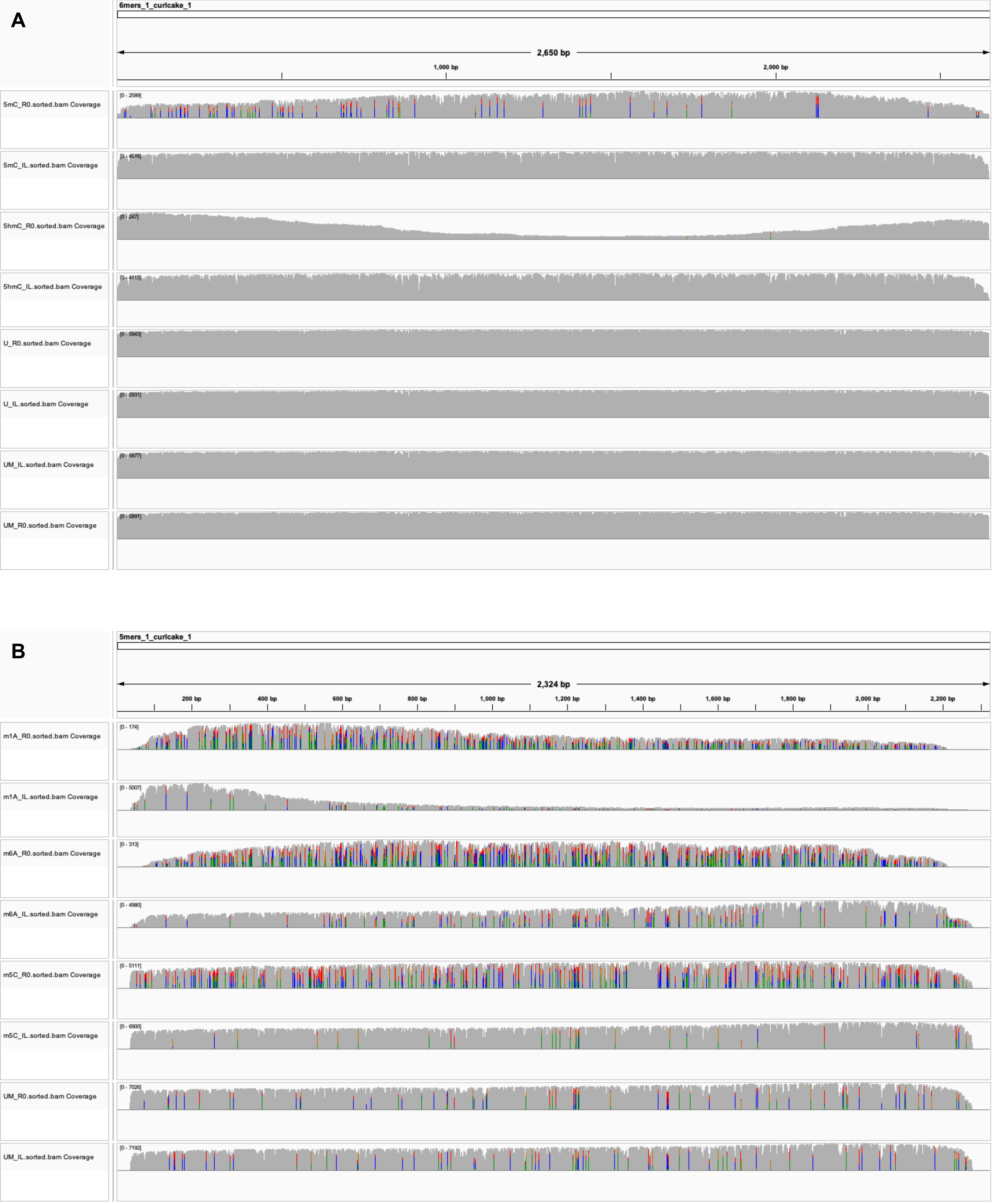

A
6mers_1_curlcake_1
2,650 bp
1,000 bp
2,000 bp
5mC_R0.sorted.bam Coverage
[0 - 2589]
5mC_IL.sorted.bam Coverage
[0 - 4516]
5hmC_R0.sorted.bam Coverage
[0 - 247]
5hmC_IL.sorted.bam Coverage
[0 - 4113]
U_R0.sorted.bam Coverage
[0 - 5943]
U_IL.sorted.bam Coverage
[0 - 5931]
UM_IL.sorted.bam Coverage
[0 - 5877]
UM_R0.sorted.bam Coverage
[0 - 5891]
B
5mers_1_curlcake_1
2,324 bp
200 bp
400 bp
600 bp
800 bp
1,000 bp
1,200 bp
1,400 bp
1,600 bp
1,800 bp
2,000 bp
2,200 bp
m1A_R0.sorted.bam Coverage
[0 - 174]
m1A_IL.sorted.bam Coverage
[0 - 5007]
m6A_R0.sorted.bam Coverage
[0 - 313]
m6A_IL.sorted.bam Coverage
[0 - 4980]
m5C_R0.sorted.bam Coverage
[0 - 5111]
m5C_IL.sorted.bam Coverage
[0 - 6900]
UM_R0.sorted.bam Coverage
[0 - 7026]
UM_IL.sorted.bam Coverage
[0 - 7192]

**Figure S2. Effects of IL on basecalling accuracy.** IGV plots of DNA (5mC, 5hmC, U, U.M.) and RNA (m1A, m6A, m5C, U.M.) oligos were shown in (A) and (B), respectively. R0 and IL denote pre and post-IL, respectively. Without losing generality, only the DNA contig 6mers_1_curcake_1 and the RNA contig 5mers_1_curcake_1 were visualized.

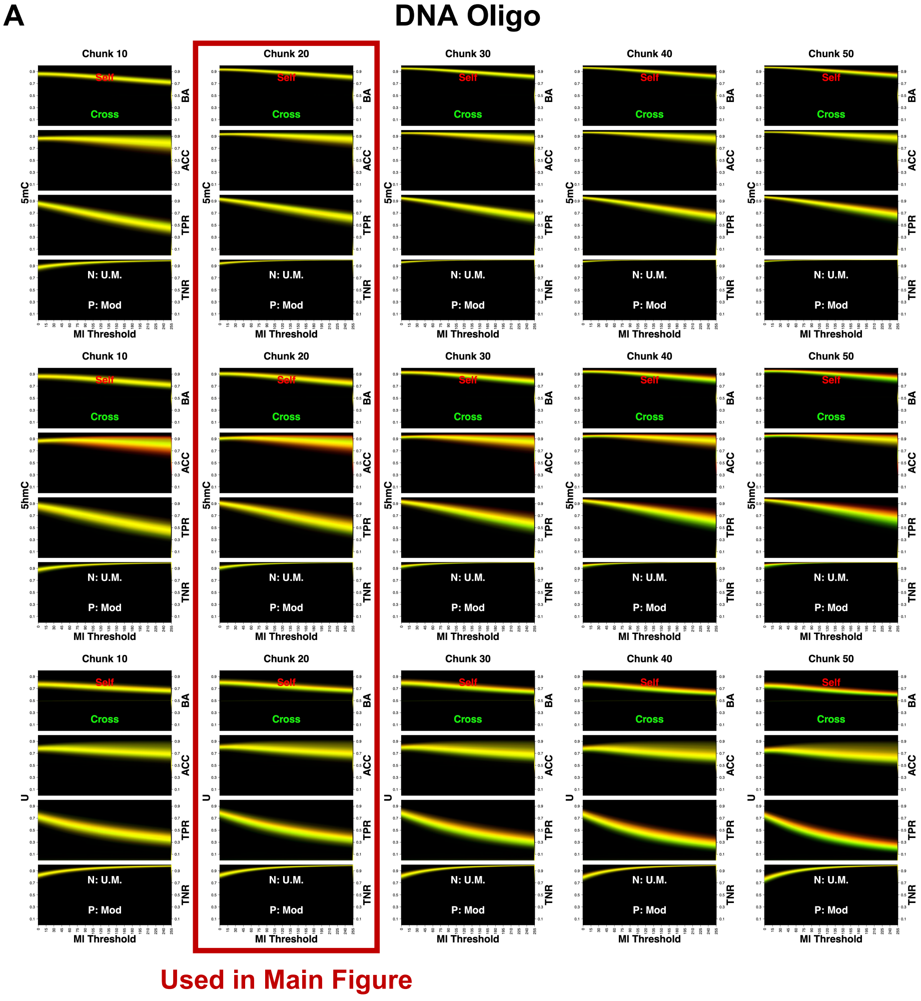
A
DNA Oligo
Chunk 10
Chunk 20
Chunk 30
Chunk 40
Chunk 50
Self
Cross
BA
ACC
TPR
TNR
N: U.M.
P: Mod
MI Threshold
5mC
5hmC
U
Used in Main Figure

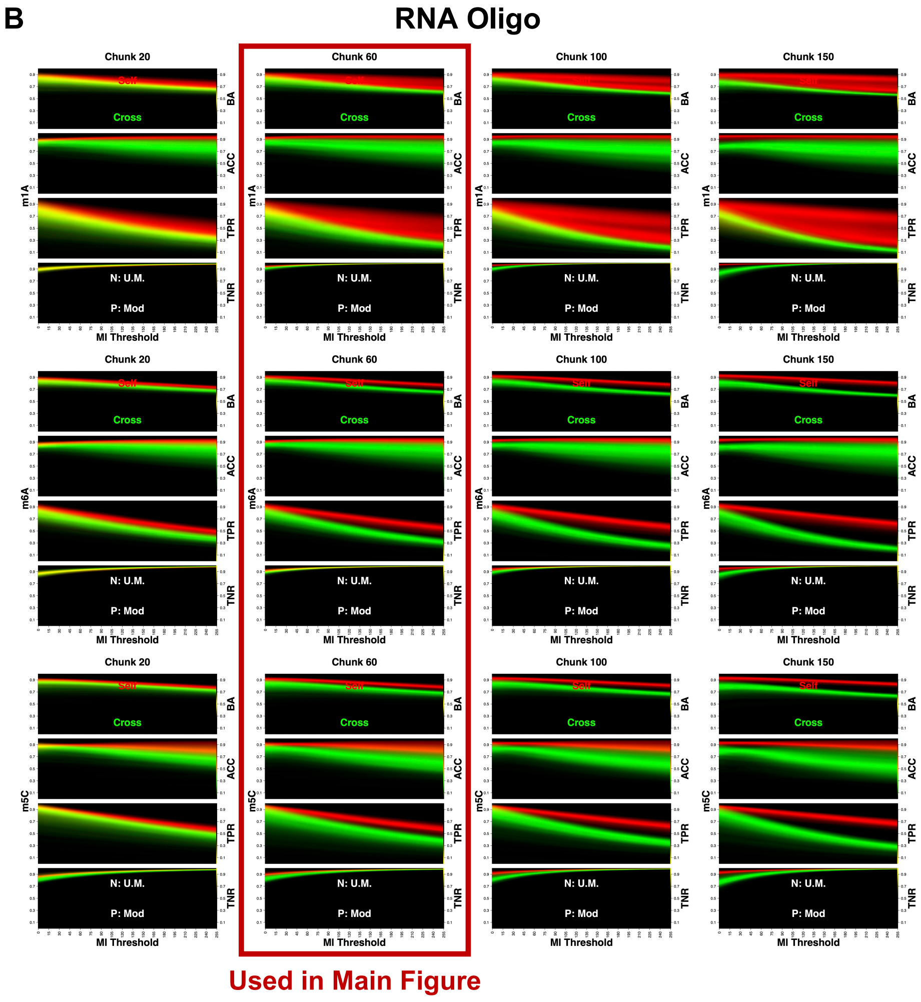
B
RNA Oligo
Chunk 20
Chunk 60
Chunk 100
Chunk 150
Self
Cross
BA
ACC
TPR
TNR
N: U.M.
P: Mod
MI Threshold
m1A
m6A
m5C
Used in Main Figure

**Figure S3. Effects of signal chunk size on AD performance.** AD performance was quantified by confusion matrix statistics, including Balanced Accuracy (BA), Accuracy (ACC), True Positive Rate (TPR) and True Negative Rate (TNR). These statistics were calculated for each modification site at different Ml-tag thresholds, and visualized using density heatmaps. DNA and RNA heatmaps were visualized in (A) and (B), respectively. Self and Cross denote executing the AD model trained with set1 oligos on set1 and set2 oligos, respectively. Positive and negative classes denote modified and unmodified nucleotides, respectively.

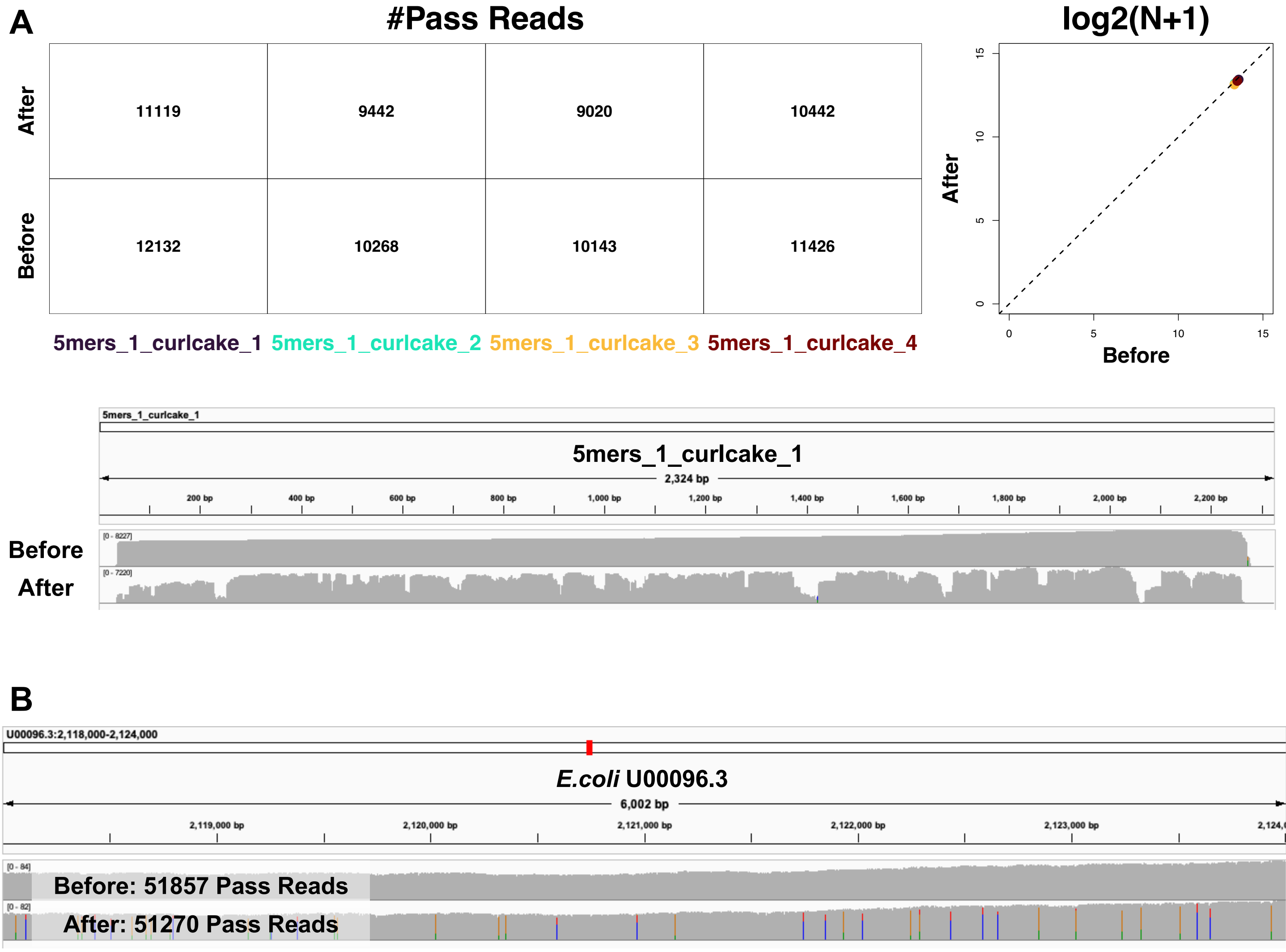
A
#Pass Reads
After
Before
11119
9442
9020
10442
12132
10268
10143
11426
5mers_1_curlcake_1 5mers_1_curlcake_2 5mers_1_curlcake_3 5mers_1_curlcake_4
log2(N+1)
After
Before
0
5
10
15
5mers_1_curlcake_1
2,324 bp
200 bp
400 bp
600 bp
800 bp
1,000 bp
1,200 bp
1,400 bp
1,600 bp
1,800 bp
2,000 bp
2,200 bp
[0 - 8227]
[0 - 7220]
Before
After
B
U00096.3:2,118,000-2,124,000
E.coli U00096.3
6,002 bp
2,119,000 bp
2,120,000 bp
2,121,000 bp
2,122,000 bp
2,123,000 bp
[0 - 84]
[0 - 82]
Before: 51857 Pass Reads
After: 51270 Pass Reads

**Figure S4. Assessing potential catastrophic forgetting effects resulted from IL.** (A) Mappabilities and basecalling accuracy when applying the yeast native tRNA-tuned IL basecaller on unmodified RNA oligos. Without losing generality, we selected the contig 5mers_1_curlcale_1 for IGV visualization. (B) Mappabilities and basecalling accuracy when applying the *E.coli* CpG and GpC methylated genomic DNA-tuned IL basecaller on the unmodified *E.coli* genome. Without losing generality, we selected a 6 kb segment for IGV visualization.

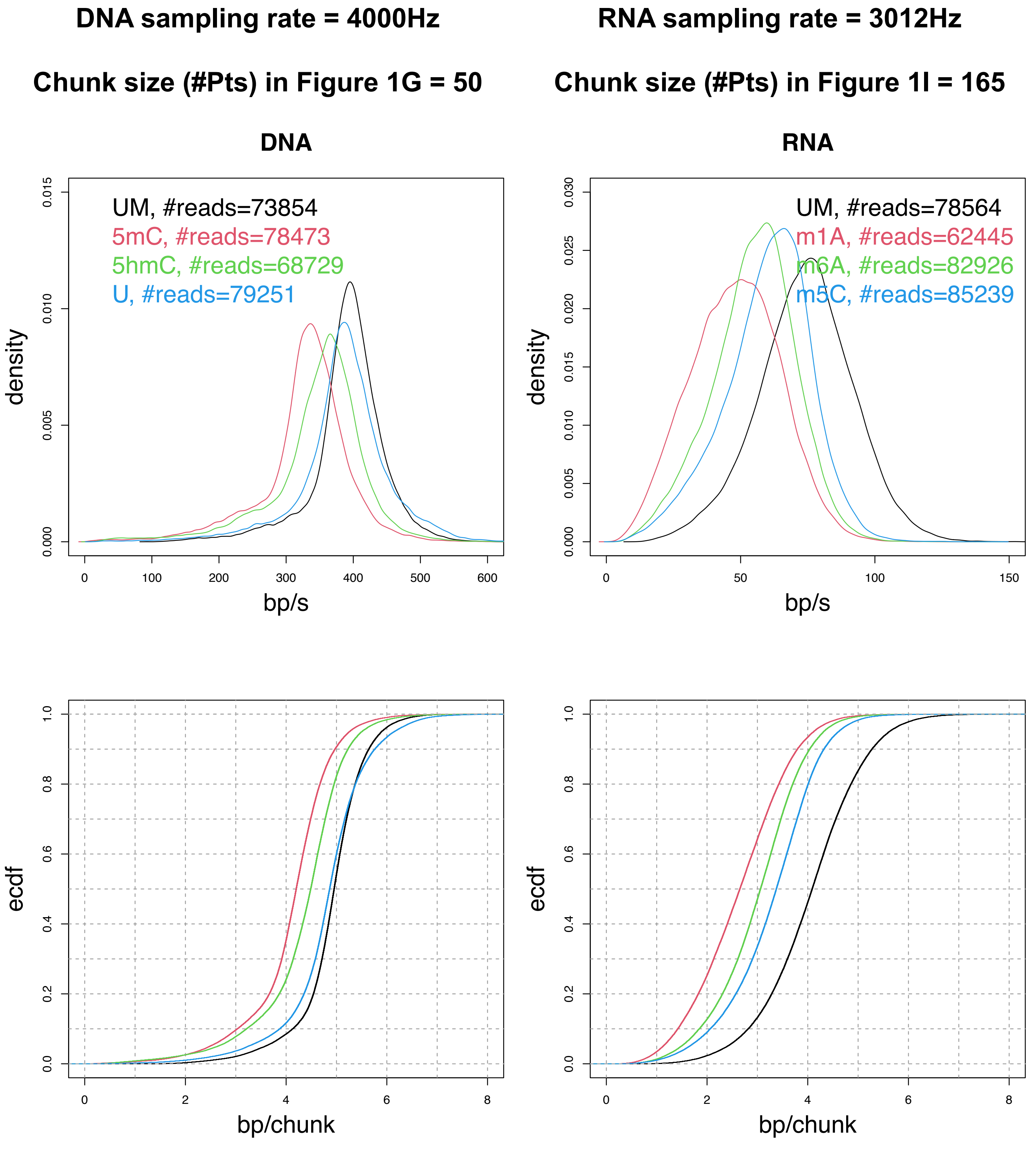
DNA sampling rate = 4000Hz
Chunk size (#Pts) in Figure 1G = 50
RNA sampling rate = 3012Hz
Chunk size (#Pts) in Figure 1I = 165
DNA
UM, #reads=73854
5mC, #reads=78473
5hmC, #reads=68729
U, #reads=79251
density
bp/s
RNA
UM, #reads=78564
m1A, #reads=62445
m6A, #reads=82926
m5C, #reads=85239
density
bp/s
ecdf
bp/chunk
ecdf
bp/chunk

**Figure S5. Effects of dwell time variations on “sequence context scopes”.**

### Table S1. Sequence backbones of synthesized DNA oligos.

>6mers_1_curlcake_1
AAAAACCCGGTTGTACGACATCATGTCATGTATTTCTCAAAGCATTCGCCTCGCCTTTTACGGATCAACCAAAGGAAAGGGCAGCCGCACACCTTACACTACACTGTCAAGGTACCAGAGCCCACCCCACTTTGAACCTCACATCCCGAGCCGAGCTATGCCGCGACAGT
TTTCATTTCACGTGAACGCCCATTGGTCCAGTCCAATGGTAAGCAAATGCAGCTAAGCTAATACGACCTGGATCACCTGTCCTGTAACAGCTTCTGCTTAAACCGAACCGGGAGTTTACAACAGGGGCGAGGCGACTGATCCCAACGAGCTACCTTACCTCGTCCTGCTA
TCCGAGTCAGGTCAGTCGTTCAAGTTAGAAACTAGACTAGATGTCCCGCTCTTACCTGAGCTGAGCCGACGAGCTGCGCTACTGTACTGTCGCTGACAACGAAACCATCTCATCTACCGACCTCGACCGCAACTGAACTGATATGGCTCGTGGCACATTGCATTGCTGCA
GAAGACGGGCCCGTGCCGTGCGGACTGCTTTCGACCATACCATACTCATCCCTGGCTACTCACGACACGAGAATTAGTTTTAGAGGCAGAGCAGACAGTAGGATGAAATCGAGGCGAGGCCATCGCTATAGCAACCGCTTCGCTTGTGTAACGCTCGAACGGGACGGG
ACGTGCCATTCAGTTCAGCGATGCGATTCTAATTGTATGTCATCACTTCACTGTTGGCGCCCCTGCAACATAACATACGGCCCATTACCTACGCCTCGCCTCCGTCTGCACTATACGTCGTCTCGTCCGCGACAAATGAGTTCTGCACTGCATAAAGATCAGTATTTTCTTCT
CTTCGGTCGCGTGTAGGGAAAGCTAGCTTAGAGCAGGGGATCTTACATTACATCATGCATTCGGCGAAAAAGAAAAGACAAGACTCGATACCAACCCCACCCCGAGTAAGTTTATACCGCGAAGCGAACTTTGTGGCTGGGGCACCCTACCCTGGAGATATGTTTGCTGG
CATGGCATAATCGTTCCTACCCAGTGGGGTGGGACATGTGTCGGGGATGGTAAGGTAAACAACGCGTCAGATTCCATGCCATGAATGTGCCATACTTAACTCCACTCCTCTGTGCGATTAGTGATCCAATCCAATTGACTCCACGGGTAAGTCAAGTCTCCCAACTATCCT
CCTTTCTTTTCTGTAGGGCATGCTCGGGTGCGGTGCGGGGGCTATTATTACAGTCGCGTCGCAAGCTACTGCTATGAACAAAACAAACCTGTGTGGCTCGAGTCTTCTCTTCCGTTTATCAACTCCCGGTGCGGTGCCCGCGTTTGTTACTTTTTCATTTCATTATGATTGA
ATTAAGCAGTGCAGTCCATAAGCGCCCTAGTGACCTGACCCGTGACACATAGGCACTGAACTGAATTTCGCCCGCATGTTTCCTCTCCTCACCCGTGGTTGTTACTATTCTATTCGCCAAACTCCAGTGCCAAGACAAGACGTCACGAGGGGGATACGGAACGGAAATCC
TAGCTATCCCGATATGATATCGATCCTGAACACAGTTCCGTTCCGGCATCCACCTATATCCGAATCGAATGGAGCGGTGAGTCAGGACGCGACGCTAGCAACATCGACGCGCATTGCATTACCCATTACCCAAGACATTGCATTGATGCAATCCCCAAAGACCCCACCCCA
AGTCATTCAGGTCCCGTCACGTCAATCGGCAACCCTTTTACTTTACTTTCCGTTTTGGACCAGGCATGCCATGCCCCAATTCTTACAGAGGACCGGACCCATATCGCGATACCTTGGAGTGGAGAGTGTATGGGATTAATTGCCTTGCCAATCAATGCATACAGCGAACCG
AACAGCCTAAGAGGTGCCAGCCAAGCCATGTGCGCCACGGATTTGCAATGCAAACTATTACGGCAGCTATACGATACGAACCGTACACGATAGTGATTTGATTTCCAACGGGGTGCTTCAGGACAGGAATTGGATTCACGTCCCATGGCATGGCGGAAAAGTGGTCTTCT
CATCTCATATTCGGATTGCAGCGTGGCGTGGCGCGTAGTCTTTCATCCCCTGCCCTGCTCCCACCGACGCGCTATGCCATGCCATGCCGCGAACAGCACCTTCCCTTCGAAAACGAATTATCCTGATGTGATGGCCTCCAAAAGAGTGTGACGTGACGAGAACGTGTGA
GATCTGCGATGCGATGGTATTGCTAAGGGTAACATAACAATCCGGCTATATTGTAAGCGAAGCGAGGTTAGCAGTGGGCTTATCTTATCTCGAATTGTGTGGCCCTAACCTAACGATTTCTGCGGAAACAGGCAAGGCAGCATCGGATGGGGACCCTACCCTACGGAATAA
CGATTATCCTTTCCTTTCAGGCGGCTACAGGATTCAGATCAGATGGGGTCAACTTATAGCGAAGCGAAGGGATTGAACTGATTCATTGCATTGACCCCCTAGACCAGCTTTGTTTTGTAATGGTCCCCCGCTGGTGTCGTGTCTACGATGATCTGTAGCTAGCCTAGCATGT
AGAGTATCTTTATCACATCACACCTAACATCGAACTCTCGTCTCGTTTCCCTTAGCAATTGCGAATCGAATCTAGAGGGGCTCACTAAAAA
>6mers_1_curlcake_2
AAAAATACAGACTTCTGGTTTTAAGTTAAGGCTAGACGTGGACGGCGGTACGGTAGCCAGTTCACTCTGCACAAAACAAAAGCTTCTGAAGTACAATTCGATTCGTGTCAAATACCATCTCCTACCCTACCTGTCTCTAGCCTTTTATGTTATGTGGAGATTCGGACGTAGC
CTCGCCTCTCTACCCTTCGGTCGTATCCTATCCCGATCACGTCGAATATAGTATAGTAGAATATAACCGGCCAAGCGTAGCGTGTATCTGTTCTTGGCGGTCTGGTCTCCTACTCGTATCGCAGCGGTGCGGTACAATTGAGATAGAGACCTAACCTAAAAGTTTGTACGCG
ACTACGCTACGGGACATAACGACAAGAACAGAACAGATAAGGATAGATTATATACTATACTACTAGATAGGGACGGGTTAGGTTAGTTAGAAGTCACAATTCATGACATGAGACAGCATTTAGCCCATCCTATCCTCCTAAAGCGTGGGCCAGCTGAGCTGCAACTTGCCAAA
CATTTACGTTACGCCCCACCCGGAAGCCTTAAATTAAATTGTTAAGATGCCCGTCAAGTCAAGGATGTACAGATCTGGCCTGACCTGACCTGTATAATAAAAAGTATAGTATAATATAATAATGATACTGTTCTGTTCGATACAACTGAATCTCACATCACATATGTTACGTTATTA
ATCTATTCTATTTGCTTGACAGAGTCATACTATACTGGAAACCTACACCGAAGATTAGATTTACACCTTGCGCCGGACGTAACGTAGGATAACAGTCAAACTCGGGTCGGGCAGCTGTCTTAGTTTCAAATCAAATACGCGGGAACTACGGTAGAATAGAAAAAATAATGGGG
CTACCTTCCCTTCTAGAGACCTACCTGAATGAGATGAGTAATTGGCGATACCTTGCGATGCGAGCGGGGGTCGAGAGAAGAAGAGAAGGCAAAAACTGGATTTCGATCCGATCTACAGAACGGAGAGTTTCCATTCCACAAAATACTCCGCAATTTCGTTTCGCGGACCG
ACTCACCCTACCATACCACCCACGCGCCACCGGAAGCTAAGCTGATAACCCTAATATTTTGGTTTGGTAATGGTTCTCACGGGCAAGTCAAGTAATCTATCACGCGATATATAATATATCATTCAATATGCGTTGGAGTGGAGGTAGAGCCTATGGAGCGTATCGTATACCGGT
GTGGTGGCCATTTCATTTCGCTACGATAATCATCGGCTCGGCTCCGGTCATATTCACAGTGCTCTGCTCGGTCCGTAGAGATACTGTATTGTATCGATGTACTAATGAAGTATAGTATAATTTTGGACTATCGTGCGCAGCGCAGCATGGGTGGAGGTTTCGTTTCGTTCAAC
GTATGAGAAAACAGGGCAGGGCGCCTCAAAAAACCACGTTACGTTATTCTGATTAGGATCAAAACTAAACTTGTTCAAGCCGGGAGAGATGAGATGAGCGGGATATTAATCGATGAGATGACGCCGGGGAGCTAGGAGATCAGATCCTTTACCATAGGTATGTCCAGTCCA
AGTGAATAACACGCGAGATGAGATGACGCCTGTACAAAGATTGGTTTGGTTTCATGGATGCGTCGGCCCGGCCCGTCCCGCGAATGCAATAAGGTAAGGTCTTCATAATCCATATACTCGACTCGCAAACGGTCTTGGGTGGCAGGGCAGTCTTTATTCAATGGTCCATG
CCATGGTTCATAAACGAGTACTATTCTATTTCAAAAATTGTTGAAGGATTGGATTTGGTAAATAAGCTTCGCAGGGCAGGACTTAAGTAACTATGTTACATTACATAAGGTATGCCGGATCCTTCCCTTCACCAAACGCCTCTTAATCAGATCAGGTTTGATTTTCCGGAGCATT
GCATTGTCCTAATGAAGTCCGTATAGTATATTAGATTTACCTTGAGTTATGTTATCCGCCACACTAGTACACGTTACGTTGCGTAACAGAGGCTGGACCCGACCCCTGTTGGAAATCACTTAAAGTAAAGGCTGTCAGCTTAATACTTGGCTTGGGGTCCGATCGCGTGTTAG
TCTAGTCCCCCTTTTCGACAGCGCTTTGCTTTCACCTCCACAGGAGCCCCTACCCTACATCATAGGCTACCCCAGCTCAGCTCAGATACAAGCCGCGTGCGATGCGAGATGATCGGGAGGAGACACCACACCGATCTTGCATTGCCGGGAGTGGAGTGAGCCTAATTTA
AAGTCGGCTCGGCCTTGATGCACTGCCCCCAGGCCAGGGCTGCTTTATCGGTTAGAAAAGAAAGCCAATGCGAATACCCGGGCCGGGCCGCCAATGGAATCAAGTTCTGTTCTGGGCAAGGCTGTTGGCGACTCGACTGAGCTTTTGATTTAGACATAACATAGGTCCT
CCCACCACGCCGGCCCGGCCTCGTTTGAATTAGGAAGACAAGACTGTCCTCGACTTTGTGCGCGGCGCGCCTGGTAGCACACAATACGTTACGTGATACACAGCTATGACATACCATACTCTCGACTTGGGGACTTTACTTTACGTTTTAACATGCGGGCTGTCCTGTCT
GCTGGCCCAGAACATTGCCTTGCCCTGCTTACGTTGCACTGAAGTGAAGCCTGTGATAGTGGAATTTGT
>6mers_1_curlcake_3
TTTGTTCGCCGTATTGACCACTGGGCTGGGCTATATTGACACCTCACTAGACTAGATAAAGTTGTATTCCCAGACCAGACCGAGGTGCCGATATACTTAACTTAAGACCCGATCGATACGCAACGCAACGACCCCTCAGCAACCTTTAATTTAATAACTACGGTTCCGCTGC
GCCGCGCCGCTAAGATATTTATTGCGTCGCGTCCCTTCCTAGATCTCATTTACTTTACCGCACTCTTACTGGTCGCCCCGCCCTTTAGTACAGTTAAAATGCCATGCCCCGGCGCCGAAAAACCGTCTCGTCTCGGCTGAACTATTCTGATGAGATGAGCTCCTGTATGCG
ACGTGATGTGATGTGTTAATCCCGTGAAAGATAAGATGGAATTCGTGACGTTCGTATCGTATGTCATGCTAATCCATCTCCCCTCCCAGTGCTCCCTATCATAACTTAACTTTCGCTGTGAGGTGTACTCTACTCTAGAGCCCAGGTATAGTTTGCCTTGCCGTATGCGCTCT
CAGCTGTAGTGTAGACGTCCAGACGATCTTTAATTTAATCGTCATGCCAGCGGCCGCAGCGCAGACGATAAGATAAACGGCCGGGCCGGAGCGGGGTATATTCTACATGACATGAATATATAACCCCCGCAATCCAATCTGGACATTAGCTGACCCGCACCGCAGCTGAA
CTGAAAAAGGAGAAGAGAATAGTCAACAGCAGGAAGCCGTACCCCTCCCAGATGAGCCGAGAGGAGAGGACCTTTCAACACCCTAACCCAACCCTTCTAACCTTTTTCTATCCGATCCGTACCACAGAAGCACACGAATCGAATCTGTGCAAAAGCTAACGTGCCGTGC
CGACAGAAGTTACTCATTACATTACGCCTAACGGATACGTCCTTGCCTTGCATCGGCACCATCCTCTTGTCTTGTAAGCATAGCTGCTCTTGAAATGAAATGGCGTAGATGCTTGTTTTGTTTTGGGGTGATTAAAAGGCCTGACCTGACACCGTCTAGTTCGATTGTTGTG
TTGAGGCCCAGTAGCACTTCCACTCCACTCCTATAGCGTCGGCATTCAATTCAAATAGGCTCAGTTTTGACCGGACCGAGAAAGCATAACTGGCCCGGCCCGGACATACAGCTATTTAGGTACGGTACTCGTTGGCACCCCTTAGGACAGGACGTAGTGTTCCCGCGCAT
CGCATCGCCCTCTCTCGCCTTCTATCCTATCCTTCAACGGCTGGACCAACCGCACCGCGCTGGGATATCATCGAAATCAAATCTCATGGCGGTGCCCGGACTTGACTTTAAACCAAGGATTGTAGCCCAGCCCGAGCTGTACAAGTGACTTTCCTTTCCCCCAATACCCC
GTCAGCCAAGCCACTTTAGATTTCCGAAACTATACTATAACACTTCGCAAACCCGAATCGAATAATCGCGCCATAAGACTAGGCTAGGTTCGCGCTCTGCCCAAATCGAATCGTCAGCTTACATTGTACCCTCCCCTCGGTACTTCATCAGCCCCTCCCCTCCGAACGAAGT
TCATCCTAAGGTAAGGATCGTTCGTGCTGAAGACAAGACAACTAGCCTAGAAGGAACGCGTCGCGTAAAACAGTTTCGGTTTGTGCTGTGCGTTTCGAGACTGGGTAATGTAATGTTCAGCGCAGTTCGGGTACAGTACACGAATCTTATATGGACGAGGCGAGGGAGC
GTTACCACTTATTCTTTTCTTTTCGTCTCAAGGAGCCGCGTCGCGTATACTTGGTTGGGTGCCCCGCCCCGCTTGACCTCCGCCTGGATCGGATCGAACTACGTAACTCAGATGGAATGGATGAACTTTTTAAAAATCAGCTCAGCGGTGGGTAAAACTTTGAGACGAGA
CGGATAGCTACAATAAGCTCTGCTCTTACGGTTACCTATTGAACCCAACCCAAGCCACTCCTTCCGCCTATCCTATTGTCGTGAGAACGCTAACTCAACTCAGGAGCCAAGTCCTCGCTAAGCTAAAGGGATGGACTGTCTGCGGCCTCGCGCGGCCTTAAGGGATAACC
CAACCCTTAGTGGCGCTTTCTATTATATTATCGCAGTTGTCACGCGCAAGGCAAGGACTAGGGGGCGGCAGTGGCATGGCAGGTTGACAAGAGTTTGAGTAGAGTACTTAGCCCTGGAAAGAATGCAATGCCTACCGGAGGGTCCGTGCCTTGCCTCGGAAGCTGTCT
CACCGTCCCGTCCGCAGGTAGTTGCTCACAGACCAGACCTAGGTGAAAGGAAAAGATTAGATTAGTCCCTCAGCGATGTAATTTAATTGATGAGTGAGCATGTTGGGTTGGGTCTATGGCATCGCATATTGGGTTGGGAACAGTAACTGGCATTTTTATTTTAATATGGAGT
TTGACTTGATATGATATCCAATCTGACGCACGAAATGAAATTCCGATGCGTGTTAGGGCTTGGCTTTGTCGTCACTATCTTGATAGGATAGGAACGAGAATACCAAATCTCATCTCGCGTAACTAAGGGCCGTCTGGTCTGGGTATGCATGCACAACCACTCCACTTGACAG
GAAAAAACGAATGGAATGGTTATAAGTTGCACCAGTCCAGTCCACCCTAAACAAGGCGATATTCTATTCCTTGCTAGAACTAACGGAAA
>6mers_1_curlcake_4
GGAAAATAAGCCGAGTTGCTTCTAATCTAATACGACGGTTGGTGATAGCGTAGCGCATATATCCAGTTTGGCCCAGCCCATGATGAATGATTCCTGCGAAGCGAACCCGACATGCGCACCGAGGCGAGGCGTCTAAGAACCATATGCACGGCACGTGTGTCCGAACCGAT
TTAGATTAGAGTGCTTATCTCGGTGACAGAACAGATCCCTTAACTCCGTCCTGACCTGACTACCGTAAATCTCTTCCGAACCGAATGAGGATGTGACTGACTTCTCTTCTCACTCGAGTGCACAACGGCTCGGCTTGGTAATTACTTTCGGGATAGGATACATTTTAGCGAC
ATCAGAGCAGAGCTCCCCAGATAACAAATCCTATCCTAAGTTCCCGGGGTGAGTGTGCTGTGCAAGTGGTATGGCAGTAAACTAAACTCTCATAGAATCTGTCACAGCACAGCATACCAAGGCAGGCATAAGTTAAGTACCGACTAACCTGCCCTTTGCTTTGAAGAATTG
AAAGTTCTCGCTTCGCTCGCATCTGCCGCCCCATATCATATCTGGAAGTGGATCGCATGAAATGAAAGTCACGGTCATACCGCATATCATATACAAACCTAGCCGGCATATGATATGGTGTTAGTGTCTAGTCTATACTATAAACTGGAGGGGGAAGGGTGTGGTGTCCACT
GCCCTGGGGTTAAGCTAAGCTGGTATCGACGCCCCCAAGACAAGACTACAGTATTAAGGCGTCACGTCACAAAGAGACGCAAGGTAGCTCAGCTCTCACCAAACCCCATAAATGAAATGATGTCTCTCGGACCTCCATTCCATTCCCGCCTGAAACAAGCGAGTGGAGT
GCGCTCCGGGGGAATTTAGTTTAGTTATGGTTGTACCGGAAAGCCCAGCCCCAGCCCGTCACTGTAGAAACGAAACCAGGAATCAGCCGCTAGCATAGCATATTAAACTGCCAAGCTGTGGTGTGGACGCAGGAGTACCGGTAAAGTAAAGCGTGCCTGCCCGAGACA
ATACAATATAGGATTAGCCTTACTCAACTCAACGTTCCGTCGGCAAGTGAGTCGAGTCAGACTTCCATTGTTCCTGCACTGCATGAGCCATCGGAGCCCCCTCCCCTCAATCTTCTGTCGCTACAACGCAACGTCTTGAACAATCGTTTACTGTACTGGAGTCGTAATTTTA
AGAACCGAACCCAGTCACCAGTCTCGATCGAATCGAATTGCTCCTCTTGGCCACGTCACGTCTGTGTACGACCAGAGTTACGTTACAGCGCGCAGTCGACCACAAAACAAACAGCACCGCCGCTATCAGACCAGACGGTGCAGGTAAGAGGAATGGAATGGGGCGGG
TAGGCACAGTAGTGTAGTGACGCTCCTGGCCTCCGGCTTGGCTTGATCCGTAGACTTAGTACATTACATAGGCCTGCCTCACAAGACTAGACTATGAATATGTGCTGCACGGTACGGTACGCGTTCTGTTTTGCTAGTCTAGTCGCCAGTAAGTACGTTGATGAGATGATT
GGTACTTGCTCCCTTGGGTTGGGAGCCGGACAGTGAGTCTTCACTTCAATTCAGTAGTTAGCACTTGTCTTGTGCGACCGCCGTTTTTCAATGCAATGCTACAGCTCGGCCGTTTTCATTTCAGCATCCTCATAACCGCCCTCCCCTCCCTAGGACTAGCTTGGATGTGAT
GTGGGGAAGTAGACGATAGCAAAGCAATGTTGTGAAGATGCAGTGTAGTGTAATAGGGCCAGAGTTGATCTTATCTTGTGAGATGCACTATGTACTCTACTCCACCATACACGTAAAGATGTGATGTGATGGCAACCGGTTCAGTTAAGTTAGGCGCTATTCGACGACACTC
CACTCGAGGAGGGCTGCCGTTATATTATATGCTTCAACATTTGGGAGAAGAGAAGACCCTCTAGGCACCATTACGTTACGGGTGAGCTACGTATGTATTTTATTTCTTTGGTCGGGCCCTCGAGGCGAGGGCGTCCCAACTCACTCTACGCTACGAAAGTGCTACTTCTAC
ATTTCATTTGACTGACGCCAGGCTTAGAGTAGAGTTTATGGGTAAGGCCTTTACTTTACCCTTGTAACTCGGCGTTTGGTTTGGCTAGGGTGTTTGGAAGGGCTGGGCTAAGGTAATGGGCATACGGTACGGTAGGAGTAGGATAGGTAACGGGACGGGGCCTAGCATCA
TTGGACGGTACGGTCCTGGGCAAAACGTGGTGTAGTGTACATCGCCTCATTCCCAACACAACACGGTTGAATAGGGCACAGGCGAGGCGCTTTAGTAGAGTATGAGGAGGAGGTCGCGCAAACAACTAAATGTAATGTCCGGGTTTCTAATCATTATGTTATGCTTTTC
GTTAGCCGGCAGAACAGAAGTTCGTACCAGGATCATCCCATCCCCTCCCGGCCATAGCGTTCAATTCAAGTTGACGACAAATTGCACGTCACGTGATCACATATAGCTTTGGCTTGGCTTCGCCTTTGGGTTGGATGGAATGGAACACAGTCCTAGGGTTGATTTGATTCC
GCGTCTTTCTTCTTAATGTAATGTTCCAGACTAGTCGTTCTTCTCTTCCCATCCGCATACTGTTCCGTTCCGTCGAACTGCTGCCGTCCCCAC
>6mers_1_curlcake_5
CCCACGTAGCGCGGCTAACCAGTAAAGTAATAGACTATCTTCGGAAGAGAAGAGAATCCGTTCCTTACCCGCGGCGCGGCCTTCATCGACTGGTTTTTGCTTTGCAGAGAGCGCCCAGACGAAAAGAAAATGTCGGTAATACTAAACTTGACTTGTATACCAGTTAACGAA
GTTTAGTTTTATAAGAAAACTAACAAACTAAACTTCTCTGAAATAGCTTAGGACAGGACTGCCACTATGTGACCAACTTAACTTGCAGGCCTATAGTCCTTATATTATAGATCAAATATTTTGCCTTTACTTTAAAACCGAAGCCCTAGTACTGTACTGACCTTAGACCACGGGCT
TACCTTACGGCGCGCTTTGCGGAGAATGGAATGTGTGAGGCGACCGTACTGAGCTGAGAGCGCTCACCGGGGTAGGTTAGGTTGACTCCCCCGAACGGCGCGACGCGACCCTCCTCATCTGACTAAGGTAAGGCTCATTTCTCAACCTCTCTGCTCTGATCCCTCTTT
AGATACTAACCTAACTTACGTAAAATGATTGCTGGGCTGGGCGCCATAACACACATGAGGTGAGGGGTGCTAGGAGCAATAATGCAATGCTGAGGTACCTTAAATCCAACCCAACAACGTTAGCGGCGAGCCAATCCAATGATCTACGCTAGACGTGCATTGCATTTTTGC
CGATGTTCTCCATCCCATCGAGAACACAAGCCTGCGGCCCGGCCACGACTATAGAAACACCCCCCCCCCCGAAAGTGTACCAGCGTGGTGTGGTTCATGACCTTAGGGTAGCGCAGCGCACTACCTGCTCATGGATACCATACCCTGTGGGCCAGAAATTAGTATAGTA
GCCGTATCATTAATTATACAATACATTAGCTTCCCTGTCAGCTTTGCTTTTGAGGAACCATGCGCTGCTATGCTACCATGCTAGTTTCACTGAAATGAAACGTAACCTGCCTGCGCCTATCCTATACTTCATGGCACCCCGGCGAGGCGAAGTGGAACGACCCCCCTGTCCT
GTCCGGAATCGGTTCTTCTTAAGTTAAGGGCGCTACGGTCATGAGGATAGGATAGTTGGTGCACCACAATGGAATGGATAAGATCGGTATTCAACGCGACGCGAGGACACGTAGATTATTGCCTTGCCGTTAATAAATCCCTCCGACTCGACTCAAACAGGACCTTCGAGC
GAAGCGAGACCTTTCCCATCGGCGTCCCGTCCGGGCCGATTACTCCGCTTTACTTTAAGGGTGCGATCCGGTCCTGACCTGAGCGAAGAAGATACGTGGTGGGGTGGGAGTGGCGTGACATGGCTGAGCTGACACCCAAAGTCCCGTTGAGCTGAGCAGCTTTTTTA
TAAATGCTGTGCTGTCCACATCTTCGGTAACCAAACCAAGGAACACGAAGCGTCTTGTATTGTATGTCTGACACACGCGTTGTTTTGTTTTTGAATCCGAGTCCTATGATATGAACCAAACTAGCTCAAACGCAACGCATTATTGGCAGAGTCGGGGGGGGGGCGCAAAA
TTTTAGCACATTCCATTCGGTGACTCGGAGAAGGCAAAGCAAATCACTCTGTTGCTACGTCCAGTCCAGTACCTCTGACCACCGAAGCGAAGCTAAGCCCGCCTTGTAGAACTGAACTCACGTCCTCCCCGGAATACTATACTGATTGAGTGGAGTTCTCGTCTCGTCAC
TTAGCTCGTAATAAAGGCAAGGCGACGGGACTCAAAAAGGTAAGGTAGCTCCACATCCGGCGAAGGGAAGGGGCGACGTTAGAACATCCGCTCCGCTCCGACAACAATGCAATTACATTACAATGAGTAGTAAAGAGGGACGGGACGCGGGCTTTATATGACGTATAG
TATAACGTCAGTTCCACGCAGCCGAAGTTAGCCGTCCGTCGAAGTAGGCTAGGCCTGGCGAGCAGTTGTGTGTTGTGTTTGATGGACGGAACTGCAGGTGTCGCAGGTCAAATCTTTTTTGTCGTTGTCAAGAACCTCGCATAGCTCGCCTCGCGGTTGCCCTAAATGT
GTGCCGTGCCGACGCGGATGCCCAATACATTACATCCAACGCTACCCGCGCCTCTCCTCTCAGACAGCCACATATCACACCACACTGGGTCATTCTGCTACTCTCCTCTCCTTTGATCGATAAGCCCACACCACAACCTCAAGGCCGTCTCCTCACCTCAGCGTCAGGT
GTTGCGACAGCACAGCCCTCGGAACTGACCCATGTCATGTCGAGCCGTAATAGCAGATTGCATTGCAGTATCAGAGAGACTCGCTCCGCTCACGCCCACTAAGTTGTTCGGTTCGGCACACGGTTGGGCGTTCTACTCTACTATCTTCCCGCTCCAGGTGAGGTGAATA
GAATGACTTATTTCCTTTCCTTCCCTGCGTTTGCGTGCATCGCATCGTGCGACTAATGCCCACGTTCCGTTCGAATCATATGGTGACAGAATAGAATCGCCATAATAGGGGTCACTACACTACGTGTGGCACGCTTCAAGAAAAGAATAAACACCTCTGGCGGGCGGGGCG
GACTGGAGCTCCAAATATGAGATGAGCTTCGTTAATTACTGACTCAACTCACTGTGACTCGCTCTCTTAAATTAAAGTGCAGCAAATTCCATCTGA
>6mers_1_curlcake_6
TCTGAGAGAAATGGATATGTGCTTGGCTTGAAAACATTTCCCCGTACCCCACCCCGTTGCATAGATTTTACTCAGCTCAGTGCCTTCTGTCCCCGCTATGCTATGCACATTCCGCAGCCATCGTCTACTCGTCGACGATGGCAAGCGCTAAGCGCCCTTGACGAGTTCGG
AGTGGAGTGGCTGGGTCCGATTAGATGTAATGTAACCAGCCATTATATGCTTTTATTTTATTAGGTCGATGTTGTGTCCGGTCCGGGTCCCAATTGAGAGTGGTGTGGTGATTCTCTCCACAGAGCTTGACTTGACCTGTGCCCCTATTTTGTTTTGTTTCTGGGGATCTCC
GTTGCGCAGCGCAATTGCAAAAATGGACAAAACAAAACGCCGATGCCTGAGTAAACCTAACCTAATTGGAGGAGACAAGGGACGGGACTATATCGGAATCGGGCTCAGCTCAGCCGAGATAGAAGCTACACTCCACTCCCGTAATGTCAGTCTAGCCAAGCCAACGCTG
AGGTACTGCGGTCGGGTCGGTGGATTACGCAGTGTCTTTTCTTTCGTGGCAACGTCGTGCTTGCCTTGCAATCGAATACCCTCATGTAGTGTAGATGTTTCTTTTCGAACTTCACTTCATTTAACCATTCGATCAACACAACACGGCCGCCAAAGTAGCGTACCGTACCGA
ATTTCTATCGAGATATAATATAAATAGAGGTTTATGTTTCAGGTCAGGATGCCAGGGCTGCATGCGAGGCGAGTAGGCCGACCTCCTAGGAAGGGAAGAAGCCAGAGTATGCGGTTAAGTTAAATTCGTCGGATGCTCTAAGCCAGTAAGCTCTGAAGGCGCGATCCACGA
TCTGATGTCTTGAGCATAAAGCAAAGCACAATATCTCTACTGGCGTTGCGTTGGCCGACTTGAAGCGGAACCGAACCCCTGATCCATCTTGTGGCCGGGCCGAACAACCCAAGTATCTCGCCTCGCCAGAAATAACGCAGACACATGACATGATGCTCTAAACCACTCAA
CCCAACCAGGGAGTAAAACATCGTAGGGTAGGCGTATTATGATACCTCGAAGCGAAGGACAATATCGTGTAGTGCCCTGCCCAAAAGGACCAGAATTGTCGTGTCGTAGGTATTCATGTTCGGGGCCGGGCCAAATGTAGTACGGACAGGTTAGGTTGTCCATAGTATCC
CCTAGTCTAGTCCAAGCGCGTTGCTAATAGATTAGATAGCCCTCTCCGCTCTGAGTGGAGTGACGTTTAACGCATCTGCAAGAGCAGCTTGGTGCACCTGGTCCTGTCCTGGCGATAGGCGAACACACCAGACCAGACCCTGATTAACAAAGGTTAGGTTACGATGGGC
AATGTCATTTCATTTCACCATGGTGTGTCGGAGGGTAGGGTATATCATGAACCACGACATCCCATCCGCCGAAATATGACTGCAGATCAGATTTTCTACAGTACTAGAATGTAATGTGTCCCAGCGATGAAGAGAGTAGAGTCAACGGGTCCAGCGCGAAAGGAAAGGACG
GGGAATGTATAGCGGAGCGGAAGAGGGCAAAGGTCGAGCGCAGCGCTTCACAGATATAACGAGTCGAGTCGCTGAGACCCGCAACTAGGACAGGACGAACGTTTCATAATTATTTGATTTGAATGGCGCAAGGGTTTAGCGTAGCGATGGCCAACAGACATTCGAGTC
GAGTAACCTGGTCGCACACTCAGCTCAGAGCTAAATATTACGCAGATGAGATGAGAGCTTGTTGTTTAGGGAAGGGAATGCGTCGATGATGTTGGGACGGGACGTTGGTTTATTTCGGAGGCTAGGCTTTACTTTCCGACGGACGAGCCGAGCCCAGGTCACACTAATC
AACTCAACTACTGCGCGCGTACAAGGCCCGGCCCGATTGTGGTACGCTGTGTACTGTACCCACATTAACTTACCCCCGTCCCGTTCCCCGAGGAGTTAAGCCACGCCACTCATTGATTTGCCTATCGCCTCGCCTAGTGCAGGAAACGGTATTATATTAGCCTCACAGAA
AATCGACGAGACGACGTCAATAAAATTAGACATAACATAAGTGACCATTTGCATAGAGGAGAGGTGCGGGTATTTGGAAGTCCCGTCCCGGGCATGATACTATGGTACCGTACCAGCAGAAAGACCGTCGCGGTGCGGTAGTATTGGTGATTATAGGGAAGGGAGAGAAA
ATTTGTAGCTGGACTGGACTGGGACAATCACTTTTCTCCTCTCCCGAATTTAAGTAAGCGCACTGCACTCCTGTGGGAACGTATCCAGACCAGAATAGCCCGTTCAGTAATCCGATCCGTGAGCACTCTGTGAATTCTCTTCTCTAGGGTGTCAGCTAGGTAGAGTAGATC
GCCCGACCACACAAGTCGAGTCGTCGGACGCTAGTTCCTGTACTGTACCTGGCAATAGTCAAAATGGCATGGCTTGTGGTTCTAGAAGGTTACGTTACAAGGTTTCCTGTTAGTGTGGTGTGGTCTAACTGCGGTGTCACGGCACGGCCAAGGTTACGAGAGGGTACC
GTACCCACCCGGCGTCCACTAAAGCAAAGCCAGTGGGTCGTCCATGACATGACATTCCCAGTTGCCGCTTAGAGCAGAGCACCGCCACCTTGTCCTATTT
>6mers_1_curlcake_7
TATTTATGATATCCCTGAGGGTTCCGTTCCTACAAGCAGTATTACATGGTATGGTAGTTCTAGTGCACTCAATGTAATGTTAAGTGCCGGTCAACTTCCGTTCCGGATAAAACACTATTCCACGCCACGCATTGAAGTTACCACGATTTAATTTAATTTGTGCAAGTCATACGAG
ACGAGGTAAGACAGTGTCGATGGGATGGGACTTGATCAGGCGAATCGCGACGCGAAAGAGACACGATGCGGACCGGACCGAACGGAGATCACCCCACAATACAATCACACTTTTACATCTTCGTCTCGTCAACCTATCGTTAAAGCTCTGCTCTGCAGCCTCCAACTGT
GACCGCACCGCGCCAATAGAGAGGATCCTGGCCTGGGAGCAATACACATGATCGTGTCGTGTCACTAGTGGGTTTACAGAGCAGAGGCTTGCAAACGACGTCGTAGCGTAGTCATGTCCTCAATCAGCACGGCACGTGCATCATTAGACCGGTAGTGTAGTCAGGGGC
ATCTCCGGGTCGAGTCGACTCTATATTTTAGGTGAATTGAATTCTTATGACATCAAAGTCTGTTCTGTATTCCGTAAGTCCGCATGGGATGGGACCTCAGTGATAGTAACGGCACGGCACTGGCTAGCGACCGATAAAATAAAAGCACCTAAGCTAAATGACTTGACTACCAT
TTTTCCAAGAAATTAAATTATCTATCGGGCGTGAAGTGCGGTGCGTTATAGGAATGGGTTGCCACGCCACGATGTCTTGCGTACGCCCAACCCAATCCAGCTCATGCATCAGTGAAGTGATCTTAGCTGGCATGTCGAAACGAAACGCAACAAACGGTCTGGATCGGATCT
CTGCCAGATTGGACACGTGTAAGTACGTGGGTATCCTCTATACCATACCGACAGGATGCAACGCACATCACATCGTACTGAGCGTAGTAGTGGAGTGGACTACCCGTGTAAAGAAATAGAATAGTATTTGACTAAGTCGCCGTGCCGTGCTCCTGTCGTTTCGTAGCTGAG
CTGGCCTCCCTTGAAAGTAGTTGAGTTGAACGTACGAAGCACTTGAGTTGAGTTAGTCCGCTGTTTTCCGGTCCGGTCGGTCTCCTCCTTGGTCACAACACAAGGGGTTCTATGTATAGGAAGGGAAGCAAGATAATGTCCGCGGGAAGGGAATAAGAGTCGAAGAGCC
TGTTCTGTTAATGCATTTCAGCAACGATCCGATCCGTTCCATCAATCCTGCGAAGCGAAACATAAAAAGCGCCGCCTTACCTTATGCTCGCAACGGTGTCCCCTCCCCTACTAATCGAGCGAGACGCACCATGCGCACGAAATTCACGACGTCAACGTCATCTAGGCAACT
CTTTGGGATGGGAGACTTCGACAATGAAGCGGTGCGGTACATATGTTGCACAGGGCGTGGCGTGAAGGAGTCCAAGCGGGTGGGGTGGTGCGGTATAAGGTGGAAGGTAAGGTGTGATGCGAGTATAGTGCAATGCAACCGCAAAGCCTCGGTCGACACGACACTG
CGCCCACACACGCTGCGCTGCGACTGTAAGAGGAAGTGCCGGGCCGGTTTGGATTCAATATCGTACAGTACACTAGTAATTCTTGCCATGCGATGCGTCGTTGTGAACTACATACATTACATGTGGATCAGGGAGGACAGGCCAGGCTAAAATAATGCTCGCTTCCTTTC
CTCTACCAACTGTTACGAACCTAACCTTTGGTGGCCGTGTGCAAGAAAAGAATCATATCCGAGTTCACTTCGCTTCGCTAATGACCGGCAGTTGCTCTGCTCAAGGTTGCAACAGATACTCGACTCGTTGAGGCGGCATTGGACTCTACTCTACCGCAGGCCCCATCCCC
TTCCCTTCTCGACACGTCGCTGCCGAACCGAAGTCACGGCCCTGCATCCATGCCATGTCGCCCCGGATTCTCTTTAGTTTAGGAGTTGGATATTGGCAGGCCAGGCCGGTCCATGTCAAAGCGTAATGTAATCTAGAGCGCTCGATTTTCGCTTCGCGTCAAGATTACCG
GGGAGGTGAGGTCTTAGCGGATGGTCCCACCTCACCTGCTGCCAACGATCATAGTTGAGTTGACGGGATTACTCTTCACGATACGATTGGCTGAAACGTATTCCGGCCCGGCGCATGGAATCCCACACAACCCAACCCGGGTGAAACGTGTTTTTTCTTTTCTCTTCAAA
CCCAACCTTATATTATAAACTTAGTGGCGTACCTACTCTACTCTGCTTACGAATGCCGTGATGTGATAACTGTGCTGGAACGTCATGTCATGTTTGCAGAGAGTACGTCTATTCTATGCCTCTGGTAGACTTATTCTATTCTGGCTTATCCAAAGTTAACCGAACCGGAGTCTAT

TCAGAACCTTCACTTCAGACTGCACGGCTCGCGATGCGATGCTGCGTAGGGCACGGAATTCCATTCCTCGTGAGATTCCTATCAAGACAAGAGCGCCGTGTCACCCATACCGTACCGTGCCTTCAGTCAATACAGGGCAGGGACCATGGGTGTAGTTTTTTGTTTTGTC
GAGTGTTGTATCGTTAGCTTAGCGTTGTCAGGGCCCATTGAGATGAGAAAGAAACCGTACGGGGCTGCGCTGCGTGGACATGGAAGGTGCTAA
>6mers_1_curlcake_8
GCGTAGCTAAGTCTGTTGAAAAGGCAAGGCACTTACTTAACCGATAAGTGAAGTGTTCTGAGGGAAGTGTGATGTGATGTGAGAAGGTGTGCCATGATAAGATAATTCGCTTATCAAGCAAATTTAATTTTGTTGCCTAGTCGGAACTCGACTCGGGTCTTTGTACTTCGTA
CGTTACGTACTCGGACCGGCTAGACCAGACCAGTCGACTCCGTGTGTTGCGTAAGGCGGCGTATTATCCACGGAGGCGAGGCGCCTACACAAAGAATTCTGAGCTGAGTCTCAATTTGCATGAGCGTTGCGTTAGGCAATTAATGGGTAGAGAAGAGAAGTAAGACGCT
TAGATCATCGACAGTCATCAATGTGGGGTTGCCCTGCCCTTATGTCGCACAACTGTCGGGTCGGTTTGTAGGTCCCTATTGTAGTGTAGCGGAACCCACTCATACGAGTCGAGTTGGAAACCGAGGCCCCCACGCCACGTTCTTGCTGATATCTCGATTCGATTGACAGG
TTTTCCTTAATGCGATGCGGCTGTTATCAACCCTCATGACATGACCGGCTCCAAATTCGCAATGCAATGTAGGAGCACGGCACGTACACTACACCCATACCCCTCTTGGTTATCTTATCTAGGCGTTCTCGGTGGTACTGTACTTTTGCGGCAGTCTGGCTTGCCTTGCCTT
GTCCTAACCTAATTAAATTAAAGGGTCAGGATTATGCTATCATATCACGGGTGAGGATAATACAACACAACACGCAATCGCTTCGGGAAAATAAAATTGGGCCCTATCTGCGATTGCATTGCTTGCGGTCTCTCAATTAAATTAAATAGCCAGCACTAGCGCACTTGACTTGGG
CTTCCAAAAGTGAAGATTAGATTCGACCCTGAAGCGACGGCATGGCATCCAGCAGGGTTTGGCAATCTAATCTAAGTGTTATGGGCCTGGGGCGGGGCTACGATCCACTGGGCGATTCGATTCGAGACTCGTTTACGCATTAAATTAAGCCTGGTTGTCTACATGTGGTG
TGGGATGTACTAGGTTCATAGTATAGTATGGAGGCGGATTTCCTGGGATGGGATCTATTGTGAACAGGCTGTGCTGTGGATGAAATTAGAACGCCAACCCAACAATTATACCTAGATGTTCGGTTCGGCCCTGACTAATCCTTGGAACGGAACCACCCCATGCCACTAAGAT
CAGATCGCTTGTCTTATGCAAAGGGGAGGGGGTACAGCGAAAAGACACCAAACCAATGGCCGGTTTCGGGTTGCGATGCGAGGGATGACATGTGCTCTATGCTATGCGCAGAACTTTATAGGTCCGGTCCGACCGTTGGCATCACGCTCCGCTCCGGTGACTTCACAT
TCATAGTTTAGTTAAAATCGCACAGCCTCTATACTATAAGGTCCTTGTTTAGGCTAAACTAAAAGAAGACGGCGGCGAGGTTCGGTTCTGCCTGATAGACAGTCATCACATCACCTAGCAGCCGGGCTCCTTACCTTACTACTGCTCAGAACTTACTATACTAGCTAGCGAAT
AGTGTGCCTGGCCTGCCTCGTGACGAACCGTTATTTTATTTATGATGCGCGGATCGCTTAGCTTAACGCCGCACTCGCCGATGACATGACCAAGAGTCCCTCGCATTGCCTTGCCCGCCGCCGGCCTTGAGCGTCGCGTCGCTGAGTATATACGTGGAAAGGAAAGGAC
TCAGGTCAGTGAACGAAACGAGGAACATTGCACGAGCCTGTCCTGTCGGATATGAAAAGTATGGCCTGGCCGTAGTCTTCCTCGTATTGATTTGATATGAGGTCTCGCGGCGCGCTGCGCTGGTTGGTCTACCGTCATTCCATTCCTCGCCATTTGTCTGTCATCTCATCT
CAGGGGACCATGTAGTGCTCTGCTCATATCTAACTTCGAAACTCGACTCGAAAGGTGGCTCGTGGATTGTATTGTGTAGGCTTTCCAGCTCGTCTCGTCTAGTAAATCGTAGATCACTGAACTGATTTGACGGTGAGGCTCTCCCCTCCCGGCCGGATCCACTGCGTTTG
GTTTGCGCGTGCTTACAAATGTAATGTAATCTCTGGTTAAGTCTACTTCCCTTCCCGTATCAGTACATAATCATCTCATCGCGCGGCGTAGCATTAGCATAGCATGGGATTTTCAAGTCGGGGCCGGGCCTCAGGCAAATTTCACGTACTGTACTACTCCCTACGCTGGTGAT
TAGATTAACTAGTGTAAGGAGGCATGCCATGCTGTTTCTTTGCTTCAGAAGTGAAGTCGAGCCAGGACACTACCATCCCATCCTACTACAAAATCCGCCAGGCCAGGGGAGTCGCGCGAGGACGAAGCGAAGGCCACCACACCAGCCCGGTACGGTAACCAAGTAACG
TTGAAGGGGAGGGGGCCGAGAGCAAGAGATCTACTCTACGCGTGTCGATAGTTCCGATCCGATCGGTACGACTTAGGGGTCTGATCTGAATCAGGGTCGGGGACAGCTGAGCTGCTGACGGACCCGACCAATAGAATAGGTTCGCGAGCGACGACGGCACGGCAAG
ATCCTAGCTAGGTGAACCGAACCGGGTCTACGAGTGAATGCCA
>6mers_1_curlcake_9
TGCCACCTGCAAGGGAAATCTTAGGTTAGGTCAGAATGTTATCACGCTTAGCTTAACATGTTAAACGGTGGTTCCAACTGGTTCCCTGTTCTAGCGCGGAGCGGAGGCCGGGAAAGCGAGATTGAATTGATCGTGACCCAGCAGCAGCGCAGCGCTGCTCTAAAGTTTA
AACCTGACCTGGAATGACCGTTACTAACTAAACTAAAAGCTCAACAAGTTGTCCCTTCCCTCGCCGAAGCAGACCTACGCAACGCACCCGACGCCTGTTATGCCTAGCCTACGTTTTCCCCCCATTTACCGTACCGATTAACGTTACTGGTGCAGAGCAGATGCCGGTTA
CCGCTGATGGGATGGGCGAGTTATGGTTTTATCGACTCGACCCACTTCAGGTACCTATGGCATGGCTAGACGCTTCCTTATTCGGCTCGGCTAATTCCCCACGGAGATTACATTACAACAGTTTAGTCTGAACGCTACGCTGTAAGGAGAACTCTATGAAGTGAAGCATGA
CAAGCTGAGCCGTTTCGTTTCTGGTCAATTTATCTAGGGGAGGGGATCCGGTATCAAGCAAAGACAAGACCGCTGGAGTCCGGAGTACCCTACCCGTAACATAGCCACAGGTTCCGTTCCAGGAGGTATCAGCAATGAGGTGAGGAAACTGTAAAGTCTGGGGGAGGG
GAGCGGATACGGGCGTTACGTCACGTCTTATCGCTCAAGCAGATCGAATCGAGTCTCTCCCCAGATTGTTCTGTTCTTGTTGTTATTTGGCGTGCATTGCATTTGGTCGTGTTTGTTAGGCTAGGCTCTGTAGCAAGCACATACGCTACGCTTGGCTTAATTTAATATTTATT
TTATTGAGCCATGACTGGTGTATGGTATGATTCATTGTCGTCCGTGGGTTGGGTCACCTTCCACCAGAGATCTCATCTCCAAGGGGTACGGCTGCCGCACCGCACCGTTACAGTGTCTCTAAGCTAAGCGGGCGCAGGTGGAAGGAAAGGAAATGAATCCCGACACTC
GTCTGGTCTGCTGTATTGCGCATGTAAATTAAATTGGTTAGACAAAGCAACTCTGCTCTGGATAGAAGCGGTTCGCTCGGCTCGGGCAGGACACGGACGCAAGATAAGATCCATTCTCCGACTTGCCCAACCCAAAACTGCAATAACTTCTTAGTTTAGTATACAGGTCATT
GTGGTCAGGTCAGTGATCAGTAGCAGACGGCTTGGCTTCTCCGCGGGTTTAACGTCTCAACAGTCTCGGCGGGCATACACCGACACCGGCTTCTTCACTCGCGAGCAGAGCAGTGCGCTAGTTCACTTGTCTAGTCTAAATCAAGAGTGCGTATGGGCTGGGCAGAGA
TTCTCCATTCTAACATAACAAGATATAGACATCTGATGATATGATCATGCTGTACGCGCGTTAACTTAACCCCGTTGTAAACATATATAGTATAGAGGCATTAAATATTCGTTGAGTTGATTGCGGGAATACTAGGCCACGCCACCATTAAGCAGGGTAAAAAATCAAAATCAATT
CACAAGTATGAGTGGAGTGGTAGGGGCACTGAGATGGTGTGGTGGTCAGAGCAAGTTACACATGACATGGTGGACCGCTAACGGGGGACGGGACAACTCCAGGCCAGTACTGTGCTGTGTTCAAAGTAGCCAACCATATCATATTGCGGGGTAGGGATTGGGCTGGG
CTCTAGCGGCGTGAGGGGAAGAGAAGAGCCTAAGCATCAAACACCACACCACCGCGGACACTTACCTCTAAGCTAAGAGCACGCACGAGGCTTCGATTCGATTCGCGTCCTTCCATAGCAGCTCGGCAGCGTACAGGCGTAAGTTTCGTTTCCAGGCGGTTTTGTCAT
ACCTTACCTCAACTTGTCAATCACCGCTCCGCTCAGTCCAATTTCATCGTCGCCTCGCCGGCACGCATCTTAAGAAACTAAACTCGAGGTCTGGTAGTAAGCGAAGCGATCATGGTCTAACATCTTGAGTTGAGGTCGTGGGGGGTTATTAGGTTAGGTGTTGATAAATTT
TTACGATACGATATTAGTTATGTGAAACACTGGCAAACTGGTATGACGTGCGTCTTATCTTATACACTCCATAGACTCCTGAGCTGAGACTCTTCTTTTGGGCCGCTTCGCTTGCTTCCTATGGACAATCCAGTCCAGGGTTCCAACACTTCCGCGCGGCGCGATCCCATGG
AGCTGATCTGGTCTGGGTGGCTATTAAGTGGGGGTGAAGGGGTGGGCTCGAAGGCTGAATCGAATCCTGCATAGTGAAGGTCGCACCGCACCCGCCCGCACGGTAATCTTTTCTTTCAGAGGGCCAGGCACTACAATACAATTTGTGATCGTAAGACGTTTCGTTTAAG
AAGCTCTTGCCACAGGTTAATTCAGGTAACACCCAGGTGCTGCCACCTGCTGACGGACAGACCAATCCAATCGGCGGCTCCATTGGGTTCATTTCATTTTTTAGTCGATTCTAGTAGAGTAGAACGACTGAGTTGGTACAGGACAGGGACGAACATGCCTTACCAATCCA
ATAGCACCTGAACGGATCCGGACGCCGGATGAGAACCTCTGTTCACC
>6mers_1_curlcake_10
TCACCTCCAGCCTTCGCAGTATCTGATCTGGCTCAAGTGTCTAAACGTGACGTGAAATGGCCTAAACGCGGTGCTGTGCTGGGAAATGCGATTTGAGTCCAGTCCTTTCTCGTAAACAACCAGGTCAGGTGGGGAGAGGCCAAAACGATTCGATTCCAGTTATAAGTACG
AGCAGAGCAGCTCATCAGTCGGTGTATACAATACATTATTGTCTGACGACTAGAGTAGAGTAGACTGTGCTATAAAGAGCGGATGGAGAGCGACCTGACAAAATGGAAAACCCGTAGCGTTCGAGGCAGAGCAGAAAGGTCTCGTGGGCGATCAGATCAGGCTCCCCGG
TGGTATCCCATCCCATCAATAGTCACCGTGCACCTAACCTATGTATCGGACTCCGTGAGCTCCTGAGCTTGGAGCTGGCGCACCCGGCACCACCGGTATATGCAGGCTATTCTATTGGCTCTCGTAAGGAATATCTTATCTGCCAGTCTCATCTAATGTACACAATGTACGT
AAGGCGGAGCTCCGCTCCGATAGCCGCATCCACCTCGATTCGATCGGATTCGTGCTAGTTGGGGTGGGGGGTTCGCAGAGGTGACAACACAACATGGCGGAGTAGGTGGAGACGAGACGCGACACAGCCCCGATGCTATGCTATCTACTATAATTCACACCGCCCCG
CCCGAAATACGCTCACGACCTGACCTGATAATTCCAATTACTGCTAATGCTAACACTGCACCGTGGCGAGAAGGCGAGTGAGTTCGACGGCATACCTAGTTCGTCTGTAACGCGGTTATCGGATCGGGACCTAGGCGCTGTTTAGCTTAGCTTATTATGGGAACCGCGTC
ATAAGTGTCATCTAAAGGGCGGGGCCTGGCCTACTTATGTAAATTAACAGACCAGACACAGTCCGAGCACGCCATTGGCATTGGGGCAGCGGCAATGAAAAGACAAATGCGGAGTGCGTGATGCTGCTTGATGGCTGATCAAAGGCAGGGTCAACGGGTCTGCGTGGT
TATACGGGACGGGAGGAGGGAGGTTAACTGGCACGGCACGAGAGCTGCAGTCAGATGGTATGGTCTGAGGGTCGTATAGTCGTTTCGTTGTGCAGCAAAGCCCTTTGATTACCCAAGATTGATAAGGCCCGGATATCGACGAACGTATTGATACCAAACAGGCGTAGAT
ACAGTTGCAACTTTCACGATACAACGGGTAGTCAATTCGAGCATGAACAATTACAAATTACCGAAAACAATGCTTTCTGTGTAGTCTAGCATAGTAGCTAGCCGAGGGCCCCCCCCCCCCTTGCGAAGAACAAGTACCGTAGTCCACCGTTCGTACTCCCAGGAGAGGAG
GATCGTCCAGTGTATCTTTTCCGAAACTTTTGAGCAGGGAACAATACCAATCTGTCCAGATCCCACTACACTACACTTTACTACTTGCTGTGGCTTGCACTACGCTGTGGAGACGATCAGCGATCTCTGGACCCAGCACGCCACACCACATCTAATATCAGAATCGGTTCG
GTTCGAAGATGACTGTTTACACGGAAAGATCACGGGCGGTCCACGACTCGACTCTTTCGGCTCTTTGCTCGTCTCGTCTCAGGCCCCGCCCGGTGTACGAGGTGTGTTCCCTATAGATAGGACCGGACCGGACTTTGCCTACACACGCCAACGCCGTGTAATGACGTA
CGTGTCGTGTCGCACGTTCACCGGGATCTATTCTATACGCCAGCGTCTATTACACCACACCAACCGAGCATTCTTGACTAGTCTTGGCCATCACATGCAGTTGACGTTTGACGGTGCATGGGGGGCATTCTGGGCACAGCGGCAACATAGACCATCTCTACGGCATTCCA
TCACCCTGGTCTCCCTGGTGCGGCCCCTTCGTCGGCCGTCGCGGCTGGTCGGCTTTCTTTGGCTTTGCGCGGTCACCAAGCTACTTGTCAAGCCGTCGTGTTTAACCGTAGACGTAGGTTGAGACCGTGGCCCGGAAGGACCCGGGCCTAGAATTACCGTGACTACC
GTGGAACTCATCCGAGCTCAGCTCGGCCATCAGACTGCCTCGGGTCGGGACTCTCTTCGTTAACAGTGGACGAAGTGTCATTCCGAAGTGGTCCATATATGTGGTCGGGTTTAAACAACCAGCTTGGTCATCCTAGTGATACCCAACGGTCAAACGATATTTCAGTCACA
TGAACATGCAGTACGGAGCATCTCATCTCAGTTACCACTGTACATGAAGACGACATTAGGCCGCGGGATGTGTTACAGGATGTCTCCAGATCGGGTACGGTACGGCGGGTTAGGGTGTTCAGCCAAAGTGCAGCCTTGGAGTGTTGAGGTAAATTGAGGCATTCTACTT
GGCCCAGCTTGGCCACGCTCGACACTATATAGGGGGGTGCTAATCTTTTAAGGT
>6mers_2_curlcake_1
AAAAACGACCGATGATCCGGCCACGCCACGAAGGCCCAGGACCTCTAGCGTAGCGCGTTCCCGGTTGGCTTGAGATGAGACTGCAGCGTTGTGGCGAAACGAAACACGAGAACTGGACAAGCGGAGCGGACCCAGATCATTAAGGAAATGAAATGGCTACCGGCAC
TAGTCTTTACTTTACAAGGCGTAGGCAACTTACCTTACCCTGAACTTCTGACCGTACCTTACCTGATCGGAACATCGTGTACTCTACTCCTCCCGCTAGCGATCAAGTTAAGTTAAACGCTTGGACGAAAAGATAAGATTAATATGTACGCAGCTTGTATTGTAGGTTAAACCT
CGTTATGACCTGACCTCCTAGTGAGAATTGACGGCACGGCTTCGGGGATATTGATACACCACACCACATCCCAGTTCGTGAGCCCAGCCCGAAAACAATACGCGATCAGATCAGAGCGCTGTACCTTTAGAGGAAAGGAACAAACATCGACTGCTTCTCTTCTCTGCTGT
AGACGATAATATTAGATTAGGAGCCGTCCGATATGCGATACGATAACATTTAGCACTCATGAGACGAGACCTACCACCCGCTGCTCACCTCACCTTAACCCCCGTGGATATCGCCTCGCCAAGTGGGTATAACACCGAGCCGAGCTGGTGGCCATAACTAAAGTTAAGTTAT
GTGTAGAATACATCGAGACGAGAACTTGAAGAGCTCAGTTAGTTTAGTCGCGGCGGCATAAGTCAAGACAAGACTAGCGTGCAAGTAGAGGAGAGGAGCTACTAGACACATTCTCTAATCTAAGTAAACTAGCGTATGGGGATGGGATTACCGCACTTATACCGAATGGAA
TGCTAAGGTTCCCGTAAACGCGACGCGTCATCTGGATCTACTGCAGAGCAGAGTGCCCATCCTTTAAGTCGAGTCGAGTAATCAAGGCCCGGATAGGATAGGTCCGATGTTTACGATACAAGACAAGGTCCTGGCACTGGCGAACACAACACCCTGAGAGGCCGAGAC
GACGCGACGTAGCAGGACTCCGAAAAACAAAACATATGAGCATTTTCTTTGATTTGATTCATACCTAACGCTACACAGAACAGATCGGTCGAGTTTCGGAGGCAAGGCAGATAAGAACCTATAGTTACCTTACCTAACATCTGCCCGAGTTGACTTGACAGGGGCATTGGA
ATTGAAGAGAAGACGATCCAAGGTTGGACTAGCCTAGCACACAATGGCAACAACATAGCATAGCCACTAAATAGTCAAAAAGAAAAGATCGATCCCCCTTGTGTTATATTATATCGTTCTGCTCCAGCATAATATAATGAGACGGATTGGTGTGACGCGACGCCAACTACCCT
ACATCACGACACGACCGGGATCCGATTTCAATTGAATTGACCTCCAACTAAGATGTTAGGTTAGGCCCTTATTTACCGATTCCGCTCCGCCAGCACACACTACCCACAGCACAGCGTCGGACTACCCAAGATCTCATCTCACGATTAACGGACGAGTGGTGTGGTAATCAG
AAGTGCATAAATTTAATTTCTACATTCCCTGCTCTTGGTTTGGTGGGGGTATCTTGGGGAAAACAAAACGCGGGGACCGGTCAGTAGAGTAGAGCCACCCGCTATACGTAAATCAAATCTGAACGAATTTGTCTTACTGTACTGAAGGTGTTGTATACTGGGGCGGGGCTTA
CGTGGCGGCAGACTATTCTATTGGTCTCGAGTGTTTCCCGAACCGAAGCAACCCTTGGCTCCACCCAACCCATTGCTGCTTCCTGTACAATCCAATCCATTTTTCGAGCTGCCTCTACTCTACCTTTCAACCCATCATGCACTGCACGACACCAAAAGCTATCCGGCCCG
GCCACCGCTTATTCCTGGGATAGGATAAGTGAGGTAGGACGGGGTCGGGTCGCTTTACACCGATTCGTCGTATCGTACTTTTCTCTCGACTCCATTTCATTTTGGAATTCAGCTAACTTTCCTTTCCTGGCGCCAACCTTCCAAACTAAACTTTCGGGCGCCACGCGATTC
TATTCTAATGATGCGTACCACATAGTATAGTACCTCCAACGAACAGAATTCAATTCCCGTTCGCGAAACCACGTAACGTAATAATCCCGTGGTGAGTCTCCTCTCCGCTCCCATACAGTGATAATTTAATTTCTGCGCCAGACGTGGTCACGTCACCTGGGACACAGAATCAA
GAAAAGAAGTACCCGGTGTGTAAAGACGAGACGTATCTCTATACCATCTATTATATTATACACAGTTGATGATCACAGCACAGCGGGACCTACAACTACTACACTACATAGAGTATAGCCCCGTTATTTTATTCAGGCGCGATTTTCGGTAGAGTAGACCAATTCTGGTAAGAG
AGCAGAGCAAGGGAAACGCGTGCTGCACAGCACAACTTGATCCAGTGACCTTCTCTTCTTCTGTTAATTGTTCCCCACTCCACTATGTTTCATTCAATAGAGTGGAGTGTATCGGGAATGCTTAGGAATGGAATCATCAGTAGCCCAGTCGAAACGAAACTACGACCTGGT
CTGGAACTGAACTAGAGCCGTGCAAACTCTCTCCTCTCTTTCTAGTCGGATTATGATGTGATGAGAGAGTAGGCACTTGTATA
>6mers_2_curlcake_2
GTATAACGAGAGAACAAGTGTACCCTACCCGCCCTTCTCTAAACCCATCGCATCGCTATGGTATTTGATCCCCTGCCCTGTGTGGTTTAAGAAAGGAGTTGAGTTCTACTGAAAATATATAACGGAACGGAGTTCTCGATTGCTGATTCGATTCGTCCTTAGCTATGGTGGTA
GAGTAGAGGGCAGCCGGGGACCTAGTATAGTAACAATTCGGTTAACTTGGGCTGGGCTAGCTAGTAGTATGTGTTGAGTTGAGATTGATGCCCTATGTTGCTTTGCTATATACCCGGAACAAAAAGTAAAGTGCGAGTTGCAAAACCTGTTGTGTTGCAGTGTAAGGTTAG
TTGGCCGAGTATGGCCATCGTATGAGCGGGGCGGGGGGATTAGCTGGGCAATTATATTATAAAACTCTGGTAGTGGCAGTACAGTACACGATGAGCCCCCACGCCGCGCCGTTTTTCATCAGCTACCCCTACCCTAATAGCCGCGACCTGCATCGGATCGGCTGGGTTTT
CACCATACTTAACTTAAACCGCCGCTCTAGGGTTACGTTACGTTGAGAGATCTGCCAGTGGAGTGGGACGCATTACATGGCCAATGCAATGCGGGCACACAGCAGAATCAAATCAACGCAACCTCAAAGGGCTTTCCTTTCCTATTTCTTGAGCGCCTTGACTTGACAAG
CGCGTGTTCTGTGGGCTGGGCCACATGTACCGTGCGTATTCTATTCGTCCTCATTAGGCCCAAGAGAAGAGATGGCCAGGGATACTAATTCAATTCGCTCCTGTCGTGTGATGCTATGCTAGTGCTACCTAAAATGGAAGAGAAGATGGGTTGCTCAAGCTGAACTGAACT
TAATGTGTCAGCCAAATGCAATGCAGTCACCGTGACTTTGGAAACGAAACAAGGGGGGCGACAATCACGCCACGCCTTAGGTGCCGGAGCCGCTACGCTACGAATAAACCCACGGTTCAGTTCAGATCCTAGAGAGTGTTGCGAGGCGAGAAAGAGTATGATTTACAC
GTCACGTATCAAAGCCCAGGGTTAAGGTAAGGCACCATGCATACCTTGGATGGGATGCGTCAAAAGCAGTGGAGGCCAGGCCTCAGTTAATTGTCATGCTCCGCTCCCTATAGTAGACTCGTACTGTACTGTTGAAATGTCCTGCTTGCGTAGCGTAGGCCCGTCATGTT
ATCGGAGCGGAGTCGTTCAGTCGCGGTGGGGTGGGGTATGCCAATAGTCATAATTTGATTTGCTTATTAGACCACTATTAGATTAGAGAACAGCTTGTTATTAACCAAACCAGTATGAAGAGCTGCCTGTGGTGTGGTCTTCCCTGTCGATTTTTGGTTTGGCGACACGCAT
CATTTATACGATACGCCACCCAATGAACGAGGTCAGGTCAACGGTACTTTGGGCGATTAAATTAAGACATTCAACTCCCAACACCACACCGAAGACGTCGTAAGATTGCCTTGCCCCCTAGCTTCTAGCCATGACATGACCCTGCCACAAATCTCCACAGCACAGTAGGAC
GATCTGATACCAAGCCAAGTCCATCCGCATAGTAACAGTACAGTTCGTGGGTCGCAGCCGTCGCGTCGCCCGCCCACAGTAAATCTCGGCTCGGCTCCGCTTTTTAGTGTATTATATTACGCTTCCGGGTTTTAATTACATTACTTGGCTACAACTCCAAATGAAATGAACC
GGCAGTATCCACTTTCATTTCAATTTATGGTAGTGTTCGTCTCGTCTTGGGGCCCGTATCAACAAACCAAACGGCACATATCTAGGCGTGATGTGATTTGGCGTTCTACACTAAGCAAAGCAGTGGCCGCACAACTGGTGTCGTGTCAAATGAGTACGTGTCAGCCGAGCC
GGGACTATTTAAAGGAGGATTGGATTTACCAGGTTTAGCTCACCCGACCCGTTTCGTCCCAACAGTGTGCGGTGCGGTTCTGCGAAAGGATAGATGAGATGACTGACAAAATACCAACAGTACAGTGGAGAAACTTCGAGGGACGTGACGTCGGGTAGCATCAGGATTC
TGTTCTGTGAATACTTCAACTGTGCGTTGCGTCGCCTCCTATGCCCATATTCTATTCATTGGCCGTACCGGAGTGCCGTGCCATTCGTTCGATCGCAGAAACGAAACCCGATATTTCTTTTAGAGGTGAGGTCAAGAGTCTCGCCACGCTAGGCTAGTTTTGTAATACTGAC
AGAAAAGAAATTGAAAGTCTGCGATGCTTGGCTTGAGGAATCGTACGTCACGCTTCGCTTAGTACGACTACGGTAACGCCACGCCTGTAACTCCTGTATACGTAACGTAACAATTACTCGAAAGTTTTAATTTAAATGGCGCCTATCGACTCTCCTCTCCATTGTCTAACCGT
CTATTCCATTCCGCAAGTTACTCCAAAGATACGATACAAGTTGTGGAAAAAATTCTATTCTATCCGCATTATTGGACAATATAATATACTCACAAAACTGCGCGGGTGGGGTGACGCGGTCTTATCACATACAATACAGCTGAACCCTAATTTCGATGCGATGATAGTTAAACACA
TGCCCTACCCTATGCACCATATAATCTTTAGTTTAGTTGCTCGCATGGCTGCAAAGTAAAGTCACGTACTTCGATTCACTAC
>6mers_2_curlcake_3
ACTACTATAGCTATTGTGTTTCTGCTCTGCTGGGGGATTTGCACTGTACTGTACTGGGACTTGTTATATTCTCCCCTCCCGAACTATCTAGTAATTGCGGTGCGGGTCTCAGGGAATTCACGGGCCGGGCGATGCGGAGGGGATGCTTTGCTTTGCCTGCAGCTGTTTTGT
GCGCTGCGCTCGGCACTAACCCTCTTTTATTTTAATCTTATCAGAGCGATAGCTTAGCTAGCATTTCGAGCACCCGTAACGTAAGCCCGTGGGACTGGTATAGTATAGTACGCTAAGATATCTGTCCAGTCCAGGCACCCCAGCGTATTACGGGACGGGTGTGCATCGATG
GTCGCTGTGCTGTGGTTGCAAGATTTCGCAAGACAAGATCTTCAAACTGTTGGCGCGCCGCGCAGTCTTGTTCTTTGCTCCCGTCCCGTCGCGGTTGTCCGAGTAATACGTAATAATACCGCTCCTCCAGGGCAGGGCCGAAGCGCATATATGGGATTGGATTTCCTGC
GGTATCACTTGCGCTGCGCTTGAGGTTGGCATTTCTACCCTACCGCGCCGGTCCTGAGCAGAATAGAATTGGGAACCTTTGGGCCAGCCCAGCCAGGACCTTGCGCCCGCCCGGCCCGGGTTACTAACAGTCTCTGTACTGTACTCCTACATGGTAATCAGACAAGAC
ATCTTTACGCGGTGACCACTTCACTTTATTTAGAACGAGTCGTTAGGTTAGGGCTCTCGCGAGGACAGGGAAGGGAAGTCGTGTCTTGATATTGGGTTGGGCTTACCGTAATGATAAATAAAATAACTGCCGATTGCGTTTTACTGTACTGTTCATCCATCTACATAAGCTAG
CTTCTCGGTGGGTTCAAGTCGCGTCGCCGAGGCCTAATACATGATCTGATCTTCCATCAGACCGAGGCTGTGCTGTGCTGGCTTCATTAGACCATTACATTATAAACGCTGCGATCTAACTGAACTGATGATTGTTTGACGAGGTGGGGTGGCGTGAGGTTGGTTGTGAA
ACGAAACGAACCAACTGCACTGCCCTTCCCTTATATCGCCCGCGTTTAGATCAGATCGTTAACTTATGGAGGAACTTAACTTTTCACGAGCATCCCATAATCTAATCGATACATACGCTACTCTATTCTATTGGCGAGGGATGAGTTACACAACACACAACGCGTCTTCTGAT
CTAGTCTAGTCCCGCGTAATGTTCAACCTAACCTAACGCCGTTCGACCCACTGAACTGATCTTCGAGGGGCTGGTCATGTCATGCAGACGTTGCGACGAAGGTGAGGTGCAGAGGACAGAGTGTCGTAACGTAAGTAAGAAGTGCCTATTCTTGTCTTGTTTAACGACTG
AGAAACGTAACGTAGGAAGGGGGGACGAAAAATCAAATCAGTGTCCCAAAGGAGTGAAGTGAAGCCTGGTACCAAACACATGTGATGTGGGGTTCCGTTGAATTTTTAATTTAACCTTAAAGAGATTAGGCACTGCACTCACTATCCTCCCTCGGGAAAGGAAACTATAAT
CGAGTTACGGAACGGAACAGACACTTCTATGAGCGATACGATAGATTACAAGAATACGAAATAAAATAGAGAATGTCCTCATCCTGCACTGCAAAGTGTTGGGCCTTGGATGTGATGTGGGATCGTTGGAGACGCTGGGCTGGCCCTGTTTTACTATCGTTCGGTTCGCCT
GCGTGCCCGTCGTCTGTTCTGTACTGTGCCCCTCTCGACATAACATATATGAAGATATAGGTGTGGAGTGGAGTGCCTATAGTATCCTCAGCTCAGCATGCCTTACCCTCCCCATCGCATCGGGTTCGAACGGTTTATATACTATACGGGTAAAAGTACTGTCAAAGCAAAG
GCGACTTCTCCGAACCAAACCAAACAGACCGGTTTCACTGAGCACAGCACTGCAATAAGAACTCAGTTTAGTTTATGCACTAGTTTGTTTCTTACCTTACGGCCAGAGACCTAGGATGGAATGGAGGGGGGCTGTGCAATAAAGAAAAGGTATCGCCAGTCCTTAAACA
AAACATGAAATTAGGGAGCTAGGGAAGGGACGGGCCGCGAAGCGTTTAATTTAATAGAAATGCGCGATGGACTAGACTAGCTCCGACCACCAAGGACCGAACCGAGACAAGGACGCAGCCCATTCCATTCTTGGCATATGGCGGAACTGAACTGAGTGATCTCTAGAAT
TCGAGACGAGACACGGTAGTTGGGACTTAGGTTAGGAGCCTGAGCTTAGAAATAACATAACTCACGTGGACGACATTAGTATAGTAACTGCTAATCCGAACTCGATTCGATATTGACCGTTAGCCTCGTCACGTCATCACACCCACCACCTTTTCTTTTCTGACATTCACCT
CACATATGCTATGCCGTTGCTAGATAAACACATCACATCCGGGCTCCTCCTTCCCCACCCCACCGCCTGCCACGAATTAGCCTAGCCTAACTAACGTGACGTATATTAAATTAAAGCTAGACGCTTCATATTTCATTTCGGAAAGCACATTTGCCAAACCAAACGACACGAGG
GTAGTCTACGTTACGTCGTCTATCATCTGGCCTAGTCTAGTGATATGTAACGCAGGAAGGAAAGGAAAATGTGCTATTACTTCAAG
>6mers_2_curlcake_4
TCAAGGATCCAGACTAATGCGTGGAGTGGACACTTAATACTAGATCGGACCGGACGGCCGCCGACACCCAGCTCAGCTCAGATGCATAACTATACCGGTACGGTATATATTAACGATCGCGCTGCGCTGCCGGACTCGAATAATACGCAACGCAAGGTCTACTTATCGAAT
TACGTTACGTCTCCTCTTGATTGGTGCAGTGCAGGGTATGCTACGGACATGGTTTGGTTTCACGGATGTCAGAATGGTTTGGTTCGACAGTGAAACAGTAACTGAACTGGTACCATGTCATCAGCCTTGCCTTGCCGGTGAATCGTATGGAAAAGAAAACTACTCGTCATG
GGTGCGATGCGATATCGGCAGCATAGGGGGAGAGGAGAAGAAGGACTACAAAATCCAATCCAATCTGCCGATCTGAAGCATATCATATGCAACCACTCGAATCCTAATCTAATTACTCTCACAAGAACCATGGCATGGGAGATTCGAAGATCCGTCTGGTCTGAGGGTCGC
TTTGTAGTACGTTACGTCAGGAAAGCCAAAGGCCATTCCATTCCATTTCAGCCGGGACGCGCCGCGCTTAAGCATTAGAAGATCCTGTCCTGACTAGGTAAAAAACCACACCACACCAGTTAGCAGATAGTGACCGCACCGCCGCTCTTATACAACGTTAACTTAACGGA
GCACGACAGCTAACACTTTAATACACTATCTTCTGGGTGCTCTGCTCCTAGGACTGCGACTGTAGCTTAGCTGTATCCGATACCCACGCTCTGCTCTTCTGGTGCCGTCAGCGACATGACATAGTTAAGAGCATCTTATTAGATTAGCTGTAACCTTACGTAACAAGACAAG
AGTCATTTTCGCACGACGGCACGGCAAATTAGATCTTAGCCCTCCCCTCCGCCACTCCACTTCGCGTGCAGTGCATGGCTTATAGCGAGTAGATAAGATAGTCCCTTACAAGTACTACGGTACGGACCCTTCGCCTGACGCTCAACTCAAAAACGTACAGGATCCCACCT
CACCTGAATCTCTCATTTCATACCCTACCCCTTCCGTGACGGTTCATAGAATAGAGTAATCCTGCTATGAAAGGCAAGGCAATCAGTGCAGGGTTCTCTACTCTACGCATCAAGGCTCAAATCGAATCGACGAGCAAGGTGAGAATCGACTCGACGCCAGCCAATACATAC
CGCACCGCAGACACAATGAGGACAAAATAAAATAAACAACCGAAGACTCTATAATATAAAGCACAATATATGTCCCGTACCGTAGTGCTCTCAGACTGTGAATGGAATGATTCCGCGGACAACCTACGCTACGCCATGTCTAGCAACAAAGTGGAGTGGTTCTCACGTGGC
AAAGAAGCGAAGCGATGGCACAGTTGGTAGACGAGACGGTCTACCTCAACGGTGCTCTGCTCTCGCTGTGATCAATCGTCAATTCAATGTGTGAGAGTCAGGGCGAACCGAACGGCTATATTTTCAGCAGTGCAGTGCACAGGCTGACATAACCCTTTCCTTTGGTGAT
GAATGCCGCACACTACACTGGCGGCTCAGCATATCATTCCATTCTAAATTAAGCTCCGTAGTCCAGTCCCCTAAGTGGCTGCGTGTAGTGTAGTGTATCCCAGAAGGACGTGTGGTGTGTCCTCTTCCAATTATTTAAATTAAAACGTGCCACAATCACGCTCTGCTCTCCT
TGGTCCGTGTGATATCCTATCCCGAGTGGTTAAGTCTAATGGAATGGAAGCTCGTAGCGGCATGTCGTGTCGTCGTAAAGATTTCCGAAACTAAACTGTCTCGCGGCTAACGGGTACGGTACAACCGCTGTCATAGGTAACCTAACCATCTTGATAAGCAAAAGCAAAGCA
ATGACAATCTGTAGGATACCTTACCTTCCAGGGGCGCTACAGGATAGGATATAGAAATTTATCATGAAGGAAAGGAAGAAAGATAACGGCCTACTTTACTTCCATATGGGAACTATGAAGTGAAGTGATACGGCAAAAATGGACCCGACCCCAGCTATAAGCTCTCTTAATTT
AATGACGTCTGATAGCTGGACCGGACCGCCGCGAGCCTACCCCGACATGACATCATCCGTAGAACATTGACGAGACGAGCCGGTCACGTTACAAATTTAATTTGTAATAGGGCTATCGAGTAGAGTAGCCCGGGATCATACGATTTGGTTTGGTTATAGACACTTTATCATG
TCATGTTTGCAGGCTGGATGTATGGGATGGGCAAGTCCGGACCGTATAAGTTAAGTGTTCGGTCGGCTGGCGAATCGAATCTCGCCCCGCAGGAGTAGCCGAGCCGACAGCCGCGGACGTAGTTTAGTTTATTCACTGAATGATCGGCCCAGCCCACAGAGCCAAGAC
TCCCGATCCGATCTCCGTAGCAGCTCATTGAATTGAATAGACAGTCGGTGTTAGGGACGGGACATCTAAGAGGAGGGCAAGCCAAGCCAGGCCAACCCGAACGATCGTATCGTGGCTACACCTTGACTAAGGCAAGGCCACTTAGAGCTTGTGCGGTGCGGTGTTTAC
TCAGGCTATAAATCTAATCTCAATAACCAATCGGTGCATTGCATTCATGGTTCATCCACATTTGGTTTGGCCCAATCACCACGAAGCCCC
>6mers_2_curlcake_5

GCCCCCGGGGAATGGTCCTAGTTAGGTTAGTAGGTTCCGTGCTGGTGGTGTGGTGCCTGGACAAACAGCCTTAAGTTAAGCGAGTGAATATTCCGGTGCGGTGCGTCCGCGGAGAGACTTTAAGCTAAGCGGTGCGTTACGATGTTCACGTCACGCTGGAGGTCTTAG
TCATTTAATTTAAGTAATGACTGGCTTTGTTATGTTAAGGTAGGTCGCCGTATTGTATTGTATGCGAGACAATGTGGCAGAACAGAAGGGTCTAGACTTTTAGGAGGGGAGGCCTAGTGCGGTAGTTGACTCGACTCTAGGTGGGCGGACCTTTTGGTTTGGTTGTTCGAT
CACACTCACCTCACCTCTGAGGCGGTAATTTCCAAGCCAAGGATGTGAACATATACTGCATTGCATGCTGGTCCCCCTCCGATGACATGACCTGCGGGCGAAGTGAACTTTACTTTAAATGAGTCCTTGTCACAAAACAAACTTAATATTATTGATAGCATAGCAAAAAAGC
CTAAGGGTAAAAGAAAAGGCTTTTCGTTTTCCACGGGACGGGAGTAAGTTTTTAGGCTAAAGCAAAGCCTGGACCCCCCTGTCGATAGGATAGTGCTAAGTACCTCGTTATCTTATCTTGTGAGGGCCCCTGATTCCCTTCCCATGATGTAAAGTGCGCACGCCACGCCG
GCATACGATACTCTTACTTTACTGGGTCATATTGAACTCTTCGCTTCGTTTAACAACCTGAAGCTGGCCTGGCATCCTCCTATATGTGAAAATAAAATGAATTATCTGATGTCTGCATTGCATCTCCAGTGACAGCAATGTTGTGTTGTCATATCAGGTACATTAACCTAACCAC
ACGCGGGGGCAACGTTGTGTTGTTAGGTGGTAATCTACGTCCGGTCCGACCAGCCTTATTCTCGTGGTGTGGTCACACCCAAGTTTGTCGATCAGTAGCGATCGCTCGTATATCCGAGCCGAGCAGCAGACTATGCCTAGATTAGATTCAGTTTTTTGCCTCACTTGGCT
TGGTACGCTCATTCCATCGTAACGTAACTAGAGCGAATATAGCAATTGAATTGTGCAAACCATCCATGGATCCGATCCTTGCAGTACTTGAATAGAAGAGAAGCGAATATGTCTGCCACCCGTCCCGTGCTAATTCAAAATGAAAACGAAACGGCAGGGCTTGTAGGGGTAC
AGTACACCTCGAGGCGGTTAGATATGATATGCTCGAAGTTCACACTTGCGTTGCGTCCACGAACTCCCCTATGCTGTGCTGAGTGTGACAAAGCGTACGAAACGAATCAGGAGCAAAGACCGCATGGCATGACCAAGCATGAGTATTTATTTTATGGATACTCTTCTTTAC
AGTCCAGTCTCATAAAACCTCCGTTTCCGTTCCGCCTCAAAGCACCACGACTCAACTCAGTCAGATGACAGTCTTTGTTTTGTTCTTCGGCGTTGGCACCCTCACCTCATGTGCCCTGGCCCTTTTTTCTTTTCGCGACGGGAAAGCGTTTGGATTGGATTGGACTGCCC
TAGGAACGCAACGCCTCTGTGTGCAAGACTACCGTACCGGTGCAGATTCTTATGAGCAAAGCAAGACGCACGGTATGGTTTGATTTGATTAGGGGTCCAGGGATACAGAACAGAGTCAGCGACGTACCGATCGAATCGAGGTGAACTAGTCCACGTTTTGTTTTGGGGC
GGAGAAGCATGTATAGTATATCTCGGGCTGTAAGCAAATGAAATGTTAGCCCTAAGCCTATTATCTTATCAAAGAGAACGTGCACATCCCATCCCCCGCAGTTCACCCTGACCGTACCGTGTTAAGCCGAATGATGAACAGAACAAGAGGGCGTGAACCAGATTAGATTAAT
TGTAAATTCTCGTGCGGCGCGGCCTTGAGTAGCAACGTCTGTCCTGTCAGCTTGACCGACCGCCAGACCAGAGGCAATCGATCAGCGAACATAACATGGGGAGCATATAATTAGTGGAGTGGTGAAACCTGTTACATAACTCAACTCGGATGTAAAATTCAGGCCACGCC
ACCGTAATGGAACCTTGTCATATCATAGAATAGGCGCATCCACAAGTCAAGTGTGTGCCTAGATCATGTATTGTATTCTGCCATTGAGATGCGCACGGCACGCCGAACATGTTCCAAGCAATGCAATATCCAGCGCTTAGAATGTACTGTACATCATATTGGTCTTAACGAAA
CGAAATAAAAGTGAGCGTCAATTTAATTTCAGGTCAGAACTGTGTCAACTCAACCACTCATGACGTCCCGCATAGCATACCTTACTTGTTTTATGTCCCGTCCCACTGTTGTACGGTAAGTGATGTGATTTACGGTGGGGCTTGGGAATGGAATGTATTGTGACACCTATTTT
TTTTTTGCTTTCGTAGCGCGCCGAAGCGAAGTCACCCTAGCCACATTACACTACACTATTCCCCGCCTCAGAAGAAAAGAATGCGCCGGCGTGAGTTACTATACTATCACCCTCACAAGCCGCTCAGCTCATCTGTATCTGTTACGCAACACAACAGGAGTACGTTGAGGG
TAGCTTAGCTTGACGTGAAACATCTAATCGAATCGCAAATTTTACCAGCAGTACAGTACAACAACCAATTAATTCCACGT
>6mers_2_curlcake_6
CACGTGATTACATGCCCCCCAAAACAAAACAATAGGGTCATTTAGGTCGTGTCGTTCTTTGAGGTTCTAGGCTGAGCTGACGACTACTGCGCTTGGTATGGTATGATAAAGAAAGAGGTGATTACATTACCTCAACTCTGATAGCGAAGGGAAGGTAGGAATTGAGCTCTG
GTTGGGTTGCCAAGTGTCCGCTTTTGTTATGTTAATCATAAATGCAGGTGGATTGGATTGAAACGATAAGGGCATCAATTCAATCCACCAAGCAGCGGCTCTAATCTAAACGGGGCTGAATTTAGTGAGGTGAGGCTCAGGAACCGAATTTATTTTATTCATGCACGCGTTA
GCACTCCACTCCCGGGCCTCCAGGTGCACGCAACGCACCTCGATGTGGATAGACAGAACAGATACCAAAATTCTATACTAATCTAATGGTCTCAATCTGCAGATGAGATGAGACATAAGCTGCCGGATCACCTCACCCCGGCCCCACACCTGTATGCTATGCCGACGATG
CGACCTATACTGTACTGACCATCAAATATCTATAGCATAGCATCGGTTCCTACAGGTCGACACGACATGCGTGGCATCGACGGGAGCGGAGCTGATCATAATGTTGTTGGGATGGGAAGCTCGGAAGTTCTAACCGGACCGGACTCCTCGATTACCCTTCGTTTCGTACTG
TCCCAATGCACAGCCCAGCCCCTTGTATTAATCCCCAAATTAAATTGGGAGGTTATCTTGCATGAGATGAGGGGTGGAAGAGGGGAATACAATACAATTTTGAAGGCGATAACCGCACCGCACATAGGAACGGGAGGATATGATATGACGGAGCAGCGGCGAGTTCAGTT
CGGGGCATTGTGGTAGGGTGCGGTGCTTGTCCAATTTGGTTATCCTATCCTAATAAGTTGAACGCAATTGGATTGGCACAGAAAATTGCAAGTATAGTATAAACTTCTAATCCGCAACGGTACGGTCAGGCAGGGTTGTACGAGTCAACCCGACAATTAGAGTCATCTCATA
TCTCGAGATCGCTATTAGTCTAACCTAACTCAAGATTGACACGCCTACACTACAAACAAGCGGCGGTCCCTCAGCTCAGTACAAACCCCTTCCTCTACACTACAGATTAAGATGAATTACCATTCCATTAACTAAACTGTATGCATTACATTACCTCTCCCACTCCGGTCATCG
CATCGCAGTTTGTGTTACACAATGTAATGTCTCAATATAGGACCATGTAATGTAATTATGGTATAGGCAAACGGTACGGTTTTGGCTCCAGCTCTATGGAATGGAGCACTAGTACCTGCAACGCAACGCACCCCAGTTGTGGTACAGACCAGACCCAGTAGCGTAATGCATT
CCTTTCCTTAGGTTGCCTCGCGAATGACATGACATCTCATACAAATTCCAAGACAAGAAACCGCGGATCACTTAAGGTAAGGTTCGCCATACTACTATATCAAATCAATGAAGGAGGATGCCATCTTTTCTTTGCTTCCTCGGATTATGCTCCGCTCCAATGACTAAAGGTAC
GTGACGTGACTCGTCATGGTGTGGATTCACTTCACATGTACAGGTCTAGACCTTCCCTTCTATCTGTTAATGGTGTAATTTAATTATCTCTGTGATCACCAATACAATACCTGTGTGTCCACTCAACATTACATTAAATATTTGCACTGGATCAAATCAACACCAGCGGGACAGT
TTACCTTACCGTCTGTTCTGTAGAAAAGTAAAGTAGGGCCATAGCGGTAAAGGGAAGGGAGGTCAATTGCATGCTACGAAACGAATGGGGACCAGAGCTTTAATCTAATCTAGATGTGAGCCCTACGTGCCGTGCCACGGATAGACTCTCTCCATTCCATTGTTGCGTTCT
GGCCTTTGCACGTTTTGCGGAGGATAAGACCCCCACCCCTCCGTCACATCCATGGACTGGACTGAAATCCTTACCATATTAAAATAAAAATACGATTATGTCTCTGTCCTGTCCATGCGATCCCTTGATGCCTAGCCTAAAAACATGAATATCCAACGCAACGCCCAAACCA
GTGGCCCGACCCGACCCTGCATAGCCTGAAATACCGTACCGTTGGATCGCAAATACACTGCGCTGCGGCGCCCCAACCATAACGTTCCGTTCCGTTAAAATCAAAACCCCAGCCCAGTACATGTGCTACCAGAACTCAACTCTCGTCTTAACAGCTGATTGCATTGCTAT
CACGATACGTTCGCAACGCAACTAGGTCCGCGTCAACCAACACAACAAATCACGATCTCCCCGATTAGATTAGCTCCTTCTAGCCATTGTGCTGTGCTGCCTGATCTACCTACCTGGCCTGGGTAGCTTACAGTTCCGACCTGACCTCCCTATGTAGCGAGATATTCTATTC
GGCTAAATTAGTGAGGCAGGGCAGGAAGTATCTCGGTATTGCATCGCATCTCAACACGGCCGCCCTGGAGTGGAGCCTTTATCCCTTTGAAATTTAATTTGTGCACCTGTCTACTCCCCTCCCCTGTTCCACCTGCCCGGCACGACACGAGAGTTTGGGTCCGGCTTCA
CTTCACTAAGCCGACTCATAAAGCGCAGCGCATTCCTGATATGGCCGGATGGGATGGAGGGAGATGCAGTTATTAT
>6mers_2_curlcake_7
ATTATAACACGGACAGATCAACGACACGACGGCGCGGGATCGAACAGACCAGACCTGGTCCATGTTAGATTTATATTATACGCCGTTACAGCGATCCTGACCTGATGTCGCGCGTTACCTGCTCAGCTCATCGGTTGAAGGCTAGGCTAGGCTAGGGCAACGCTAGAAAC
CGAGACGAGAAAAAAAGAAATGTCCGAGAAGAGAATTCCTAAATGCTTTCCGACGCGACGTTTCGATGAGGACTCAGCTAAGCTACTTACATTTGATCCTGGTCCGGTCCAGTCCAACGTTCACCTTAAATTAAAAAATTAATGCTCTACTTATGTTATGTTTTTGAGGCATT
CGCGCGGCGCGGCCGGGTTATTTTAGCACCTCACCTCTGAAGATGTCACTCTGTCCGCGTCCGGGGTGAGCTTTAGGAGGATTAGGATTCTTGCGGAAGTCTTTTTATTTTACTCATAGGACATGGATAGAATAGAAGTGGAAGGGTGGCTGAGACTAGACTTCAGTATG
TTCGGGCAGTGCAGTGCGAGGAGAGCCGATCATGGACTGGACGTTACCCTACTGGAGACAACACAACGAAGGCGGGTGCGGGCTGTCCTGTCTCGGGATACTCGGAAATCGGATCGGTTAGCGTGTATCATTCGACTCGACTGTCTACAATCTTTTCTGTCGTGTCGT
CTCTGAGAGGTCGGGTCAAGTCAACTTGCCGCACTAAAGACCCGACCCGTAGGGAACAACATTGGATAAGATAATATTCGGACGTGCGCTAGAATAGAATATCAGGTCCCCCGGCTCCGCTCCGGATAATTCGCTTGCTTTGTGTTGTGTCAGGTTTGTGAGTATTACGTT
ACGGGAAGCGTGGGGTTGTCTTGTCTTGAACAGCCTCGACCAGAAAAGAAAAGTCTGGAGAGCACGAGCTGCGCTGCGTCGGCCCCTCCTGTGACCTGACCTGACGTTAATGTAGCTTATCGAATCGAAGTATTACAGCCTCGTTGCCTTGCCGTGGGCCCTTGCTGA
GTTATGTTATCCAAGTCGTGCCATCTTCGTTTCGTGGTTCTTAGCGGGAAGTTCCGTTCCCGGTAACGATTCCAGCTGTTCTGTTCTAGGAGAGGCTTTTGAGTGGAGTGCACAAGCCGCCGGGGGAGTCGAGTCTGCAATCATACGAGGAGTTCAGTTCATGTGACAC
TCTTGGACCCAACCCAAGGGCCCCTCATTCTGGATTGGATTTCCCTGCCCAGGAGCGTGGCGTGGCTCTACGACAGAACCTCGCCGCGCCGATCCCCCCTACGAAGGTTAGGTTAGAGTGGAATACCCCGACGCGCCGCGCCCGCTGAGAAGGTCATTATTTTATTGC
CTCTTTTTCTGAAGTAGTGTAGTCGACCAACTTAGCGGTGTATTGTATACCATAGATTGAGCGCTCGCCTCGCACCTTCACTACACCCCCAGACCAGATAACATGACTCTAAAATCGTATCGTATCTGGGCACGGCGGTCACATCACAGGTGAAGCAGGCATGATTATATTAT
AGACAAGAAGACATTGATCGGATCGAAACTCATATCTATGGATCTGATCTATCCCTACGTACAGTAAGGCAAGGCCGATGTTCAAACGTAGCAACGCAACCCGATGCCTTTACGCCCCGACCCGATATACACTACGGCCAATCCTATCCTACTGAGTTGACTGATTTGAATT
GAATCTGCGTGGGAGTAACGTCCCGTCCAGATTCGCCCTCGCATAGTGTAGTGCGTATATAAGGCTCGAGACTAGACTGTCAAGCCGTATCGTCGACACGACAGGAGGTTTGCGGAGAGCTACGCTACGCGCGGCCCGAATGGATGGAATGGAAATCGCCTTCAGATAC
GGAACGGAATCCGCCTTCATCTACCACGGCACGGTGATAAAAGGCACAACGGCTCGGCTGCGAACTGATTTTGAAAGTTAAGTTCCAGAGCGCCCAGCACCCCGCCCCGTCAGTGTCTCCGCTTCGTTGCGTTGCCCCCGAGCATGGCCTGGAATGGAACGAGTCCT
ATTTGTACGCTGCGCTGCACGTTAGACGCCCATTACGTTACGCCTACCGCAACTCGCTAAACTAAACTCGTAGGGGCCATGACTTCGCTTCGCCAAGCTAAATGTAGGTATGGTATGACGAACGGAAGCATTGTCCCGTCCCTCTTCACAGGTAGCCACACGACACGTCA
GAGATAGCCGAGCGTCACGTCACTAGGTCTCCGGAGCGAGCGGAGCGACGGTCTCAGCGCAATGGGCTGGGCAGTCTTTAGTTCCAGCATCTCATCTGACGGTGATTAGTATTCTTCTCTTCATTGCCCAATTCCTAGAAGGGAAGGGATGCAAGATAGTTTAGACGAG
ACGGCTATTCAAGACTGGTTAACTTAACACTCCGAGGCCTGTCCTAAGCTAGCTTTCGCTGCACCAACCGCGCCGCGCAGACGAGATCAAGGTGTCCTGTCCAAATCAAATAGCACGGCGTCGCGTCTCTCTTAACCGTAACCTCTCCTCTCATCATTAGCGCCCACTAC
AGTACAGGTTCCTCTGCAGCACCTAGACTAGAGCAGGCGAGGATTTTACCTA
>6mers_2_curlcake_8
ACCTAGGCGAGGAGACCATGATCGTATCGTGCCGAAAGAATAGCATTTAATTTAATATGATGTTACTGCCGCAGGGCAGGTACTCGCGTGAGATAGGATCGGATCGGGCATGCTAATAACGCTTAGCTTACTTGCAACTTGGGTACCACCCCACCATTTGTTGGAACGCCAT
CCGATCCGGAATAATCGCGCTCGCACAGCACAGAGGCATACTAAGCGGAAAAGAAAAGCTCCACCACTAGGGATGAACTGAACCTGGCTATGTTTCATTGGGGTGGGGCGTAACTTCCGGTTTTTTAATTTAACTTTGGTGCGCGTAGATGGCATGGCGTCGGATCAGTC
ACGGAGGTGAGGTGGTGTACTATGCTGTTCAGGTCAGGGGAACGTGCTCGGGAGTGGAGTGGAGCGGCTTATCGGATGATTCGATTCGGCAATTTACTGCGTAAGGATAGGATGGTGCTGGGTGTTTGTCACTTCACTGACCACCGGGCACCGCTGAGCTGAGGTGA
CGCATGGGTAAGGCATGGCATTCTGAATCTAAAACTCCATACCATACGGCTGCTTCGTGCGACTAGACTAGGTATTGTAACACTGTAAAATAAAATAAGAGCGTATGTGGCTAATTTAATTTTCCCATATCACCGCTTGGATTGGAAACGACCACGTGTCTTGCATTGCATTAA
AGTCTAAGTGGCAATCCAATCCAGTCACTGCCTTGCTGTTTGGTTTGAGCCCTCGGTTGCCAGCGCGGCGCGAAGACAGGAATGCCAGTAAAGTAAATCACTCGGGGTCCAATAGTCTAGTCGTAGTGGGCACGTAAATAGCATAGCTCTGCGCCTCTCACCAGAAGAG
AAGCTACCAGCCCGCCCTCGCTCCGCTCTGCATCTCTTGCCCGGACTTGACTTGGCTCACGCACTTCCCACACCACACGTTTATCCGCGCTACGCAAAGCAAAGGTCGTTTCCACAACATCGTTGTCTATCGTGAATGTCGTGTTTACTTTACATTGCGCAATATCGTTGA
GTTATTATGAGTCTGGCTGTGGCGCTCCGCTCCCCTGAGGAAATGGGTTCAATTCAAACCGGCTTGGTAACCTGAGGTGAGGGTGTAGCCGTGATCAGCAAGGCAAGTGCCGTATACAGTCGATAAAATAAATAGGATCTTGCCGCCAAAAGAAAAGAGTTTCGCATGTC
GCATGAAATGAAAAAGACAGTATAGGCGCAAGGCAAGCGCCTGGCATCGCCCCAAGACAAGAGCCCTGCCTTCGGTGAGAAGAGAAGAACCGTTTGGAGTTATAGAGTAGAGGGTGACTATATGGGGTTGACTTGACGATGTAATTGGAGTCGGCCGGGCCGGTACCC
AATGCGTTATATTTTATTTGACCAATAACAGGCCGCTATGCTATTTAAATCCGTTCAATGGGCGGGGCGCTATCGAAAGTAAGACCGAACCGAATAGGGTAAACGTCACCGTTCCGTTCCGAAAACCGTTCCTCAGAACAGAACACTCGCATTTTGCACTTTTGTTTTGAGAT
TTAGGGGAAGAACGGGACGGGCTACTCAAACGGGGGTTTAGTTTAGCCTCCGGTTACGACTTACGATACGACGTCCGTTATGCGCGCTAGTCTAGTAGAAGCCATAAGGGTGAAGTGAAGTAGTCTGACTGAAAGGGAAATGAAATCCTCATCGTGGACCAAGGCAAGG
CAAACCTAGCAGTTGGGTGTTCTGTTCCTACCAAAGTTTCCCAACCGAACCGGGGTGAGTGGTCAGATTTTTTTTTTTGCTCGACGACCCTAGCTTTTCCCGGCTTTAGTCCTTGGCGCAGCGCAGCCAACGAGTGACATCAACACGACACGCAGGCGATTGTACCCGG
ATCGGATCTGGGCTCGCCGCAAACTGATCTGATACGTATGGCCTGCGGTTTTCTTTTCCTTGACCCCGGCAGTAGTATAGTATTTCGTAAGCGACCGCGCGTAGCGTACATCACTTCGAACTACGGCGCGGCGATGGCTTCTTAAGTACTAGACTAGACTTAGACTTCACA
GTGCAGGGCAGGCAGGGTTTAGCAGATCTGGTCTGGTTTCACGTCACATCCCGACTTGACTTGAACGTAGATGTGAGCTAATCTAATCTTTCAAGTTGTTTGAACCGAACCGAGGTAAGTTATGCGGCATGCCATGCGGAAGGGGTTGATCCATAATATAATGCTGATGTA
GCTGGATTGCCTTGCCATGCCGAGTCCGTACGCGAGGCGAGCCTCCACAAATACGTTTAAGTTAAGATGACATACTTCAGCTCGCTTCGCTTCAGTGCTTGCTTCTTGTCGTGTCGGCGACTTACAATTAAATTGTATTGTTGACACCATTATGCAGTCCGGTCCGGGAGG
CGTCCATGAGATCCGATCCGTCCATACTCTAAGAGGCGCGGCGCGGCAAGCTGAATTAAGGAGACGAGACCAGTCAGGGAGACTCCGTACCGTACTACCGTGTCGCTGATGGCAGGGCAGCGTTTGCGAAATTGGGGGGTGGGGTCTGTTATGATAGGGCGTGTCGT
GTCTGTGCTTGCCGGTTCGTACAGTACAATTCATGCGCTGACTTGAAT
>6mers_2_curlcake_9
TGAATGGCGGCCTTTCTCCTTCTTCTCTTCAAATGCCAGATGGTGATGGGATGGGGGTACCGACGCATCCTTTGATTTGACACACACCGGGTTATTGCTATGCTACTACACGATTTGATTGTCTGGTCTGGAAGAGTTGCACGTCCTATCCTATCAATTGCGAATCCCAGGT
TACGTTACCAGGACCATCCCTAGCCGGACCGGAGATGCAATGCCTGTTTACGTTACGTTCACAAGTCATTGGTCGATACGATAGGTTGAAGTTCCACTTGTAGTGTAGAGACCGGCTGCTAGCGAAGAGAAGAGACGTTGTTGGTCGACCCAGCCCAGCGTTAGGCTTT
CGTTAAAGGAAAGGATTGGCCAGAGGTATCGGAACGGAAGGCTGGCCCAAAACTTACTTTACTTTGCGAGATAACCCTTATGTAATGTACACGTTTGCGCCGCCTAAAATAAAACAGGGCGCGAACCGGGTGGAGTGGACACATTATCTATGAATCGGGTCGGGGGTGG
AATGTATTGCATTGCATTGCGTTCCCTAAACCAGTAAAGCAAAGCCACTGTGGGATCTCCTTTTATTTTATAATAATGCCGCGATTGTTCTGTTCCTTTCTCATGAAAAGGTCTTGTCTTCCGCTGTCGGTCGGGTGTAATGTAACGTGTGGCCGGCGACAAGGGAAGGGGT
ACGGGGCAGCGCTGGGAAGGGAACCCGACAACTAACCCCCATCCCATCACTTCCCTTAGCGCAAAAGGAAAGGGACATTTCAGGCCAAAGTTTAGTTTCCAGACGACGCCCCACGCAGCGCAGAGAGCAAGGACGGAACTGGGCTGGGGACATGAGGAAAGCGGTA
TAGTATAGTGTCGCTGACAACGGAATTGAATTTTGTCCGCCATCAATGTTGCGTTGCTATCTTACGAGCTCTTAGATTAGATACCCGAGGGACCAGGTAGGCTAGGCGGGCTAGAGACGGGTAACCAAACCATTCGCTAAGGGTCGGCCGACCCGACGCTGGTGACTTTA
ACATTGAATTGATGGGCGGTTACTAATGTGCCGTGCCAATCGCAGAGTTCTCAGGTTAGGTTTCTACGTGAACACGTCGCACCGCACCGCGCCTACTCCTGTGTAAGGTAAGACGCGACAAAGAATCTTAGTTTAGTTCGGGGTCAATCGGCGTAGAGTAGACCCGCGT
CCCCATGCCAAGCAGATTCAAGCTGTCTAGTCGGGCGCGGCGCACACGAACCCATGCAACAGGACAGGCATAAGCACGGGCATCACAGCACAGGGCAGAAAATCTAACAGGCGAGAGGCGGGTCTGAGCAGGTGCTTGTGCTTAGGTTTGGGTCCCGCAGTCCAGT
CCCGGAAACAGCACATGCTCTTACCCTCTTGTGATGCATACACAAAGCAAAGCGTGCGACTCGAGGCTCTGGTCTGGCACCAGCTGCTCACACTGACCTGACTGGTTCGCTAACTCACTGGACTGGACGCGCGACGAAATATGACGCGACGCCCTCAAGCACTCCGG
CCAGGCCAGGCCTCGACTATTGCAGCTCAACTCAACTCGTGACTCGGCGCAATTCAATTCGAACGTTCATGGGAGAAAGTAAAGTAGGCCGGCGCTTTTTCCGTGCCGTGCATGGCAAGCAGCAATAAAGTAAAGTCCTGCTGCGGGTCCTCGGTTCGGTAGCGCACT
CTTTCAGCGGGGCGGGGCCCTGGGAGTCAATACTTAGCTTAGTAAGCCTTTATTTGGAGCGTAGCGTCTCTTTCATAGTTCTCCATACCATAGGTAAGGGCCTCTCCTGAACTGAACACTTTCGGACCGTCCTTCGGTTCGGCTACAGCTTCCGGTACGAGGCGAGGTAA
TAGCGGATTATCCACCTCACCTACTTGCTGTTGTCCTACGCGACGCGACCTCACTCTGGCAGAGTCCAGTCCGCGAGTCTCCCAGAACGTCTTTCGTCTATCGTCCCTGGGGTGGCCTGGCCTGCGCATTGCCCACCTAAGTGCTAAGTCAAGGCGCCACTGGGGGAC
AGAGGGGGCACGCGCACGGAGGCGAGGCGTCTCGTAGTTTCGACAACATAACATACATCGAGAGTATCCCGCAACGCAATGCAACCCCTTTCTCGTGAGGTGAGTCGAAGTCGCGGAATCATCGTTACACATCGGCATGTTGATTGGCCTGGCCGTGGACTTCTCCCT
GTATGGCCCTGGTTTCTTCGGCACGTATTTCATTTCCCCGTCTTCACCGACTGCTATGCTAGGAGATGGTCGTGATGGACGGGACGCTCCCACCGACACCTATCAGATTAAATCAGCTTAAAAAGTCACTGCCAAATCCAACGATGCCCGTTTCTCTTCTCACGGTCGGTG
TCGGACACTCCACTCGGTCTGGGCTAGCTAAAGGGAAGGGGGCCTATCTCAGGATGCTGAGCTGAAAGGTGCCCCAACTGAGTCCAGTCCACTGTAGATACGGATCGTCCTTTGTGCGCGTCATTCGTCCCCAATGTCCCAGGCGTCTAATAATAGTCAGGTCAGCCA
GTGTGCGCAGTTTTCCTTTCCTCGCTCACTCGCCTTCAGAGAAGAGAATCGTCGGTCTAGAACACAC
>6mers_2_curlcake_10
CACACGGGTATTTTATCGCGAGATCAGATCAGTCGCAGGTCCGTCAGTTGAGTTGCAGCACGTCGCCGGCCCCGCCCCGCTAACCGTCTATTCGACCCGTCCCGTTGTCTGACCCGGTGGAGAAGAGAAGTCCCCGGTCCTCGGCGTTGGGTTGGTTGGAGTACTCA
AGTAAACAAAACACAGCAAATATGCGAAAGACCAGACCTTGTCGCACTTATAAGAATGGAATGAGTGACCCCATAGCGTCCTGTCCTGGAACCCATGTCACTCTGCTTTGCTTTGTACCCAGACTGGCTGACTTGACTCAATCCCTTTTATTATTGTCTTGTCGACGTGTAG
TCTATTGAGCCGAGCCATAGATGTCACCAGGATCCGATCCGAGTACATTCGTGTTGATGGTATGGTCAGTGAAGTTGCGCGCATAGCATAGCTCCGCATCCTAACATAAAGTAAAGTATAATTCTATGGCAGCCCGGCCCGAGCCATTCAGTATACAGATGAGATGCTTACC
CCCCATGGTACTTTGTTACTTTGACTTTCCGGCGCTGGACTGGAATCCCGCACAGCACACGGCTTTCCCGGCGGGCCTGTAAGCGACCGGACCGGCTTAATCGCGCAGCTTCATTTCATCGCTTCCTTTCCATCGCGTAGGCGTACACTGCTAGTTGTGTTCAGTGTTC
GGAGCCAGGTACACGCGTGTATACGCGTGTGGATGGGATTCAACGCCAACGTCTTCCTTATGAGGCTCTGTATTCTCTGAGCTAGGCTTCGAAGCGAAGCCAGTTAGTTCAGACAGCAACGCAACACCGTGAAGCCCATTTGCCTTGCCTTAGTCAAACTTCCCCGCGG
TGCGGTCCCTCTATAAGTTAGTGTGTTGTGTAGTATCGCTCGAATCCGTCGCGTCGAACCCGGGCCGAGAGAGGACAGGACACGGCAACTGGCGTAAACTAAACTATGTCGGAGTCGGGTCTGCTGAACTTTGCTGTCAGTCCGTGTACGGTACGGTTGAGTTGTATGT
GTACAGAACAGAGGTACGGTCCACGTGGTGGCAGTGGCGTTCTTACAGCGAGCCTTAAAGCCTAGGGTCGTTTTGCCAGCCACAGCCCACGGCGGATTCTCAATAGTATAGTGGGTCACGGGCGTAGAGTTTAGTTTATAGGCAGGTGTCGAAATCGAATCGGGATGAA
CGCACCGTCGAGCCGAGCGGTCGAGCGGATATCCTTCCCCTTCAATGGCGGCTCGTTTGACTAGACTAATCATCCGGAGATTGTAAGTCTATTAGGCAGTCGATAAGTCGCGAAAGTCCAGTTCGTATTCCCATTACTTTACTAGCCTGACGGATGACCCGCGCTACCCG
CGTTGCGCTAACACACAACTTGCGGCCTCACCGGTACTCAACATTGTACTCGCGTACCACAGGCTTGCTTTAACTTGCAATCTAGTGGGCGGGAATAGCGGGGCCAACAGACGGTGGTGTGGTCGTCGAGCGGATAATGGGTAGGGTAGATTCCGACTAACGTACGAG
ATTTACGAGAGGGAGCTTTATGGCTTGGCTCGTTGTAGGTGTCTTTCTAATCTAAGGAATAAGGAGCTTCGCAACGCAAGGGCTCAAGTATGTAGTAGTGTAGTTCAGAGATATAGATCCAGACCAGACCCTCTGGGCCGTGATCGATGTCGATGGGCTCGGAGTTTGCA
CCCTCAGCACCGGACAGCGCCATGGAGTGACCAGAGTGCTCGGCCTTCGCGTCGCGTCGAATTGCTCATGATACTTTTGTTTTGTCGGTACTAGTTAACAATGGAATGGTTAGAACGCGGGACTTGTGGTGTGGGTTTGGCAAGCGCTCAGAACTCAGACAATACTGGA
GCTCGATTGAGATTGAGTACCGGAATAAGGTTGTAAAGGTATTCGGTAGAACCAGGCTAATGCTAAGACATGCGGCGAGCTATTAGAGCTCGTCAACCGACTTAACTGCTCTGCTGCACGGAAAGCTATGACAGGACAGGGGAGGGCGGACGGACTCAGCTCAGCAGG
ACTGTTTTCAAGCCCCGGAGGGCCCCGTGTTACCGATAGGCGCATAGGCCGTCCAAAGAACGAGATTATAACCTATCCTCTAGCATGTGCCTCACTGTGCCTCCCTCGTGCTTGCTGTGCTTAAGTTGGTATCAGGGGCGTAGGCGTTGATATAAACTGGCTATGTGGGG
GGAGCGCGCCGTACGGGGAGAGCAGTAACCCCCCACAAGCATTTCCATGAAAGGGGCCGTCCGGGAGTACATAACTGCCCGGGCCCTCACTATGCCCCTTACCATGTAGTAGTTTCGGGCCGTGCCGTTGGGTGATTGATCAAAGGTTGTTGTTATTACGGGTGCCGG
GCGAGTAGATTGCGAAACGGTGCCGAGATTTTTTGGACTAGTGTAGCTTACTC

### Table S2. Sequence backbones of synthesized RNA oligos.

>5mers_1_curlcake_1
ATCATAATTAATACGACTCACTATAGGGAGATCCGGCTGTGGAAGTCGATGAATCAAAAACCTTGGACCCTATCTCACAACGAACGAGTAATCCTGCGTGGGACTACAAAGTCGTTTTCGTGGACCAGGCCTCGACCTCCCAAGCATATGGGGACGTCGCTCAAATCCACA
GTAGACTATCTATTAACTAATTGTGGATCGATGCTGACGCTACCCAGGTCTAGTTCGAGCCAATAATGTGCTACGTCGCGAAAGCCAAACAACGCTAACACCCTATCGTTCATTTGATACGTAACATCAATCACCTTGCAACTAGCTCGAACGGTATGGTTCAACGAGGGTGT
TAGGCAGCGAACAACCCAGAGCGTTTATCATACCATATGCTCAACATACTTCCAGAGAGTCTGTTGGTCTAACAAGGTATAGCGGCGAGCAACGGAAAAGATAGTGCGTAATAGTTCGAGGAGCACCGCATCCCCCTCGCCGTTTTCGGGCCGTTAATATGGGCCGTAAC
ACACCGGAGTTTCATAAAACGGCCTTTCCGTTGCTCGGGTGAGGGCATACGTGCTGCCTGGAGGGTATCCTACATGGTTAAATGCTGCGCCCCGTTAGGAAGCAGCAAATAAGAGTCTACCCGGTCTACCGGTACAAAGTTCGTGAGAGTTCTTCATGCTCGTCTCATAA
AGGACTGGCAAAGGCCTTTGTTGCGTCAATGAGGAAGCCAAGCGGCGAGGTCAAGGCCACGAGTACATGGGCCATTCTCCAGTATAGTGAAATAAGCCCACGCGGTAAACCAAATGTGGAGCACACTCCACGTTCCTGTTCGTGGGCCTATTGGGATATGTCTTTATCA
GACAACCTTATTTACGCGTATACATGCGGCAGTAGGCCTCGCGGTCCTAAGGGTCAGGTGTTACCACTTACCTCAATTTTCTCTGGAGTAGTACTTAGACTACCATCTTTGTTGAGAAATCATCTCACCTCTCTAGTCAGGGATAGCGTCCACTGTCTAATTATACAACTATTC
GATCAAGGTCGCAAGTGGATACTGAAAATCACTATGTAGAGCTTCGGAAAGCCTGAATTTTAAGTTATATCCTAAGAGCAGTGAACATTTCCAGCGATTTCCGTAGTTGCAATGTCAAACCCCGATCTAGGGTTAGAGTGCCTAGGTGCTATCCACGTGGTGTCCCTGGGA
GGCTACCCTTCGTGGACATAACTCGGCAGACACTAGTCCGATCGCCAGGACAAAAATCAGCGACGAACCGGTCGAATCTTAAGCGAAATAGACGAGAAGTTTAGGCATGTTAAAGCGGGTGCCCGGCACTGGCGCGAGTAACGTTCCCTCCTCCATATCTATACACCAA
CCCCGAACCAGGAAGTGGGACTGGAGAGGTTTGGGTATCTTCAAAATTGTGCATTACTGTTGGAAAGGTTGGATGCTTCGCGCAATATCGATACCGATCTACTTGCGGTGAGCGCAAATTCGCGTAGCAAATAGTGACCCATCGCGGCCATCAATGCTGACAGGGAGCA
CAACCAGATCACGATAACTTGTAAAGGGGGTATAGCAGCTCACCGGAGGGAAATACTGCAATCATGGAGTCGTAAAACAGGGCCATGTCAACATCGTACGCGTTCGTCATTCTAGGTATTTATTCAAGGGGATAGAAGCGTGGAGTTATTGAATCCAAACCCAAGCCGCTG
CAGCGAGGGCTACCACCCACGAGATCGGATGGTAGGGGGCGCCCGTGGGTAGGGGCCCTGGGTTATCACAAAAAGATCGCACTCTTCCTGGGCTTATGCGATACATCACCGGGAAATGATGCCACGTACCAGGATCGTTACGGGACCACGTCGAAGGCGCAGCAGC
GAAGTGTGCTCTAGTATGTTTGTGAGTAGCACGGGTCGACGTTTATGCGTTCTCTTTCCTCCTAGATGGCAGCCTCCCCAATGATGGGAAGGACTTTTATCGACATGTGAACCGGCCCTAACGAGACAGGTCTTGGTCTTGGTATCGAGCAAGTATAACCAATTTCTTCAC
TCGCAGAAAAATGTTGACGAGGAGCAATTCTCTATTAACGCACGGTCTGCTATTTCGGGCCGGAGTCAAAGCTCGTATATCGTGACCTCTTGGGGATCGTGTCCCCGCTTGTGAATGTAGGCTCGCGGCCAACCCTCGTTACTATACGACGGGAGTGGGGAGTACGACA
TCAATAAAATAAGCAAAACAGCTCAAGATGCGACCTAGCCAATGATTTTAGGCGATCATTGGATCCgcgccgcgggccgcTCAATCTCAATCgcggcccgcggcgcGGAT
>5mers_1_curlcake_2
ATCATAATTAATACGACTCACTATAGGGAGACAGAAGCGGATTATGGGGGCGCCATTATCGCGCGCGTGCTACCCTGAATGATACAGAGCGTGTGTCCACGTCGAGCTGTTGCGTAAGTCGTCAGCATACGTCCCTTGGACCGTGCATATTCCTCAGTCACTACTTTCTAG
ACTCAATATCTTTCGATGCTCCCGTCGATCAAGTAACTCGCTGTCTCGCTCTGTGAGAGCAATAGCGCCTCGAGCTAAATGCCGACGCAAGGGGAATCATAGGAAAAGCCAGCAAAGCCCCTAAAAAAACGAGCATGGCTGTCATCGCTTGGGAGATTCGGGACTTAACA
ACGTCAGGCTAAGGGTAAAGGTTGGCCGTGTTAACCAGTTGCTCGGTCATGTCTCCGCTACCTGTGATCATATCGAAAACATAATAGCTAACTTGCCCGCAGATGGGCCGTGTGAAGCTGGTAGCCGACCCCACGGCCTTACCGGACACGCCCGGACCCCCGACACTC
GTAGTAAAATTGATACATCGCGTTAATCGTGGGGCTGGCGCGTGCGACGGTTCTTGACATGTGTGGGTTATGTAAGTATAACAACCCTTGCGGCGTTGGCACGGCCCTAAGCTGAGAAAAAAGGTGCTCATAACTCAAACTTCGACGTTCCCCGTGGGTGGCGATAGGC
ATTGGTATGGTTGGTCTTCTGCAAGTCGCCCAGGATGTGATCAATGTAACGAAAAGGAATGATAGTAAAAAGATCCGGCTGTGGAAGTCGATGAATCAAAAATCTTGGACCCTATCTCACAACGAACGAGTAATCCTGCGTGGGACTACAAAGTCATTTTCGTGGACGAG
CCCTCGACCTCCCAAGCATATGGGGACGTCGCTCAAACCCACAGTAGACTATCAACTAACTATTGTGGCTCGAAGCTGACGCTACCCAAATCAGGTTCGAGCAAATAATGTGCTACGTCTCGAGAAACAACCCGAACACCCTGTCGTTCATTGGATACGTAACATCAATCA
CCTTGCAACTGCTCGCGGTATGGTTCAACGTGGTTGTTAGGCAGCGAACCACCCGGAGCGTTTATCACGCCATATCCTTAACAAGCCTCCAGAGAGTCTGTTGGTCTAACACTGTATAGCGGCGAGCGACGGAAAAGGTAGTGCGTAAAAGTGGATTAGCACCGCATCC
CCCTCTCCGTTTTCTGGCCGTTAATATGTGTCGGAACAGACCGGATTTCATCACACGGCCTTTCCGTGCTGGGGTGAGGGCATACGAGATGCCTGGAGGGTATACTACTTGGGTAAATGCTGCGACCCGTAGGAAGCGGAATAAAATGTCCCCGGTCTCATGTACAATGT
TCGCGCGAGTGCTCTATGCTCAGCTCATCCGTCACTGACTAAAGCCATTGCTGAGTCCATGAGGTAATAACGCGGCTCCGACAAGTCCCTAATAGTGTGCGCTTCTCCTGAATTGTCAAATGAGCCGACGATGTATTCCAAAGGCGGAGGCCACTCCACGTTCCTGTACG
TTGGGGGATTGGGATATGCATTTCTTAGACCTGCTATTATACGCGTATAAACGCCCGTCGAGGTCGCGTTACTAATGGTGAAGTGATACCACACACCTAACGTCCAGCAGTATTTAGATACAAGATATTTGAGAACTCATTAACTCCTCTGGTCAGGGATGCTTCCACTGGAC
TCTAAGACTGTTTGATCAAGGTGGCCTATAGTTACGAATAGCTCTGTAAAGAGCTTCGGATGGCCAGATTTATTAGGTACACCATAGGACCAGTGAACATATACCGCGACATCGGTAAGTGTAATGCCACGATCTAGGGTGTGAGCGCCTAGGTGCTTTCACCAGGTCTGG
GAGAGCCTGCCCTTCGTAGTCAGAACCCTCAGACTTAGTCCGATCGCCAAGTAAACATGACGTAGCCCCGCCGAAACGTTATATAGGCTAGAAGTATATGAAGGTTAAAGGGTTCCCGGCACTGCCGAGTCAATTCCTCCTATTTTTCAAGAGGCTGAAGACAGGAACCT
GGTTTGTAGATGGTCGACTAGCTTTTAAACTGAACTACCAGCTAGGAGGTTTAGCGTACGATAAATTTCGATTCCGGTTTCTCAAGTTGAGCTCCCCTGACAGTGGTGCCTCGGGACACAATACGGCAATCTGTGATTACAGGGTCCAACACAGCAACCAGACATGCTAGT
AAGATTAAATATATCAGAGCGGGGGGCACGCAGCCAGCCTTCCCCAGTAACGACGTTTCAGGCCCACTTCGCTTTAGTACGGAGTCGTCGTACTTCTGAAAATAGAGTAGGCGAATCGAAATCCCACCAAGACGTGCCCGGGGCTACAGCCCAGAAATGGGTGGGGCG
GCCCGACTTGAGGCACATGTTATCGCTATGAGTACCGAAGATGAGAGGATGACATACTTAATTGAGTGGGTTGACGGGCCATGGTAGGTCATAGACAAATTCTAATTGGATCCgcgccgcgggccgcTCCATCTCCTACCgcggcccgcggcgcGGAT
>5mers_1_curlcake_3
ATCATAATTAATACGACTCACTATAGGGAGACGCAGAGTTAGTGGCGCTGGCATGTGGTAACCAAAACGGTCGTGATGATGCGTTCTCTTCCGAATTTGTCCTCATACAGAGGGAGCTAACCTTATTTCCAGTCTTAAGCGAGACTCGCAAGAAGGCATCGTTGCACACTA
GATTGATTCAATGCAAAAGCAGACGAAAGGATAAGAGTGTCTTTCTATATTGCATCATGCCGTCTTCGAATGATCGGCTAATCTACACATAGCCGGCCGCGCCAGTTGTCTCTCTGCCAAACGATTATGTAGGGCTCTCGGCAGGATCACTTATCTGAGACGCGAAGTTCCA
TAATTTTAGGATTCGACGCAATTGGTACGCACGGGGTTTTGAATGTAACTGTCCGGACTGGTGGAGTGAGTTGCGTCAGTCGCACAAGGGCGCGGACCTTTTTGTTTCCTTTAACGGCGTCTACTATGGCGTGACCCCTTAGCCTAAACCATGCGCCCATGTCAGCGCTC
TTGTGTGGGCAACGGACGCCGCCTCTACGCCTTACAACAGTTCAGCCGTACATAACAGCTGGAACGGGTACCTATGTTTAAGTAGTAGCTATCCAGGACTTTCGCCTGTGCAAGCGCATTGAAACTTGTCGAACTTCAGATAACCCCGTGGCAGTGCCGGGAACTGGCG
ACTGCGCGCACTCAACATTGTTGTACTGCACGTACCAATGAACCGTGAAAGTACAGTCCTGGGTCACCCCAACGCTTGTAAGGAAACCGACCACGGATCGAGTTCTTGATAATACTGTGTTCTAGAGGTGTATCTTCAAAGAAAGACCATCCAATAGGGAAAGCTTGCTTA
TGACTCAGTACTCCGCGGGATCTGGACCCTACATGGATGTGACTTCCTAGCGCGTGTCCACCGGCGGGTTAGAGCATTAGAAAAAACTCGTGCGGGCGATTTGCAGTCACATCTGCTTGGTTATTCGGCGCAGTTTACTCACGGTTCTGCGGATAGTCGGTTGCCATCT
ATCGTCATCTCCAAGGCCGATTGCGGCATTCGTCCCATTTGGGCGTTGATGGAGACCCATCGCATGGCTGCAGCTTAAAAACAACGGAGGATGATGCCGTAGGCTGTAGAATGCAAGCGCGAGCGATCGTGATCGCTACCGATCCAGTCTTCCATAGTTGGATTCTGCTA
GTAGTAGCGTAGCGGATAATATGCGGCACGTGCAGCGTGGGCGTGAAAATCATCGAGGATGTGTGGATCTAGTTGGGACGTTGACTCCAGCTTATGAGAGGAATATTAGTAGCAATAAAGCGCGCGCGGGCAACCGGAACAAGGCCTGCCGTGGTCGTATCGAGTCCA
GGGGGTACGGACTGAGTGCGGTTAAGAACTGTCGTCTTAACCACGTATCTCCTCCTAGATCCTATTTTGTAAATTAGTGATTGCGCTACGGACCCTTACAACCTGAAAGAGGTGTTAGGGATATGAGATAATCGAACTGATCTCGGGCGTCCCGCTAGCCCGGTTTCGTAC
CGGGTTTGTTGACTTATGTCCATTGTACTTCAACGGTATCCCACTACGACGTAGCGACATCTGTTTGGTATGGCATCACTTGATCCAGCTGGTTCTGGGTATGCTCGACTCCGAAATGTTGACTGGGAATGAATGGTTTAGTAGGTTTGTCTGCGAACCCAGATGGCTTGC
AAGCACTGCGCAGCCGAAGACAATGTACCGTAGACAGACTGACGCACACCAAAGATTTTGCTCACTCGTGGCTTTGAATTCGCGAGGACCGGGTAACAGCGGGACGGAGGCGACGGGCCCTGTCGACGACGAGCTGCCGACTAAGGGCTCACTTTACTTTCAGGGA
CTCGATAATGGTTCCCTTAAGCGTCTAAGTAGCCCGTACAGTCTGCGCCCTTGCTCCACTTGGTTTCTGGATGCCCTTTACGCGGGACGTCCAGATGCCACTCGGTGACGAAGAGTTTGGCGAATGACATACGTTGCGTTTGATTAAGGCATCGCTCGCTTGTCTCAGTG
CTGAAGGACCTACCACCCTCAAGATCTCCTAATACACAAGACTCAGCGCAAAGGTCCGGTTGCTGCTCCCCTTTCACCCGGTCCCCTCGTGGCATAAGGTTGTCGACCGCCCTTGGTTTAATAATGGTCGCCACGTGAGCTTCTACCTTCCCCGAGAATGGGCTAACCC
TAGCTGATACATCGCTGGGAGCGGGCTGCCGAGAAACTAAATCAGGGTTCATGGTGTCCGATTTGAAATCCGTGCTCCCTTAGCCTGCGGGAACGACCCGTAATGGCCGTTCTGGAGTTAGTTAACCAGACCAGCCGTCGCACTAATTTGCTCGCCGTTGACATTATGC
AGAGAACAGAGGGCAGTACGCCAGGAATTGCGCCCGTCAACTTTTGAGCAATAATATGCCTACCTTATTATAGTCTAACAGAAGTACCTAGCGTGGTGGCAGACGGCCGATAGGCCTGTAAAGGCCATGGAGGCAGGTGTACCATCTACGGGAAGTATACGTTCGCGGAG
AATACAATCTTAAATTTTCGTGGGGAATATCACATGGAACGTGTCACCATAAGCCATGGACTGTCCATGTCTATTAGGATCCgcgccgcgggccgcAACCAAAACCAAgcggcccgcggcgcGGAT
>5mers_1_curlcake_4
ATCATAATTAATACGACTCACTATAGGGAGACCGAGACTCTCGCCCCTCGAAGCGATCCACCCCGCATCCTAAACCCAGTCAAACTCCTGGACCGCGCTCCCTTGAGACTAGGCGATGCTCCCTCGCCGTGGTGTTTTCTCTAGCGGTATAGCGCTTCTGTCTTCCTATG
GGCCGACTAGAATGACTGTCATCACTGGCATGGCATCAAGGGCCAACCTGCGTGGTCTGATTAGCTTGGTTAGGGGCTTGCCGATGTATTTAGCGCGCGTTTCGAAACAGCGGTTTCAACATTGGCCATAGCTACCTACGAATCCCTACGATTGATACACGGTCGGGACA
AATGATCCGACCAAGGTGCCTCACTGGTGGGAAGTCTAGATTCATCCGTGGGACATCGGACAGTCCTCCGTTATGCGAACCTATGACTCCCACCCCAAAGGGCTAAACAAAGACATTCCCGTACGCGAGTAGAGGGAGCCTTAACCGCAGTGGGATGCTAGGATCGGC
TAAGGGTTTCTCCTAATAGGCGGGAGTTTCTTTCAATGAATACATTGACTCCTCACCTGCGTAAGTTGCGAACCTGACGCTTGCGTGATAAATCCCAGGTTCGAACGCTAGGAGACGAGGCTGCTGCAGGCGGCTGCAGTTGGAAACCATGGATAAATTGCGCCACCGG
GCTACTTAATGCCTGCGCTCCAGCCCTTTGAGTTTTTGCGCGCGATCCAGTAGAATCCTCCCCGTCGGAATTAGGCCTGTGGCGCGCGGTGCACCGAGAAATTAAACTGTGGCTCACAACGGCTAGGTTTTTCTGGAGCATAGCAGATTGCCTCATGTCCATCGGGCAT
ACTGGTGATTGTCCCAGCGTCCTCAATTTTTTTAACGCTAAGCGAGTAATGCCATCGTGGCGAGTGAATGTCGGAGGTGGAACACTAACTAGGTTAGTGGCCGGCTCCCGGCCCAATTCGGACATGCTGTAACGTTAAGATTAGTAAGACAGCTGTCGTTCTAGCACATC
GGCAAGGTTATAGTTTTTCGGTTGAATGCTCACGTCCCTAAGCGGCAGGAATAGTATAGCTTTGGGCTCTTCATGCGGAGATCACCTCATTCTGCTTCTAACTCAGGGGCCTACCACGGTACACCGCCACATGTCGAGACTTAGTCTTTTTGACGGCGGAACATCTCTCAG
GCGTTCCAATCGGGCGATTAGAGCACCGAGCGTTACTTAGGATGGGTAACATTTGTATGACTAAGATCACTTGAAATAAATACAAGCGGGTACGTTCGAGACGAAACGTGTACATCAGGTCACGTCGTGCCGAAGCGTACCCTTCGTCCTGATGTCGCAACACTACAAGG
TCTGCCGGAGCCTAAGTTAGTGCTAGACAAGCCATGCGCCGATAACTAAAGCAACGGTCGGTTTCCTCCAGTCACGTTCTACGTTACTCAGGATTTAGCATAGCCTAGAGACAGCAGCGAGTAGTATAATACAGCGTCGGCCGACCCCAGCTAAGGACCGTCTTCTTCGA
TATATGGCGCCCCAAGCTTCTGAACAGAGGGGATCTCGCTCGCCCCTATATGTTCTGGTACTTACTAAACGAAAAGACAGGGTTTTTGGAGTCAAGCCTAAACTGGTGCAATGACCCCTAGCTACTGAGGCCCTGACATTGGTTTGTAGAACTGCTCGGGTGTACCGTCC
ACAGGTACAGCAAATCACGTCACTTGAGAGCGGTCCCGTGCTGTGTTACGGCACAATAGGCACCTTTCGTTCAAAGCATTGAGAGACGTTCCCCGAGGCGACCGGGTCACGCAGAAGAGGTCGGTATTCGCAGCCAAAGCATAACCCGGTCCTCCTCAGAACAGCAG
GACTCAAGCATAGGGACAGTGTTAGCCAGCCCAAGAAGGCAGCGGCGATTGCCGACCATCTTCCCTATAATGACAAGACCTTTCAGAGTGGACGTCTACTATGTCTTGGACTGCACGGGTGTCCTACGCATCGCGTGGGTCCAAGTCCCATTAGCAACTTCTCTCCCGT
CGGTACAATGTTACAGTTAAGCAGCGCCTCTCAATCGAGAGGCTGCACGGGAGCTCGAGAGGCGGTATCGAGGGGGTAGGCCGTAAAATCGCTGACTCAAACGACTATAGGAGGCGTGTTGCATGTTTAATTAAAGCCATCGTCCAATTTTCATAGCTTAAGCTAACCCG
GCCACCTTGAAGAAAAACAATTCCTTACGAATGTGGCACACTGTTCCATAGAGACAGGGGGGAGTTCCCTCCATAAATCACTGTTCGTATCCGAGGTCACTCGAATGGAGTGTACACTAAGCTTACGAGGGTGTACTGCACGCAATAAGTAACATAGCCTCCAATGTTGGA
TTGTGTTCTACCATGGTGTGGTATCAGTCGCTGCGCTGAGTGAGTCAAGAAATCGGGAGGTAAGTTTAGCGAAGTAAGTATTGCGCGTAGTCGACCCGGATGCCCACAATCATCCATCGACGACCCCTCTTTTACTTTATATCCTTGCCGTAGGCATGGTGAGGCCATTAG
CGGGTAATCACCCTCTTAGTCCATCTCGCGCCATCGAATTTCATTATACTCCGCCTTGGAGACTTACATTAGCAGTCTACGCTCTCGTTGCGCGGTCCACTCACTACTATCTGACCACTCGCAAACATCGAACCTGCGGGTACACTGGTCAGCCAAACAGGATGCCGGAC
AGACAACCGAACAAGAGCATGTAGGGATCCgcgccgcgggccgcTCTTCCTCTTCCgcggcccgcggcgcGGAT
>5mers_2_curlcake_1
atcataattaatacgactcactatagggagaAAAAGGATCGTACGATAAATAGTTCGTTCGTCTTGAGTCCTCCGTGGAATGAATAGATGTGACTCCCATCATAACTAACGAGCCTCAACGGGGATTTTCCTTCCCCGGCGCAAACTTCGCTCAGTCAGTGCCGCCTAGTTTTGCCTTAGC
TAGCGTTGGTATTGACAGACCGGACGGACCTGCGAGAATTAGAGCATTATTTATGTGGGAACGGATTAATTATTAATTATAAAACAGGTTTAGGTTACCAACCAGGACTCTATAGTATTACAAGCAAGCAAATTCCGAGTTCTCCCCAAACAAATCGCCTCGCATCTTACAGAA
TGAATGAGGAAAGCACTGTGAAATCGATCGCTCCCCTTTAAACTTATACGTACGTGTCTATAAGAGTGGTCAGAGAGAGAGTCAACCTCCACCGTCGGAAGGAAGCACTGGTCGAGACGTGATCTGTCTGAACTCCCTTCACAAAATACGCACGCTAGGGTTTGCTACGA
GTATGGATGGGGGCCGTCCTTTAAAAGCACCCACCAAGGCGAGAGATGAAGGTAGATAGAGCACCTTTTCGTCACCCGTCGGTCGGGCTCGGTCGAGGATTTGTGTGTGTCCACATGCGGAAGATGATAGCTAGCAGAGAAGACCCTCAACGTTGTTTGTCTTAGCATA
TGATTCCGCACGCACGCGCTTTGTGGGCTCCCGAAAGAAAGGCGGCGAAACTTGCACTATCGATCGTTACTCGCGCTCTATTTAAACAAACGCACTTACTGCTACTCGGCTGGCTGTTGTCTGGGCCGATGGGGTATGTATCCCTCGCTCGTTAACCAATTTATTTAAGCT
TAAAGGAGTTTTGTAAGTAAAACGAGGGGAGAAGTACCGAGCGAGCGCTCCTTACGTTTCTGCAATCAATGCATTCAAAACTTGGGCTTAGTTAGATAAGCCCGCCGAACGGTCCATCCAGAGCTCGCAGTTAGACTATTGATTGAATCATTGCGATCAAACTAAATAAACC
ATCAAGAACCATACCCCCGCCCGACTTTTTGGTTGTGCGAGCGTGCGTCACAATGTAAGTCTTAGTCAATCAACTATATAGCTATTCAACGCCGGCCGTTTCGAAGCGAGTGTTTGCGTGCGTGGGAGTCGGTGTAAATGTATTTATTTTATGGGAAGTAGTATGTGCCTGC
CGAAGTACTTAATTAAGGGGGTGGGTGTCCTAACTAAGTCATTTACATACATCCAATTTTCAGTAGGATAACTAACTCCACCTTCCAGCGGTCATTGATTGGACTTTTAATAATGCAGTGGCTGGCAGGCCTTGGAACGGAATTTCTTTCTGCTAGATACTTGTTGCGAATAAA
TAGTTCGGAATAGACGGCAGGAAGGAAATTCAGAATTCCCGTGACTTACTTAACCGGACGCCGAAGTGAATTTATTTCCTCGTTATGCGTTTGCGATTGATTCGTATATGAACTGCCGTTTTGTTTGTAGTAACTGGATGCGAGAGTGAGTGACAGGACCCCGCCTACCTG
CGTGCGTCAACTGAAAGCCATGCGGAGGGAGGCCGACTGCAGACTAAAAATTAATTCTCACTCGTCACGCTAATTATTTATTGATAACTAGCAATCAGGTAGGTAGCACGTGGGTAGGATACAGCCAGCCAAGTCTAGTTTAGGGTATGTCAGTCATACAACACAGACAATC
AGTCTGTCTGGCTCTTTGGCGATGCATCCCTCCCTGATCCATCTGTTCCAGGTAAGTAAATTCTTTGTCACTTCCAACAAACAATCGATAAACACCGGTGATTAGTTAGGTATTGAATGGTGCGGCGCTTGCTTCACTGACGTGCTGGAATCCTCCCTCGCTTAGTCGGCTT
GTTGGATAGATACAGGCAATGGCGGACGAAATGAATGTATCACAGCAGGGAGGTTTTCTTTCGTTAACACGATGTATACAGTAAGTAGGCGCCCTGTTTTACAATATCTATCGCCCGACGAGCTGGTCTCTGGCTGGAAAGATATTGCATTCGCTGAGTGAGAGGGTCGTT
GCATAGAACGCTCGCTTTCGACGATCACGGGCCTAATTAATTGGGCATGAAGACAAGATCTATCTAACGACGTACTAGCTCGGCCCCCCCCATGTCCCGCAAGCTTCGCGCGCGCGGAAAAGTCCAACAAAGTAACAAACAACTACGTTCCAAGCATTAAGGAAGGGTT
GGTGAACATAGGCGGCAAGCAACGTTCTTTAGTTCTAAGATTGATTGTCCGTGCTCAGCAGTCTTCTATCTACGCTTATTGGACCTCTTATCGATCGTTAACGAGGACTGTGGATCGTTCGTATGATTGCTTTCCCCGCCAGCCAGCTCAGGCGTTGATACATATGTGTGTG
AGATATGCGGTTCCGAAAAGCAAGCAATTATGCATAAAAGATACATACATATCTCTGTCAAATAGTTAACAAACAGTAGACCCTACACTGGTGGATGGATCggatccgcgccgcgggccgctcaatctcaatcgcggcccgcggcgcggat
>5mers_2_curlcake_2
atcataattaatacgactcactatagggagaCTATAAAGAGTACGACTAGGTAGGGAGTCCGGCACGAACACTACATACACATTTGAACTCCGGGCCATCGATCGGCCGGTCCCACCCCAACCCGTCCGTCTTGTCTAAAAAACCTATGAGTGAGCGGCTAAAGAAGCATGAGTATGTAT
GACTACTTACGGCCGCGTATGTATGCCTACATCGGGTCCAGCGACAGACAGCCGATTGCCCTTTCCTACTGACTGGCCTGGGAACAAGTTGAAAGGAAGGCATTTCTTCTGGTTCTGTACCTACCGATTGTGTCGCATAAGCTTTGTTTGTGAGTCAAAGGGATAGTATAA
ATAAAGCCACTTCTGATGCTAGCCCGCCCTCATCGCCTCGCCGCGCTATGTATGAGGGTGTATTTTCAGAGGCATGCATGCTGAATATTAACAGAACACACACAATGCCCCTTGACCTCCAGACGGACGGAGTGGCGCGTAGACTTGACGGACGGGTGATCTAATGTGCT
ACAGGCAGGCGCACACTGACCGCTCAATGCATGCGTCGCGCGACCAGCCAACTTTCTTTCACGTAGCTGTCTTAGAATCAATCACGCGTTTTAGGTGCCTCGTCAGTCACGCCCATTTGGCCTTCCCTGTCTGTAGTGATTCAGCACACCATTTGTTTGGAGTTAGGCG
CAGTATTCTTGCTTGCCCTATCTCCCAATCACTTCGTTCGTGCGCCACTGCCTAACATGCTTGCTTGTAGACTCGAAGCCTTTGCTTGCTCTCCTCTGTCAGGCCATTCTCTCTCGGGGCAACTAGTCGTGCGTGTGTGTACGTCAGAAATCGTTCTTTTTTTTTTAATCTAT
TGACCATTTTGATTGATTTAGCAATCCATAATTAAGGGAGGGCGTACTCGGTAATATATCATCCATCAGTGTTGAGCTACTTCCAACTAACTGTACTGCGTTGCACCTCCATCCATCCGCTGCCAGGCCCCACTTGAATGAACCCGGCCCGAACTCTGCATGTATGTTAAAAT
AATTGCGCGCCGTACGTACATTGAAACGCATGACAGCAAACAAAGGTTGACCCCCTAACAAGCTTGCTTGAGATGCCGCCTCTATGACAAACAACGGCAAAGACCGAGTGCGGCCGGCCTTATCGTGACAAGATTTCCAACCAAGTTGTTCGTGACATCCCATGGATGG
ACTGGGGGTGTTCCGCTTATATATACACCCGATGTAACCGCACACTCACTATTATTCGATTCGGCGGTGAATGAAGAAATCGCGAACCGATTACGCACGCTATTTTGAGCTGTGCTAGTGCGTGCTCCTACTCCGACTGTTTCTAACTAAGATGGAATTGCACGCACCGCG
CGCGCCACATGAATGCCAGAGGTTCGTTCTAGTCCATCGAAAACCGGGAGGGAGCTAAGTGAGCCGGTCGAGATCGATCCAAATGGAGCATGGTTACGGGCGGGAACTAAGAGGGTAAGACATAAATAAGGCCTGGACTCAGTTTGTGTTTGTTGATATTCGGTCCCTT
GCGCAGGCAGATTACCACCGCGACTTCTGGCTGGCGCCCTGAGTGAGGACACTAACTAACTTACTACCGCTAAACCCGGCTGGCTGCCTTGATTATTACCCTCCTGCCTGCGCATGCCAACGTCCTCTGAGTGAGCAGGCCAAGGCATTCTAGTGGGTGGATGTGGCG
AAATGGTTAAATTAATTTACATCGGTTCGCATCCAGGTAGGTGGACCACGATAGTCAGAGATTGATTGGGGCGTCTGCGACACAGATCGATCATTCCGTGAACCTTACGTATCTATCTATTATCTCCCTCCGCAGAGTGAGTAATTCCGTGGGGCGATCTGACTGACGCTTG
CTAGCCTCCGTATTCGTTCGCCCAGAACACATCCCAATCCGTCCGTATCGATTCTCAGAGTTAGTGAGTGCCGATGGATGACTTAATGAGGGAGGAATCATCAAGTCTGCCAGGGAGGGGACGATAGTACTACAGGGCTTGCTTCTCTTTAAGACTTCTTGTTGCTTGCTC
GACGAATGTTCCAAGTAATTAATCTCAAGCTTTGAACGAATGATTGATGATCACATGGCCCATTAAACCAACCAGTGTATAATTGTAGGACTGACTGACGCGTGTCCGACTCTTTTAAGTAAGCGCGGGGGATCTTTCTACACACACAAGTGTATTATGTTTAAGCCGTCCGT
GGGATATTCCTTATAGATCGATCGAGGATACTAGAGCCACTTTAATTAATACGTTGTTATGGAGGTACTTACTTATTTACAATGGTCAAAGTCGCTCGCTTTGAACATGGGACCACATACTTACTTTTATTTCATCTTCCTTAGGTAGGCAGCATAGCTAAAGGAGGCTCGCTC
TCTTCGTTCACAATCAAGTTGGTTGCACCAAAACAACCTAATACTCACTCTTCAAGTCCTAATCTAAGTACGTACCTAACATCTTGCACACAATTGATTGAGACGAAGACTGTCAATTAGATAGAAGTTCAGTCCGCGTAGCAAGTggatccgcgccgcgggccgctcaatctcaatcgcggc
ccgcggcgcggat
>5mers_2_curlcake_3
atcataattaatacgactcactatagggagaAAGTTGACTCTTCGAAGGAACTGGCTGGCATAGTAAGGCTAGAATTCTACCTACGGGCTAAGGGGATCGAATCTTTCTTATACCTAGGCACGAAAAACAGACAGCAAGACAAGGCGGGATTGACAGACAGCGTGTACACGGGACGAGG
TGGGTGCGTCGCAGTCGTAAGGGCTACCTACGATCTTCCCAACAGACTCATTCATTCGTGACGTCTCATCATAACTAACTACTCCCGCGATAGACATGGCACGCACCCCTCGGATTACCGTTCAACTAACTTTGAGGGGTACGCTATCAGTCAGTCCCCGAGCATCCTTCC
TCCCCGCCCGCCCGACCTGCAACCCGCCAGTCAGTTGGGGGATGGCCGGTATGTAAGTAAACGATCGCGAGGGACAGCAACCAACGGGGCGTAGGTCGGGACACGAACGACAACGTTAGATTTGGTTTCGATCGACCCTGTCCCCACACTCCACGTACGTAAACGC
CAAATTTCGATTAGATAGAAGACACCAACCCGCGGTTCGATCGAGCGAGAAGGAAAGCGTTTTGGTTGGCTAATACATGAAGTTTTGAGTGAGTCGCGCTGGGCACTATGGTACATACATGCTTAATGCCCACCTCTGCGTGCGAAGCTGCTCTCCGGACATTAATTAAAT
TTGGCGCACTATCATGAGCGAGCGAACATCTGGCTTGGACTTTTTTTTTCGAACTTGTATTATGAGTAGGTAGCACTCACTGACCGCTCGCTGGCTGGGTTGATGTCGTAAGATCGCTGGCTGTGAGAAAATTGCGCCTACTGGCTGGCCGGAGTTATACCCTATACAAAC
AAGAAACGTTGACGATAAATGCTTGCTGGTATCGGGTCAATGTTTAGCTAGCGCCGTACCTCACCCAGCACTCACTCGCCCGTCCCGATAGACAGAAGGAAGTAACACCTTAAGTAATTCATCCATCGTACGTGGCTTTACCAGTTTGTTTGCTGTCTCCGTCCCTAGCTC
ATTCATAACATTACAACATACCCGGTGGGTGCAACGCGACCCAATAACAACCAACCTACTTGTCGGAGGGAAGACTAACTACTGAGTGTGCCACACCAAAGAAAGATATGGAGATCGTTTGCCCTGTCTGTTACGGACTAGAGATTTGGTGGGTGGCCGGGTTATTAGGT
GTATGTTTGTTGCCAAACGAATCAGGTTGTGTGTGTTAACCGCTATATCCTTGTGAGTGAGTGTGCGGATTTAGCCTTGTAAGTAATAGGTCATAATCCGAAGAGCTAGCTAATATATGCGCCACCGTTTGCTTGCGTAAGTTCGGAGCCTATAGGGAGGGAAAAAGCTACC
CACGCTTCGATCGAGTGGGTCTATCCGTTTTGTGCGTGCAGCGGCTGAGCTATGGATAGCTAGCCCATTAACGTCGTAATCGCTAGCTAAGTACTGAAAGCGCAAGGTACGTACCGGGACCATGGTTGTTTCATGCATGTCTCAATACAATACTAGCGACCGACATTCGGT
CCTTACCACGTAACTAACGTGCGCGGGGTTTAGGATGCGTGCGGGCGGACTAACGGGGGTTAATTAATAACGGCCCTGAGCGTGTTTCTTTCTTGCTTTGATGGCTGCTTCCAGCCAGTCCTCATGAGAATCTGAAGCAAGCTGTGCCGTGAGAAGTCTACCAACCAGC
CGGCGAATTAAATATAGCTAGCTGTACAGTGATTAATCTCTAAGTAAGAATCCAACCGGTAAAGAGCCCGCCCTTGTAGTGCTGATCCCTCTACCTACCAGTTACAAAATCTGCCGCACGCACGTTTCGCGCCTCCACTCAATAAATATTGTTAACTTATCAAAGTGGTTGGT
AAAGAGCAGGTGGACACGTGGGTGGCCGTCCGGTGCAAATCAAACTAACTTGCTGGGAGAGGTCTTTTTCTTTCTATACGACCCTATGCTCGGCAGGCAGATTCTTCACTCCTTGCAGTTTGTTTCCGGCAAGGTGCAACAATACGTACGCGGTATTCATCGATGATGGC
AGGCACAGATCATGAAAGCGCTGATGGATGCGGATGTACTATAGACTGCTGGCTGAGAGACGTATATGTCCATTCGTTCGGCCATATGCTTAGGAAGGACAGACAAACAGGCGAGTTCGACGGTAGGTAGGTAACCGTATGGGCTTCGCGAGCGAATCATAGCAGTTACA
TCGGCTGGCTACCTTTTGGAGTACGAATCGTTCGTCTGCCCCCGGTCAATTGGAGAGAGACCTCATCTTAGCTAGGTTCAGTCAGCCTTAAATGTCTCGGGTCATGCATGCACCTGAAACCACGGTCGAGGGAGGCTCCTAACGAAAGTGTCCAGACAGATGGCTGGAT
ACTTCATGCAGGCAGGACGGTTCCCGGCGAACTCACGCACGGAGACACATTGTTCGCAACTGACTGCCGTGTGGTGCGATTAGTCGTTCGTACCATCAGGAAGTGACATGCTTGCTCACATAGGCTGGTCTTATTGAggatccgcgccgcgggccgctcaatctcaatcgcggcccgcggc
gcggat
>5mers_2_curlcake_4

atcataattaatacgactcactatagggagaTTGAAACCAATTCAGCGATCCTCCATCCAAATGTTCTTTCACTATCAAAAAAAAGGGGGACGTGAATTGTTGCGGGCGGTGCAAACTAGTCGTCCAGACCGACCTGCTTGGCGAGCCCAATTGGCTGGCTATGATGGTTGTACCCCCA
CACACACCCAGCGTCGAAGATATAATGAATGAATGACTAAAGTTAGCGTGATTGATGACCGACTTACTTTCCATCTGTCTGATACGACTGCCAGTAGAATAGATAGGATAGGGCCATTAGATGTCGATCGACACGCACGATAACCCCTGGAAGGAAATCAGAATCCGGAGCT
CAATTAATTCCGTCTTTGCAAATCGAAATGAATGGACCGTTCTACTGAGAAAGAAAGAAACAGATGTGATTCACCCCTTTCTTTATGCCCGCCGACCGTGTCCCACCCAGGGTGGTAATTAGACAGTTGGTTGGTGGTTATCTCTCATGTTAGCTAGCTGCAGCCGGGCCA
AAGTGATAAATAAGTGAAAATACGCAGGGTCTGTCTGTTAGTAGCGCAATAACCTATTCATTCATGGTCTAGGACGCCTCTCCGTCCGGGAGTTCTTCAAGGTTTTTAGTTAGTTTGGCATGCTACACGACGTACGTAGGCCTCATGTTCATACCTGTGTGTGAGAGATTGG
TAGTATACTAATTAATAAACAGACCACGCTCACCTAGCTAGCATGTCGGTGAAGGATTTTTATTTACGTCAGGGGGAGGCTTCACGCACGCCTAACGACGTCTGTTTGCTCTCTCTGGCGTCGGGCAGCGGATTGGCTGGCATCGCTGAATATTTCAATCTCTCTCGTAGC
CTAGTAGAAACGGTTCGTTCCCAGGAGTAGAGACACCAAACAAACAACAGGTAGGTGTAGTGCCCCCCCGGAGAGGCTGTATATTCTGTATGTAGTGCTATTTGTAATCGACTACCTACGTCAATCTGGGGCGGATGAAGGAAGCCCTCTATCTAATAAGTAAATAAATTT
CATCTGCTGCATACGCAATCAATGACGCGTGCACGGCAGCTAGATAGAGGAGGTGTTATTTGAGGGAAAGAAATCCGCTCAGAACCAGTTCAACCAACATCTACATATTTCTAACGCGTGCGTACGGCAAAACGTCCATAAAAGAAAGTTGCCTCGTTCATCAATAATCAAT
CCGCACAGTAATGGAGAGCAGCCAGCTTTCCTATTCTCAACCCAGCTAGCTCTGATATTACGGCTGACTCAGTCAGTAACTCAGGTCAGCTCGATACATACGACCGTTGGGTAGGAATGAATGAATCGGTATAGCGCTGCGCGGGTCGGTCCGAGCCGGCTGAAGCAGA
CGAACGATCTTGGGCCCAGATAGATGCATGCAAACACCTAGATCACCGATCCCTCCCTATGCCAGAATGATAAGCCACCCACAGCCCGTACAGGAGGACGCGAGCGATCACTCGATGTCGACAAATGGATGGTGGTCAGTCGTGAACCCGATGGATGAGCAAGAGTATC
GACGCCACACACAAAAGTCAGCCCGGTATAAGAAAGAACGAGTTTGATTCCTGCAACAAACAAAGCTTAAGGCCTGTCATAGGAAGGAATATAACCCCAACCGCATCTGTCTGGGTGCGCAGTCACAGTCGATTGATTAGACGTTCCCGACACAGTATATATAGAAGGCTC
CACGGTGGTACGCACGCAAGAGACTGTCGTGGCACAGGCAGGTCTCACACGTCCGACAGTACGTACGGTGAGGGCATACTATGAACATACATATGGGATTTACCTTTACTCATTCATGGCTCCTAGGGATCGGTCACGCACGGCGAGAACACTAGAGTCTCCATCCACG
CCCCCAGTCCTGAGGAAGCAAGCCAATTTAGTCATTCAAAATAAATAATCCATTACTCCCGTAGTCTAACTAATGCATCGGTCCGGAATTTGTTTGTCATACCAGACGCTAAAACTCACTCACCCTAAGCCGGACCCATAGCGAGCGGGTCCTAGAGTCAGATCATTTATTTA
TAAAGTGGAAGGCCCAATATATATATTGGTACACCATTGCCCCTTCCTTTCCGAGGCAGCTGATCTTAACTAACCACTGATCAAGGAGCTTCGGGCGGGGGACGCGTTAGGGCAAGGAGCGAGCATACCATGATGCCAAGTTATATATATCCACTTTTTCGCCGCGTATCTA
TCTACGAGATGGCACACATACCGACCGAAAATAGGCCAATACGGTACGTACGCCTGCCGTAAATACCCGTAAGTAAGCGCACTCGAATTAGCCTGTGTGTGTGTTCATATAGGAGGGTCTAGGTAGGGCACCAAAGGTGATTGTTTATTTATCAGGGTAGTACAAAAAGAAC
AAACACGACATTTCCCGTCACAAATGAATGCTCAAAAACCCAGGAATATGTATGTCACGTTACCCACGAAGGTACCTACCTCAAGCGACCAAGCGTTATACATACTAAACTAAAATCCTGGAATGTggatccgcgccgcgggccgctcaatctcaatcgcggcccgcggcgcggat
>5mers_2_curlcake_5
atcataattaatacgactcactatagggagaATGTAGTTACTGGCACGAGGCTTGCTTGCGGTTCAAGAGCCCCCGTTGGGTGGGTGCAGTGCTTTACTTTCTCTATCTAGGCGTCTGTAACCATGACCTTCCTTCCTTACGGTACTCATTTTTATTTATGCGCCCCTGGTTATCTCTATAT
ATAGAGCCGGGGCCGAACGTAAGGAAGGGAACTGCATCAGCGACTACAAACAACCTTGGAGAGGTCCGCATTGCTTGCAGCGGGACTCTGATCGATGGGTGGGCCCAAGAAACAGCTGCGTGCTTGCTTACCAAGGTCGTCCCATCCTGCCTGTGGTGTGTTTCAGT
TTCTCCTTCCTGGCATCACCTTTATAATCTCGCTCGGCCCGGCTCGCAAGCTCTCTATCTAGAACCTTCACTATCGGAAGCAAGCAGCGCGGCCACGTACCTCTTGCTTGCCGCCATGTTAAGAGAAACCTGCCTGGAGCTCCGTTCGCAATGACTCACTCTGACGGCA
CTTCAAGCAGGCAGGCAAACTGGTTCCTGAAAAATAGCTAGCCATATCCCCCATGTTACTGTTTGTTGGCCATCCCACAAACGCGACGGACGGGTGTGGCGGCGGTTTACGAGCGAGCGATTTTCGGCGGCAGAATGCATGCCACCGAACGCAGAGTCTCTCCCTCCT
GCGATAATGTAACTCCATATATATACCTAGCTTCGTAAGTGCATTCATTGCCTGACACGGATGAGTATCGATCGGAACCTACAGGAGCGCTTGGTTGGTATGGCGCGAACCGGGAAGGACGGACAAGCGGACAACATCACTGGTGGGTGTTGGGGGCCATAGTTGAGCG
GGCGGCTTAGCCTGTAGATTGTCTAGCTAGCTGGTCCGCGTCCGATCTCAGTCAGGATGCACACGGACGACATTACTTACTCGACCGGAGACCCTTCTGCGTGCGTTCTCGAATAAGCGACCTGAATGAAGGTCGGACATGGGTTGATACTTACTGTTCCACCTCTCGT
CGCCATGCATGGGACCACTCTTCAGCCAGCACGCACAAGGTTAGTCATTTCACCGCACGCATCCGAGGTACACTGTCGCATTCATTGTCTGGACCAGTACTACGGAAACCAACGTGAATATGGGAAAAAATTGAAATTGTCCATGTGTGACCCTGGATGGAAACCGGTAA
TGGATCAGATTCATTCGGTCCCGCTGCTGTACCCCCCCCCGGATGTACACTTGATCTTGGTTGGTCTGTGCCTTCCGCTCTTAATGAATGGTGAATAGTGGAGTGGGGTGGGTGGAACTGAGGCTCCGGCGTAGAGAGAGGTCTCGAGTCCCTATGGGTAGGTAGCATA
GGGCATTGGGGCGCGGTCGGTTACCCTTGAGGAGCAACTCTATCTACCGTAGCCGCTGCAACTCTCGCTCGAGACCTGATATATCCCCTCGCTCGCAATCTAGGGGTTTATCAACGTACGTGGCTTGGACATTACACTAGTGAGTGTACAGTGGCCCGTTAGTGACGGA
CGCAGGCTCTCGGGAACGCATGTATGTGGTCCAGGGCAATGATCGTGCGTGTCAGCTTCGTGCTTGGGAGGTAGGTGGTAAAGGGGTAGAAGAAACCGCAAGACCTGTCGATGACCATAAGCAAGCCGCGCTAGGTCGCCAGCCAGGCAGGGATTCCTTCTGACAG
CTACCGACCGAGTAGCCCTACCGAGACGGCCAGGCCTTCTCGAGCGGGGTGACGGGCGGGCTAGATAGCGGGGAGAACGGACGGAACCACCAGACGGTTATCTAGTTAGTTGCCCCTCGTCTACGTGCACGCACGTCATTGACTGCTATCCGGGTTGGTTGGCTATC
CTGGGCGTGTGAGTGAGTTCAGCACCCACATTCCCCATACATATCACAGTGTAGGCAGAGGCAATGAAGTCTCACCGTGGTCCTACGCCCGCCGAGTTGCTCCTCACCCACTCGCTCGTGTCGCGGTTTATCCTCAGGGAGGGGCAGGTTGTGGGCGTAGCGTTCGT
TGTATCGCCCAGTGTGCCTGACTGGAATCGTGACCGGGTAACTGTTCGTTCGGTGATGCAGTGCCTGGGTTTGTTTATGTGTGGGTCGCGGGCGTCTGAAAAGACACGGTCTTATAAACTACGTTACCCCTCCTCTTGACTTGAACAAGTTTGTAACGTAAGGGCTGTAA
GCACCACTTCTGTGTCACGGAGCCGTTGGGAGCCTGCACCATTACTAACAGCTCCACCGTCGACGTATCCATTGCATTGCTACCGCAGCTCGTCTTGTTTTAAACAACGCGTCGGGCTACCGCATTGTATTGCAGATCCTGCTGGCCTAGCGAAGGACGGCATACCTCT
CGAGCGTCGTAATAAACGCAGCGTGCCCTCAGCGCGAATAACTACGTGCGATGTCTGTTGTATGTGCATTAAAAGTAGGGACGCCTGCTTTTAGATTTGCCCCCCATGCGGATTTCGCCGCCCCTACACCTTGTGCACCAATCGACGAACTTTTTGTAggatccgcgccgcgggc
cgctcaatctcaatcgcggcccgcggcgcggat
